# Grape-seed proanthocyanidin extract (GSPE) modulates diurnal oscillations of key hepatic metabolic genes and metabolites alleviating hepatic lipid deposition in cafeteria-fed obese rats in a time-of-day-dependent manner

**DOI:** 10.1101/2022.07.20.500817

**Authors:** Romina M Rodríguez, Leonardo Vinícius Monteiro de Assis, Marina Colom-Pellicer, Sergio Quesada-Vázquez, Álvaro Cruz-Carrión, Xavier Escoté, Henrik Oster, Gerard Aragonès, Miquel Mulero

**Affiliations:** Nutrigenomics Research Group, Department of Biochemistry and Biotechnology, Campus Sescelades, Universitat Rovira i Virgili (URV), 43007 Tarragona, Spain; Institute of Neurobiology, Center of Brain, Behavior and Metabolism, University of Lübeck, Marie Curie Street, 23562 Lübeck, Germany; Eurecat, Technology Centre of Catalunya, Nutrition and Health Unit, 43204 Reus, Spain; United States Department of Agriculture and The Agricultural Research Service (USDA-ARS), Arkansas Children’s Nutrition Center and Department of Pediatrics, University of Arkansas for Medical Science, AR 72202, Little Rock, USA

**Keywords:** circadian rhythms, liver metabolism, proanthocyanidins, NAFLD

## Abstract

Metabolic syndrome (MS) and its related diseases, including obesity and non-alcoholic fatty liver disease (NAFLD), have become a public health issue due to their increasing prevalence. Polyphenols, such as grape seed proanthocyanidin extract (GSPE), are bioactive compounds present in fruits and vegetables that show promise for MS treatment. We have previously demonstrated that the efficacy of this phenolic extract in the modulation of liver circadian clocks was affected by the time of the day at which it was ingested. Thus, we wondered if the beneficial effects of GSPE consumption in NAFLD could be mediated by diurnal modulation of hepatic lipid and glucose metabolism and whether GSPE effects on liver metabolism are impacted by the timing of administration. Results from hepatic lipid profiling, expression rhythm analysis of metabolic genes together with liver metabolomics in rats revealed that the CAF diet impaired glucose homeostasis and enhanced lipogenesis in the liver, leading to the generation of hepatosteatosis. Chronic consumption of GSPE at the onset of the active phase was able to restore the daily oscillation of liver mass and of key lipogenic and glycogenic genes, along with the reestablishment of liver metabolite rhythms, demonstrating hepatoprotective properties by decreasing triglyceride accumulation and lipid droplet formation in the liver, thus mitigating the development of CAF-induced NAFLD. Furthermore, in vitro data suggest that catechin, one of the main phenolic compounds found in the GSPE extract, may be involved in the ameliorating effects of GSPE against NAFLD.

## 1. INTRODUCTION

In modern times, as a consequence of the rising prevalence of obesity, the metabolic syndrome (MS) has become a common metabolic disorder [1]. It is comprised of several metabolic abnormalities, such as increased visceral adiposity, dyslipidemia, hyperglycemia, and hypertension. MS is also characterized by elevated levels of circulating triglycerides, lower levels of high-density lipoprotein cholesterol, and impaired fasting blood glucose concentrations [2]. Consequently, these changes increase the risk of developing type 2 diabetes and cardiovascular disease [3]. In the liver, MS manifests as nonalcoholic fatty liver disease (NAFLD), which includes a disease spectrum varying from simple steatosis to non-alcoholic steatohepatitis (NASH) and cirrhosis, with insulin resistance as an important pathogenetic hallmark [4]. Nowadays, excess of food intake and lack of physical activity are the main basis of the growing worldwide epidemic of metabolic disorders and NAFLD [5]. Moreover, artificial light at night and late-time work together with sleep deprivation are principal features of the industrialized world contributing to the pathogenesis of MS [6]. Particularly, it has been suggested that these changes cause circadian rhythm disturbances with a resulting upregulation of appetite leading to alterations in lipid and glucose metabolism[6–8]. Additionally, there is now increasing evidence connecting alterations of circadian rhythms with the key components of MS [9–11], suggesting that the circadian system is one of the main regulators of metabolism [12,13].

As a major metabolic organ, the liver is responsible for glucose and lipid homeostasis, where opposite metabolic processes such as glycolysis/gluconeogenesis, and lipogenesis/fatty acid oxidation take place, requiring the need for temporal separation [14]. Therefore, hepatic function, besides being under control of the insulin-glucagon signaling network, is tightly regulated by circadian clocks [15]. It has been described that not only transcripts, but also enzymes and proteins, which are important regulators of metabolic processes, undergo circadian oscillation in the liver [16,17]. Moreover, it is known that many hepatic metabolites oscillate in a daily manner [18].

The master circadian pacemaker, the suprachiasmatic nucleus (SCN) located in the hypothalamus, is influenced primarily by light, while peripheral circadian clocks located in tissues including the liver can be also regulated by the time of feeding and type of diet [19]. It has been demonstrated in mice that the intake of a high-fat diet (HFD) generates a profound reorganization of the liver clock reprogramming, in consequence, the oscillation of metabolic and transcriptional liver pathways. Diurnal rhythms of transcripts and metabolites involved in insulin signaling pathways, fatty acid and primary bile acid biosynthesis were found altered due to HFD [20]. Furthermore, it has been shown that the reset of the liver clock was specifically due to the nutritional challenge (HFD), and not due to the development of obesity. Of note, such dietary effects on the circadian clock are reversible [20].

Polyphenols, and specifically proanthocyanidins, are naturally occurring secondary plant metabolites that can be found in many fruits and vegetables, mostly in apples, cinnamon, grape seed and skin [21,22]. Proanthocyanidins are known to possess antioxidant, antimicrobial, anti-inflammatory, antiallergic, anti-obesity and vasodilatory properties [23,24]. Epidemiological evidence has linked the consumption of proanthocyanidins to a reduced risk of chronic diseases, including certain types of cancer and cardiovascular disease, as well as NAFLD [25]. Moreover, it has been shown that grape seed proanthocyanidin extract (GSPE) exhibits hepatoprotective effects in animal models of diet-induced obesity, helping to prevent the progression of steatosis and NAFLD [26–28]. Growing evidence suggests that polyphenols may also affect circadian rhythms by acting on the SCN and peripheral tissue circadian clock gene expression [29–32]. Among other issues, the bioavailability and the timing of administration arise as important features when studying the polyphenols effect on metabolic dysfunctions like MS [33]. The main constituents of GSPE are flavan-3-ol monomeric, dimeric, and trimeric procyanidins, with relatively low proportions of larger polymers [34]. In this regard, based on its low molecular weight procyanidin composition, GSPE has been considered to be highly bioavailable [35]. However, most studies do not consider the timing of administration, leaving this variable largely unexplored.

Taking into consideration the above explained factors and based on our previous results, in which we demonstrate that GSPE was able to restore hepatic circadian clock synchrony in cafeteria diet (CAF) fed rats [36], we further investigated the implications of obesity and MS in the close relationship between the hepatic molecular clock metabolic function, and energy regulation. Therefore, the aim of the present study was to investigate how an obesogenic diet intake affects the rhythm of hepatic lipid and glucose metabolism of rats and to determine whether. GSPE’s restorative effects on liver rhythms are impacted by the timing of administration. To achieve this, rats were fed with standard chow (STD) or cafeteria (CAF) diet for 9 weeks. During the last 4 weeks they were treated in the morning (ZT0) or at night (ZT12) with vehicle (VH) or 25 mg/kg of GSPE. Animals were sacrificed one hour after turning the light on (ZT1) and every 6 hours (ZT1, ZT7, ZT13 and ZT19) to assess diurnal transcript and metabolite profiles. Our data suggest a vital role for the circadian clock machinery and key metabolic gene rhythms in the generation and propagation of NAFLD, and a time-of-day dependent effect of GSPE in the amelioration of hepatic steatosis.

## 2. MATERIALS AND METHODS

### 2.1. Grape seed proanthocyanidin extract

Grape seed proanthocyanidin extract (GSPE) was obtained from Les Dérives Résiniques et Terpéniques (Dax, France). GSPE was directly analyzed by LC-MS/MS (Agilent Technologies, Palo Alto, CA, USA). A ZORBAX SE-aq column (150 mm×2.1 mm i.d., 3.5 μm particle size, Aglient Technologies) was used. The mobile phase consisted of (A) 0.2 % acetic acid in water and (B) acetonitrile. The gradient mode was as follows: initial conditions, 5 % B; 0-10 min, 5-55 % B; 10-12 min, 55-88 % B; 12-15 min, 80 % B isocratic, and 15-16 min, 80-85 % B. A post-run of 10 min was required for column re-equilibration. The flow rate was set at 0.4 mL/min, and the injection volume was 2.5 μL for all runs. Electrospray ionization (ESI) was conducted at 350 °C, the flow rate was 12 L/min with a nebulizer gas pressure of 45 psi, and the capillary voltage was 4,000 V. The mass spectrometer was operated in negative mode, and the MS/MS data were acquired in multiple reaction monitoring (MRM) mode. MRM conditions for the analysis of the phenolic compounds studied using HPLC-ESI-MS/MS were conducted as previously described [37]. The total polyphenol content, the individual flavanols and the phenolic acids comprising the extract are detailed in Table 1.

**Table 1.** Main phenolic compounds (flavanols and phenolic acids) of the grape seed proanthocyanidin extract (GSPE) used in this study, analyzed by HPLC-MS/MS.

| Phenolic compound | (M-H)- | Calibration curve | Total amount (mg/g) | SD |
| --- | --- | --- | --- | --- |
| Protocatechuic acid (PCA) | 153.0187 | $y = 1E+06x$ | 1.40 | 0.25 |
| Catechin | 289.0712 | $y = 1E+06x$ | 51.88 | 5.56 |
| Epicatechin | 289.0712 | $y = 935152x$ | 62.86 | 8.32 |
| Gallic acid | 169.0136 | $y = 287040x$ | 44.66 | 7.76 |
| Kaempferol-3-glucoside | 447.0927 | $y = 1E+06x$ | 0.50 | 0.02 |
| Naringenin-7-glucoside | 433.1135 | $y = 3E+06x$ | 0.64 | 0.08 |
| p-Coumaric acid | 163.0395 | $y = 2E+06x$ | 0.09 | 0.01 |
| Quercetin | 301.0348 | $y = 2E+06x$ | 0.05 | 0.01 |
| Quercetin-3-O-galactoside | 463.0877 | $y = 2E+06x$ | 0.43 | 0.05 |
| Vanillic acid | 167.0342 | $y = 1E+06x$ | 0.09 | 0.01 |
| Procyanidin dimer | 577.1346 | $y = 664077x$ | 76.84 | 15.76 |
| Procyanidin trimer | 865.1979 | $y = 664077x$ | 13.04 | 0.64 |
| Procyanidin tetramer | 1153.261<br>3 | $y = 664077x$ | 5.14 | 0.28 |
| Dimer gallate | 729.1455 | $y = 1E+06x$ | 15.22 | 2.72 |
| Epicatechin gallate | 441.0821 | $y = 1E+06x$ | 14.24 | 2.76 |
| Epigallocatechin gallate | 457.077 | $y = 1E+06x$ | 0.06 | 0.01 |

### 2.2. Animal handling

All procedures involving the use and care of animals were reviewed and approved by the Animal Ethics Committee of the Universitat Rovira i Virgili (permit number 9495, 18^th^ of September 2019). All experiments were performed in accordance with relevant guidelines and regulations of the Council of the European Union and the procedure established by the Departament d’Agricultura, Ramaderia i Pesca of the Generalitat de Catalunya.

Ninety-six 12-week-old male Fischer 344 rats were purchased from Charles River (Barcelona, Spain). The animals were housed in pairs in a 12-h/12-h light-dark cycle at 22 °C, 55 % humidity, and were provided a standard chow diet (STD) and tap water *ad libitum*. The STD diet composition was 20 % protein, 8 % fat and 72 % carbohydrates (Panlab, Barcelona, Spain). After a 4-day adaptation period, the animals were randomly divided into two groups depending on the diet; 32 rats were fed with STD, and 64 rats were fed a cafeteria diet (CAF) during a 5-week pre-treatment. CAF consisted of biscuits with cheese and pâté, bacon, coiled puff pastry from Mallorca (Hacendado, Spain), feed, carrots, and sweetened milk (22 % sucrose w/v). CAF composition was 14 % protein, 35 % fat and 76 % carbohydrates. After the pre-treatment period, rats were divided into two groups according to the Zeitgeber time (ZT) when the treatment was administered: 48 rats were treated at the beginning of the light phase (8 a.m., ZT0), and 48 rats were treated at the beginning of the dark phase (8 p.m., ZT12). The treatment period started in the 5th week and lasted for 4 weeks. Rats continued with the diet they were fed during the pre-treatment period. All STD-fed rats were treated with commercial sweetened skim condensed milk (Nestle; 100 g: 8.9 g protein, 0.4 g fat, 60.5 g carbohydrates, 1175 kJ) as vehicle (VH). CAF-fed rats were divided into two groups; 32 were treated with VH, and 32 with 25 mg/kg GSPE diluted 1/5 in condensed milk. The treatment was orally administered daily using a syringe. The rats were fasted for 3 h, then sacrificed by decapitation under anesthesia (sodium pentobarbital, 50 mg/kg per body weight). Each diet-treatment group was divided into 4 sub-groups (n=4), depending on the time of sacrifice (9 a.m. (ZT1), 3 p.m. (ZT7), 9 p.m. (ZT13) or 3 a.m. (ZT19)). Part of the liver was fixed with 10 % formalin, the rest of the liver tissue was quickly frozen in liquid nitrogen and then stored at −80 °C for further analysis.

### 2.3. RNA extraction

A liver tissue portion (20–30 mg) was mixed with TRIzol reagent (Thermo Fisher, Madrid, Spain) and homogenized by Tissue Lyser LT (Qiagen, Madrid, Spain). After a 10-min centrifugation (12,000 × g and 4 °C), the homogenate was placed into a new Eppendorf tube, and 120 μL of chloroform was added. Two phases were separated after a 15-min centrifugation (12,000 × g and 4 °C). The aqueous phase was transferred into a new Eppendorf tube, and 300 μL of isopropanol was added. After an overnight incubation at −20 °C, samples were centrifugated at 4 °C and 12,000 × g for 10 min. The supernatant was discarded, and the pellet was cleaned twice with 500 μL of ethanol 70 % and centrifuged for 5 min (8,000 × g and 4 °C). The supernatant was discarded again, and the washed pellet was resuspended with 60 μL of nuclease-free water (Thermo Fisher, Madrid, Spain). RNA concentration (ng/μL) and purity were measured by a Nanodrop ND-1000 spectrophotometer (Thermo Fisher, Madrid, Spain).

For AML12 *Bmal1-luc* cells, 750 μL TRIzol reagent (Thermo Fisher, Germany) were added to each 35-mm well and, after a 5 min incubation on ice, cells were collected, placed into Eppendorf tubes and 150 μL of chloroform was added. Two phases were separated after a 15-min centrifugation (12,000 × g and 4 °C). The aqueous phase was transferred into a new Eppendorf tube, and 375 μL of isopropanol was added. After an overnight incubation at −20 °C, samples were centrifugated at 4 °C and 12,000 × g for 40 min. The supernatant was discarded, and the pellet was cleaned twice with 350 μL of ethanol 70 % and centrifuged for 15 min (12,000 × g and 4 °C). The supernatant was discarded again, and the washed pellet was resuspended with 15 μL of nuclease-free water (Thermo Fisher, Madrid, Spain). RNA concentration (ng/μL) and purity were measured by a Nanodrop spectrophotometer (Thermo Fisher, Germany).

### 2.4. Gene-expression analysis

Complementary deoxyribonucleic acid (cDNA) was obtained by reverse transcription of the RNA extracted using a high-capacity complementary DNA reverse-transcription kit (Thermo Fisher, Madrid, Spain). Quantitative polymerase chain reactions (qPCRs) were performed in 384-well plates in a 7900HT fast real-time PCR (Thermo Fisher, Madrid, Spain) using iTaq Universal SYBR Green Supermix (Bio-Rad, Barcelona, Spain). The cycle program used in all qPCRs was 30 s at 90 °C, 40 cycles of 15 s at 95 °C and 1 min at 60 °C. The analyzed liver genes were normalized by the housekeeping gene peptidylprolyl isomerase A (*Ppia*) and the cell samples were normalized by the housekeeping gene Eukaryotic translation elongation factor 1 alpha (*eEf1a*). No marked changes in cycle thresholds (Cts) of the housekeeping genes were seen across the different times of sacrifice among the groups or cells samples. The primers used for each gene were obtained from Biomers (Ulm, Germany) and can be found in Supplementary Tables (ST) 1 and ST2. The relative expression of each gene was calculated using the 2^-ΔΔCt^ method, as reported by Schmittgen and Livak [38].

### 2.5. Liver lipid profiling

Liver lipids were extracted following the Bligh–Dyer method [39], and levels of hepatic cholesterol, TAG and total lipid liver content were measured using a colorimetric kit assay (QCA, Barcelona, Spain).

### 2.6. Liver histology

Liver portions fixed in buffered formalin (4% formaldehyde, 4 g/L NaH_2_PO_4_, 6.5 g/L Na_2_HPO_4_; pH 6.8) were cut at a thickness of 3.5 μm and stained with hematoxylin & eosin (H&E). Liver images (magnification 40X) were taken with a microscope (ECLIPSE Ti; Nikon, Tokyo, Japan) coupled to a digital sight camera (DS-Ri1, Nikon) and analyzed using ImageJ NDPI software (National Institutes of Health, Bethesda, MD, USA; https://imagej.nih.gov/ij, version 1.52a). To avoid any bias in the analysis, the study had a double-blind design, preventing the reviewers from knowing any data from the rats during the histopathological analysis. A general NAFLD scoring system was established to diagnose rats with NAFLD/NASH. The key features of NAFLD and NASH were categorized as follows: steatosis was assessed by analyzing macrovesicular (0–3) and microvesicular steatosis (0–3) separately, followed by hepatocellular hypertrophy (0–3), which evaluates abnormal cellular enlargement, and finally giving a total score of 9 points of steatosis state. Inflammation was scored by counting cell aggregates (inflammatory foci). The score 0 to 3 depends on the grade of the feature. It is categorized as 0 (66 %), and this scoring is used in each feature of steatosis and then added to the total steatosis score [40]. Ballooning was not included in the scoring system, because only quantitative measures were considered for the rodent NAFLD score. It is important to highlight that hypertrophy is not a sign of cellular injury and slightly refers to an anomalous enlargement of the cells without recognizing the source of this enlargement [40].

### 2.7. Metabolomic analysis

Metabolomic analysis of the 96 rat liver samples was performed at the Centre for Omic Sciences (COS, Tarragona, Spain) using gas chromatography coupled with quadrupole time-of-flight mass spectrometry (GC-qTOF model 7200, Agilent, Santa Clara, CA, USA). The extraction was performed by adding 400 μL of methanol:water (8:2)-containing internal standard mixture to liver samples (approx. 10—20 mg). Then, the samples were mixed and homogenized on a bullet blender using a stainless-steel ball, incubated at 4 °C for 10 min and centrifuged at 19.000× *g* rpm; supernatant was evaporated to dryness before compound derivatization (methoximation and silylation). The derivatized compounds were analyzed by GC-qTOF. Chromatographic separation was based on the Fiehn Method [41] using a J&W Scientific HP5-MS film capillary column (30 m × 0.25 mm × 0.25 μm, Agilent, Santa Clara, CA, USA) and helium as carrier gas with an oven program from 60 to 325 °C. Ionization was done by electronic impact (EI) with an electron energy of 70 eV and operating in full-scan mode. Identification of metabolites was performed using commercial standards and by matching their EI mass spectrum and retention time to a metabolomic Fiehn library (from Agilent, Santa Clara, CA, USA) which contains more than 1,400 metabolites. After putative identification of metabolites they were semi-quantified in terms of internal standard response ratio.

### 2.8. Alpha mouse liver 12 (AML-12) Bmal1-luc maintenance

AML-12 wild-type cells were originally obtained from ATCC biobank (cat number CRL-2254) and grown in DMEM/F12 media containing 15 mM of HEPES, 1 % of penicillin-streptomycin (10,000 units/mL and 10,000 μg/mL, respectively), 1 % of insulin-transferrin-selenium (ITS), 10 % of non-heat inactived serum (all items from Thermo Fisher, Germany). Dexamethasone at 10 nM (Sigma-Aldirch, USA) was supplemented for regular culture maintenance.

To generate *Bmal1-luc* cells, HEK 293 cells were transfected using CaCl_2_ solution containing *Bmal1-luc* plasmid (17.5 μg, pABpuro-Bluf [42]) together with packing plasmids psPax and pMD2G (12.5 and 7.5 μg, respectively). Supernatant containing virus was collected 48 h later and 10-X concentrated using Lenti-X Concentrator (Takara Bio, Germany). AML-12 wild-type cells were transduced using concentrated virus and selected using 3 μg/mL of puromycin. From the heterogenous cell culture, single clones were identified and isolated. AML-12 *Bmal1-luc* clone 4 was selected and used for subsequent assays.

### 2.9. *In vitro* model of NAFLD by supplementing palmitate

AML-12 *Bmal1-luc* cells were grown in the above-mentioned media without phenol red and dexamethasone. Serum was replaced for charcoal-dextran treated serum that has a significant reduction in endogenous bovine hormones (HyClone, USA). Cells were pre-treated with (+)-catechin or (-)-epicatechin 10 μM or 100 μM for 72 h, using dimethylsulfoxide (DMSO) as a solvent 0.01 % (v/v) or 0.1% (v/v), respectively. 2×10^5^ cells were then spited and seeded in 35 mm dishes containing medium with (+)-catechin or (-)-epicatechin 10 μM or 100 μM. Twenty-four h later, cells were synchronized using 200 nM of dexamethasone for 2 h. Cells were washed with PBS and loaded with media containing (+)-catechin or (-)-epicatechin 10uM or 100uM. To mimic a high-fat scenario *in vitro*, cells were loaded with sterile bovine serum albumin (BSA) conjugated with palmitate (0.25 mM) or with serum albumin (0.04 mM) as control, both with 0.1 % (v/v) DMSO. In all conditions, stocks solutions were highly concentrated and underwent serial dilution to remove organic solvents. Stock solution of 5 mM palmitate:0.8 mM BSA was acquired from Cayman, USA and filtered using a 0.22-μm filter. Luciferin (200 μM) was added to each dish, which were covered with round glass lids, and sealed with parafilm. All dishes were transferred to the Lumicycle and bioluminescence was measured every 10 minutes (Actimetrics, USA) at 32.5 °C for 4 days. Bioluminescence measurements were collected, and the raw data were imported to the Lumicycle software. Baseline subtracted bioluminescence, calculated by 24-hour running-average subtraction, was used for rhythm evaluation.

### 2.10. Rhythm analysis

Rhythmic analyzes were performed using CircaCompare algorithm [43]. Rhythmic parameters were compared in a pairwise fashion using CircaCompare. Presence of rhythmicity was considered when *p*-value < 0.05 was achieved. Comparison between amplitude and MESOR was performed for genes or metabolites, regardless of rhythmicity status. Phase estimation was performed only on gene or metabolites that showed significant rhythms in both groups.

For bioluminescence data, rhythmicity was assessed using the Circasingle algorithm as previously described [44] using the normalized bioluminescent data per hour. CircaSingle algorithm was set to consider amplitude a decay factor and period was not pre-established. Rhythmicity was confirmed when *p*-value < 0.05.

### 2.11. Statistical analysis

All data, except for bioluminescence data, were log2 transformed. The data displayed in all figures and tables represents mean ± standard error of the mean (SEM). Hepatic lipids, metabolites and gene expression oscillations were analyzed as described above. Global metabolomics analysis was performed using one-way ANOVA, followed by Tukey’s post-test in R environment. PCA analyzes were performed using the factoextra package and Hartigan-Wong, Lloyd, and Forgy MacQueen algorithms (version 1.0.7) in R. Liver lipid profiles were subjected to Student’s t test and one-way ANOVA followed by Turkey’s multiple-range post-hoc test using IBM SPSS Statistics (version 25) software. Graphics were done in GraphPad Prism 9 software (San Diego, CA, USA) and R using ggplot package. For all analyses, a probability (*p*) value of < 0.05 was considered to reflect a statistically significant difference.

## 3. RESULTS

### 3.1. Hematoxylin & eosin staining demonstrates that CAF diet induces NAFLD and a time dependent effectiveness of GSPE treatment

H & E staining was performed to study the pathological consequences of CAF diet intake on the liver tissue. As is shown in Figure 1 A-B compared to STD liver tissue, lipid droplets were visible in the liver and many hepatocytes had swelling and injury in CAF-VH groups, in both ZT0 and ZT12 treatments. Analysis of H & E staining showed that there was a significant difference in the histopathology and lipid accumulation between STD diet liver tissues and liver tissues of CAF diet animals. Interestingly, GSPE treatment was able to reduce the numbers of lipid droplets when it was administered at ZT12 (p = 0.01) but not at ZT0, lowering the level of inflammation from 0.75 to 0.44 and steatosis score from 3.17 to 2.67 changing the NASH status to NAFLD only at ZT12 (Figure 1 C, Table 2).

**Figure 1.**
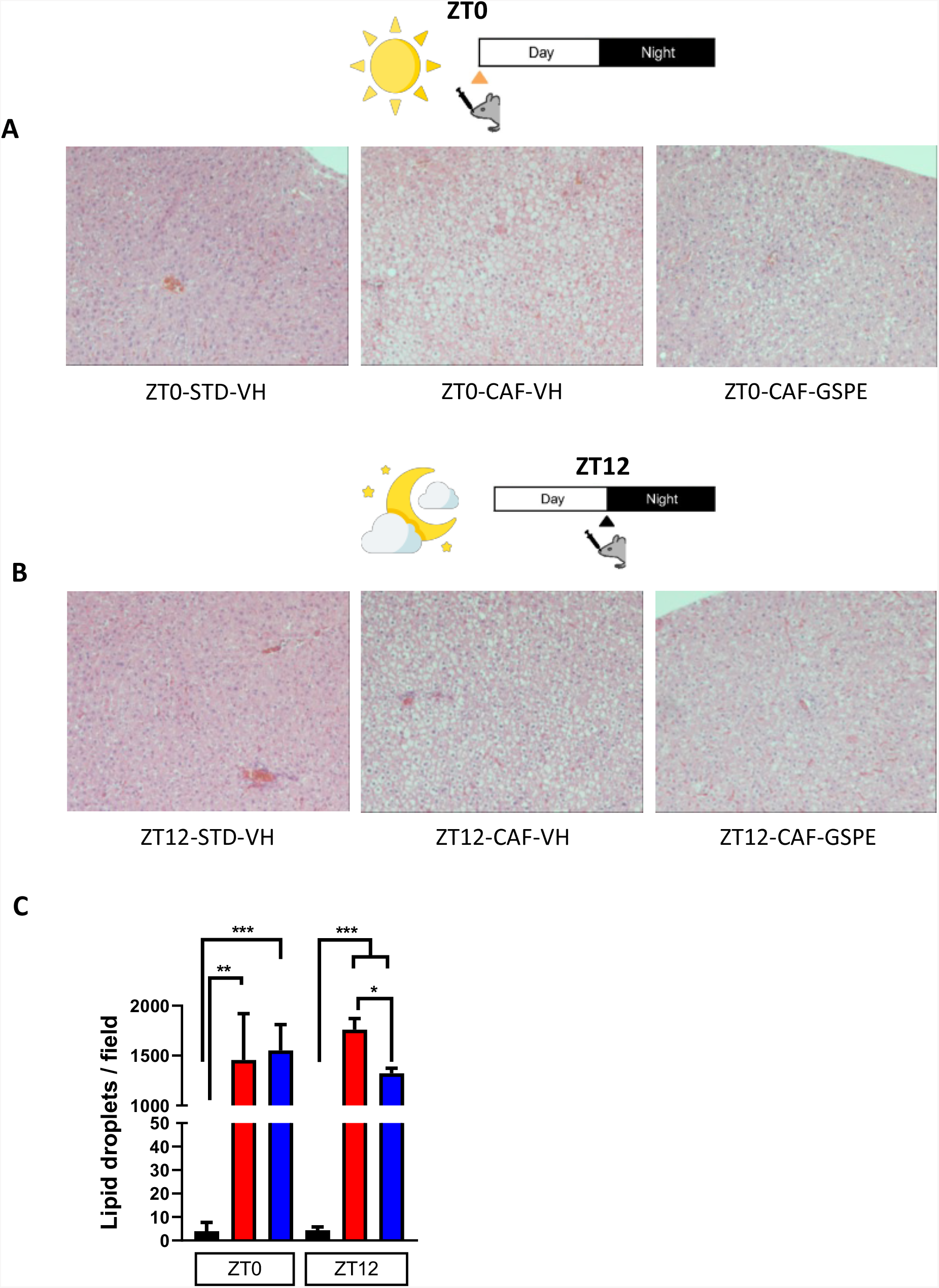
Representative pictures of H&E-stained liver sections. Scale bar, 100 μm. Rats were fed a STD or CAF diet and received a daily dosage of vehicle or GSPE at the beginning of the light phase (ZT0) **(A)** or at the beginning of the dark phase (ZT12) **(B). C** Quantitative analysis of lipid droplets in liver samples per microscopic field (100×). * Indicates significant differences (Student’s t test, *p<0.05, **p<0.01, ***p<0.001). Shown are means ± SEM (n=4/group at each time point).

**Table 2.** Analysis of liver hematoxylin & eosin staining of rats that were fed a STD or CAF diet and received a daily dosage of vehicle or GSPE at the beginning of the light phase (ZT0) or at the beginning of the dark phase (ZT12).

|  | ZT0 |  |  | ZT12 |  |  |
| --- | --- | --- | --- | --- | --- | --- |
|  | STD-VH | CAF-VH | CAF-GSPE | STD-VH | CAF-VH | CAF-GSPE |
| **Macrovesicular st. (0-3)** | 0.000 | 0.167 | 0.111 | 0.000 | 0.000 | 0.000 |
| **Microvesicular st.(0-3)** | 0.000 | 1.750 | 1.667 | 0.000 | 2.167 | 2.000 |
| **Hypertrophy (0-3)** | 0.000 | 0.667 | 0.333 | 0.000 | 1.000 | 0.667 |
| **Steatosis score (0-9)** | 0.000 | 2.583 | 2.111 | 0.000 | 3.167 | 2.667 |
| **Inflammation** | 0.417 | 0.500 | 0.500 | 0.250 | 0.750 | 0.444 |
| **Diagnosis** | No NAFLD | NASH | NASH | No NAFLD | NASH | NAFLD |
| **Lipid droplets number** | 3.9 | 1175.7 | 1590.7 | 4.4 | 1397.7 | 1323.5 |
| **Total volum (pixels<sup>3</sup>)** | 1485.3 | 792289.4 | 970609.2 | 2883.8 | 1135575.0 | 1070155.1 |
| **Total surface (pixels<sup>2</sup>)** | 396872.7 | 195104404.3 | 251846625.1 | 754340.5 | 277785713.8 | 268128385.0 |
| **Average surface area (pixels<sup>2</sup>)** | 74895.3 | 133774.2 | 176223.3 | 93125.2 | 189227.5 | 204359.4 |

### 3.2. GSPE restores CAF-diet disrupted daily oscillations in liver weight

To obtain a temporal representation of the diurnal changes in liver lipid abundance, we measured levels of triglycerides (TAGs), total cholesterol, total lipid content and liver weight at 6-hour intervals across the day. Results from circadian analysis, including rhythmometric parameters can be found on ST3 and group comparisons in ST4 [43].

In the morning treatment (ZT0), cholesterol levels and total lipid content showed rhythmicity in STD-VH (p = 0.04 and 0.001, respectively) but not in CAF-VH and CAF-GSPE animals. On the other hand, no rhythmic pattern was found for hepatic triglycerides in any of the groups. Interestingly, diurnal oscillations in liver weight were observed in all groups. While liver weight in STD-VH and CAF-GSPE groups peaked around ZT10, CAF-VH animals livers showed maximum weight at ZT16, causing a phase delay of 6 hours (p = 0.001) (Figure 2 A-D).

**Figure 2.**
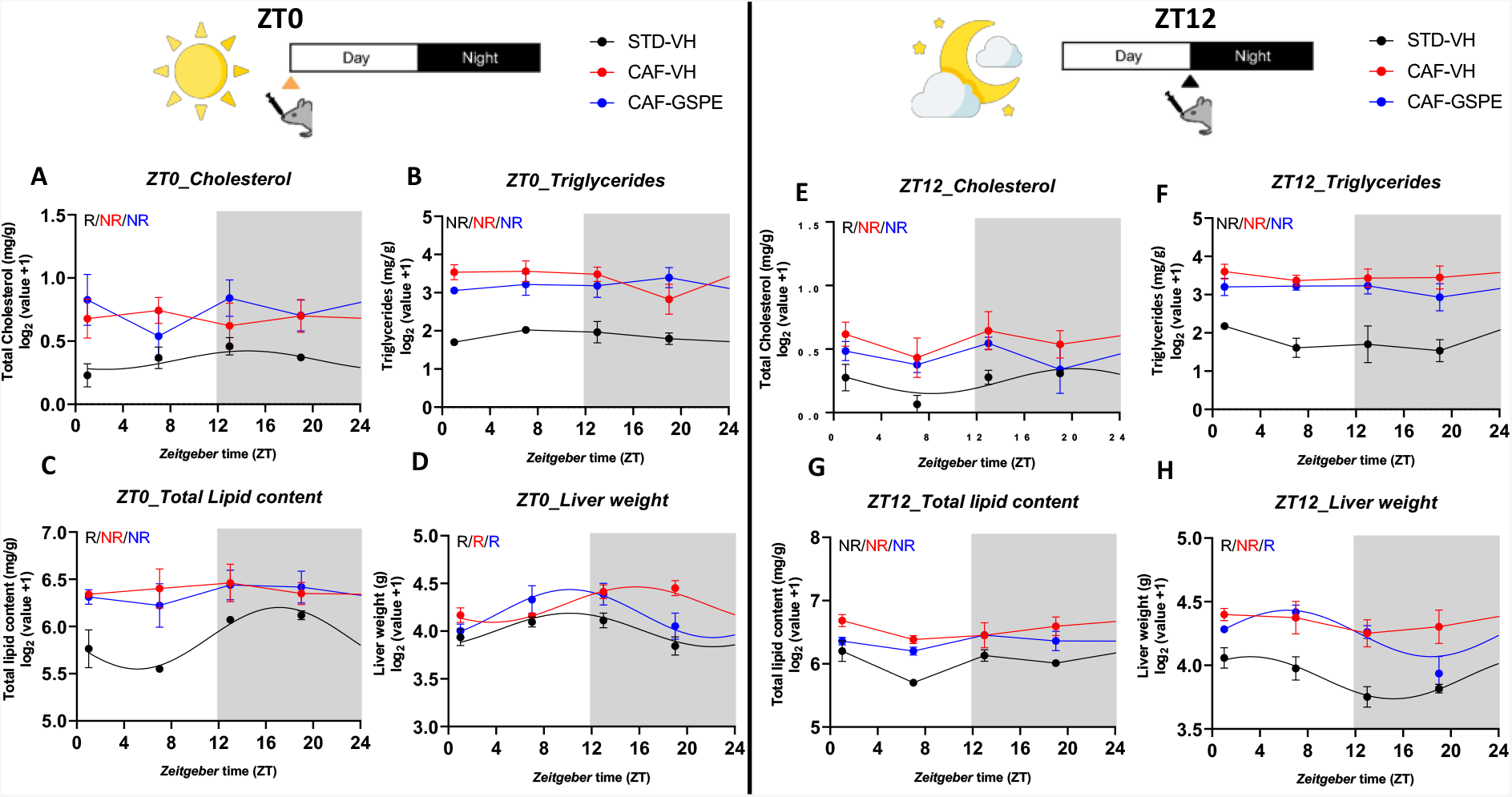
Diurnal rhythms of liver lipid levels and liver weight. Rats were fed a STD or CAF diet and received a daily dosage of vehicle or GSPE at the beginning of the light phase (ZT0) **(A-D)** or at the beginning of the dark phase (ZT12) **(E-H)**. Rhythm parameter determination and comparison were performed using CircaCompare. R/NR indicates significant/non-significant rhythmicity (p < 0.05). Shown are means ± SEM (n=4/group at each time point).

After GSPE night (ZT12) treatment (Figure 2 E-H), non-rhythmic expression was observed for total lipid content in all groups, whereas total cholesterol showed rhythmicity only in the STD-VH group (p = 0.03). Although hepatic triglycerides did not show a significant circadian oscillation neither in STD-VH nor in CAFs groups; a tendency of decreasing the overall expression (MESOR) was observed in CAF-GSPE compared to CAF-VH group (p = 0.058). This result indicates that GSPE tends to reduce hepatic triglyceride accumulation. Interestingly, animals subjected to ZT12 GSPE treatment showed lower hepatic triglycerides levels compared to CAF-VH when considering all ZTs together (p = 0.04). In addition, the CAF diet disrupted circadian oscillations of liver weight, but animals treated with GSPE recovered the pattern of rhythmic expression with a slight phase delay (p = 0.01). It is worth to mention that MESOR was higher in CAF fed animals (VH and GSPE) when compared to STD-VH rats for all hepatic lipid parameters, which might reflect the differences in fat and carbohydrate content of both diets.

### 3.3. Daily oscillations of lipid metabolism related genes are lost in CAF but reestablished by GSPE treatment

To investigate how the CAF diet affects liver lipid metabolism by influencing the circadian system, we analyzed oscillations in the expression levels of genes that play crucial roles in regulating lipid metabolism in the liver. All rhythm analysis results such as phases, amplitudes and MESOR are shown in ST5 and group comparisons in ST6.

SREBP-1c (sterol regulatory element binding protein 1c) is one of the main transcription factors controlling the expression of lipid metabolism related genes downstream of the core clock machinery. Amongst others it regulates rate-limiting enzymes of fatty acid biosynthesis [45]. In the morning treatment (ZT0), the mRNA levels of SREBP-1c showed a clear daily oscillation in the STD group (p = 0.009), which was lost in CAF-VH animals and re-established when rats were treated with GSPE (p = 0.001). *Acaca* (acetyl-CoA carboxylase alpha) and *Cd36* (cluster of differentiation 36), also known as fatty acid translocase, involved in fatty acid biosynthesis and transport [46], did not show rhythmic expression patterns neither in the STD-VH nor in the CAF-VH group, but they followed a circadian oscillation in CAF-GSPE treated animals (p = 0.002 and 0.01, respectively). A rhythmic expression of FATP5 (fatty acid transport protein 5) was observed in CAF-VH and CAF-GSPE with no differences between these two groups. For *Fasn* (fatty acid synthase), which contributes to *de novo* lipogenesis [47], our results showed a loss of rhythmicity caused by the CAF diet, and a restoration of its circadian oscillation when CAF rats were also treated with GSPE (p = 0.009). These animals also displayed a decrease in *Fasn* MESOR expression when compared to STD group (p = 0.001) (Figure 3-E). *Pparα* (peroxisome proliferator-activated receptor α) is a nuclear receptor that also connects the clock genes with lipid metabolism [48]. As is shown in figure 3-F all groups displayed diurnal rhythms for *Pparα*, although both CAF groups (VH and GSPE) showed an increase of the MESOR compared to the STD-VH (p < 0.001 and p = 0.002, respectively) and the GSPE group exhibited a phase delay of 4 hours compared to STD (p = 0.03). *Cyp7a1* (cytochrome P450 family 7 subfamily A member 1 also known as Cholesterol 7α-hydroxylase), is a gene that encodes for a protein involved in bile acid biosynthesis. It catalyzes the first reaction in the cholesterol catabolic pathway in the liver, which converts cholesterol to bile acids [49]. Non-rhythmic expression of *Cyp7a1* was seen in the STD group, but when comparing both CAF groups that showed a rhythmic pattern, CAF-VH exhibited a higher MESOR than GSPE (p = 0.04).

**Figure 3.**
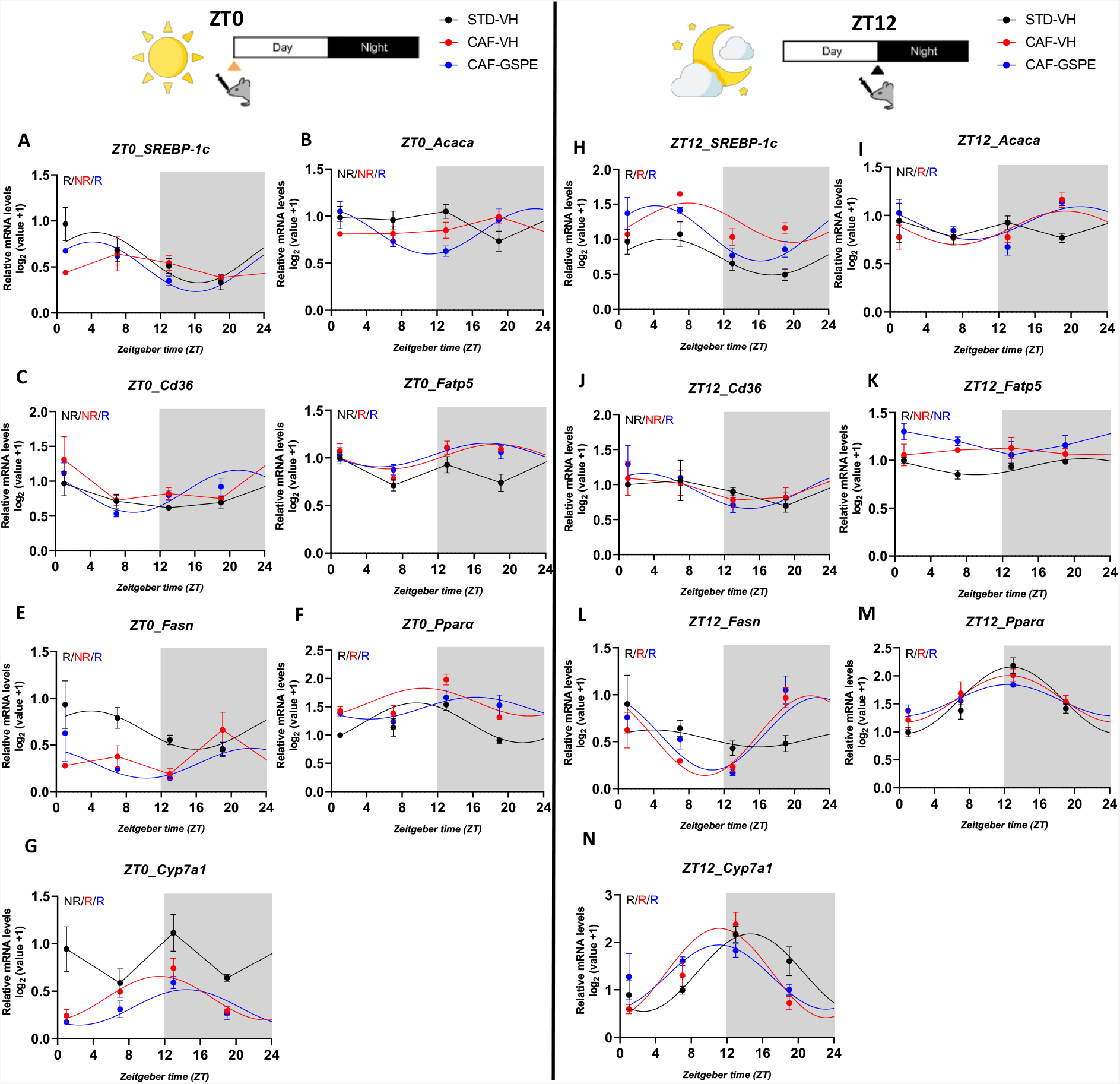
Diurnal mRNA profiles of key genes involved in lipid and bile acid metabolism. Rats were fed a STD or CAF diet and received a daily dosage of vehicle or GSPE at the beginning of the light phase (ZT0) **(A-G)** or at the beginning of the dark phase (ZT12) **(H-N)**. Rhythm parameter determination and comparison were performed using CircaCompare. R/NR indicates significant/non-significant rhythmicity (p < 0.05). Shown are means ± SEM (n=4/group at each time point).

In the case of night (ZT12) treatment (Figure 3 H-N), when compared to STD-VH, both CAF-VH and CAF-GSPE showed higher MESOR levels of SREBP-1c (p = 0.001 and 0.004, respectively). Its downstream target, *Acaca*, exhibited a 3-hour phase advance in the CAF-VH compared to CAF-GSPE group (p = 0.047), whereas non rhythmic expression for this gene was found in the STD group. Moreover, the amplitude of its mRNA oscillation was increased in rats treated with GSPE compared to the STD group (p = 0.048). Interestingly, CAF-GSPE rats were the only group that showed a circadian rhythm (p = 0.01) for *Cd36*, but for FATP5 only STD-VH showed rhythmic oscillation (p = 0.01). In the case of *Fasn* shifted phases induced by the CAF diet were seen - 5 h of phase shift when comparing STD-VH to CAF-VH (p = 0.009) and 4 h of phase shift if STD-VH and CAF-GSPE groups were compared (p= 0.05). *Pparα* showed diurnal rhythms in STD-VH (p = 0.001), CAF-VH (p = 0.002), and CAF-GSPE (p = 0.001) groups, although a reduction in the amplitude was seen in the CAF-GSPE group compared to STD-VH (p = 0.001). *Cyp7a1* mRNA expression displayed diurnal rhythmic variations in all groups, although a 3-hour phase advance was caused by CAF diet and was not re-adjusted by GSPE treatment. Additionally, amplitude was lower in CAF-GSPE than CAF-VH group (p = 0.04) for this gene.

### 3.4. Diurnal rhythms of glucose metabolic genes are disrupted by CAF-diet intake and restored by GSPE

To further explore the impact of CAF diet on hepatic glucose metabolism, we analyzed diurnal expression profiles of genes that are key regulators of glucose metabolism in the liver. All rhythmic analysis results (phases, amplitudes and MESOR) are shown in ST7 and group comparisons in ST8.

G6PC (glucose-6-phosphatase) is a key enzyme in glucose homeostasis, functioning in gluconeogenesis and glycogenolysis [50]. All experimental groups exhibited circadian oscillations for mRNA of its catalytic subunit (*G6pc*), however CAF-VH and GSPE showed a decrease in MESOR compared to STD-VH (p = 0.023 and 0.020, respectively) (Figure 4-A). *Slc2a2* (solute carrier family 2 member 2) encodes for the main liver glucose transporter, GLUT2 [51]. Although its mRNA levels exhibited rhythmicity in all groups, CAF-VH showed an increase in the MESOR compared to STD-VH (p = 0.02) and CAF-GSPE (p = 0.006). Expression of the mRNA of G6PD (glucose-6-phosphate dehydrogenase) which catalyzes the first rate-limiting step in the pentose phosphate pathway [52], showed a loss of its rhythmic expression under CAF diet, meanwhile GSPE treatment was able to restore this rhythm, albeit a decrease in MESOR compared to the STD group (p = 0.001). A major regulator of hepatic gluconeogenesis is Peroxisome proliferator-activated receptor gamma coactivator 1-alpha (encoded by *Ppargc1α*), which induces the transcription of gluconeogenic genes [53]. Only GSPE treated group exhibited rhythmicity of this gene (p = 0.04). Finally, when analyzing *Sirt1* (sirtuin 1), a nicotinamide adenosine dinucleotide (NAD)-dependent deacetylase that regulates glycogen synthesis and gluconeogenesis [54], non-significant rhythm was seen in STD group, but CAF-VH and CAF-GSPE groups showed a rhythmic expression (p = 0.02 and 0.04, respectively).

**Figure 4.**
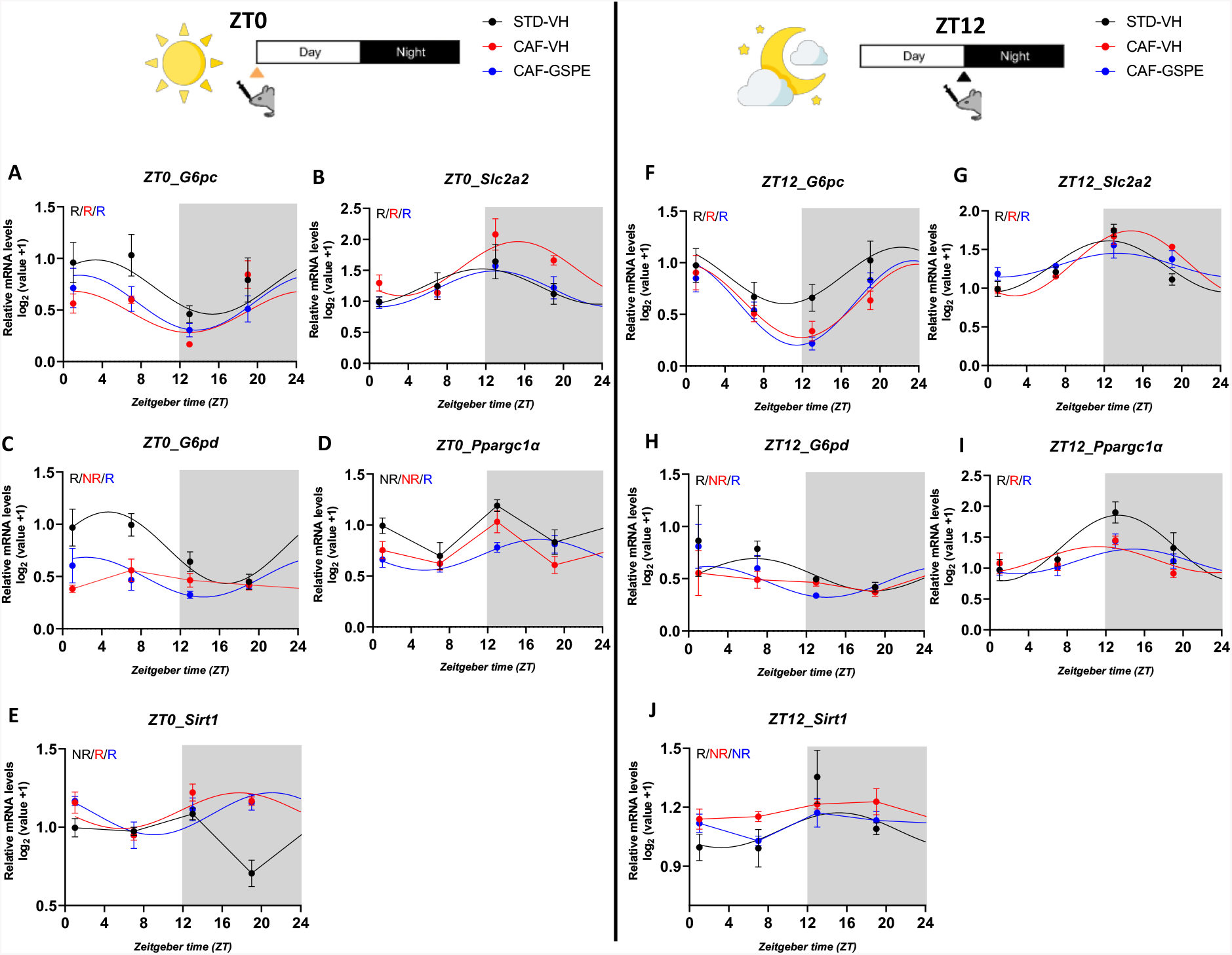
Diurnal mRNA profiles of key genes involved in glucose metabolism. Rats were fed a STD or CAF diet and received a daily dosage of vehicle or GSPE at the beginning of the light phase (ZT0) **(A-E)** or at the beginning of the dark phase (ZT12) **(F-J)**. Rhythm parameter determination and comparison were performed using CircaCompare. R/NR indicates significant/non-significant rhythmicity (p < 0.05). Shown are means ± SEM (n=4/group at each time point).

Regarding the night treatment (ZT12), mRNA levels of *G6pc* showed a clear diurnal oscillation in the three experimental groups, but a decrease of MESOR was found between the STD and both CAF groups (VH and GSPE; p = 0.02). For *Slc2a2*, a phase delay caused by the CAF diet was observed (p = 0.004) compared to STD group, whereas in GSPE treated animals this phase shift was not detected. In addition, a 2-fold decreased amplitude in CAF-GSPE rats compared to the CAF-VH group was observed. Intriguingly, circadian oscillation of G6PD was totally disrupted in CAF fed animals and restored by GSPE treatment (p = 0.01; Figure 4-H). *Ppargc1α* showed a clear diurnal rhythmicity in the STD group while CAF diet had a tendency to decrease its MESOR (p = 0.07) and amplitude (p = 0.09). Although there were no significant differences in phase between STD and CAF-VH groups, animals treated with GSPE peaked at the same time as the STD group (ZT13), whereas CAF-VH rats peaked at ZT11, i.e., 2-hour phase delay. *Sirt1* showed a rhythmic expression in STD animals (p = 0.01) but not in the two CAF diet groups.

### 3.5. Global analysis of hepatic metabolomic assay reveals GSPE increased key metabolites concentrations decreased due to CAF diet affecting several metabolic pathways

To gain direct insights into the effects of obesogenic diet intake on hepatic metabolism, we conducted a metabolomic analysis on liver tissue. 61 metabolites were identified and quantified in the liver of all experimental groups.

Principal component analyzes (PCA) showed a clear differentiation between the groups when treated at ZT0 and ZT12, thus arguing for a time-dependent effect of GSPE treatment (Supplementary Figure 1). As a first step, we conducted a global analysis without considering sacrifice time points (ZTs), to have a general view of metabolite concentrations. They are plotted in Figures 5 and 6 where statistically significant differences between groups are shown.

**Figure 5.**
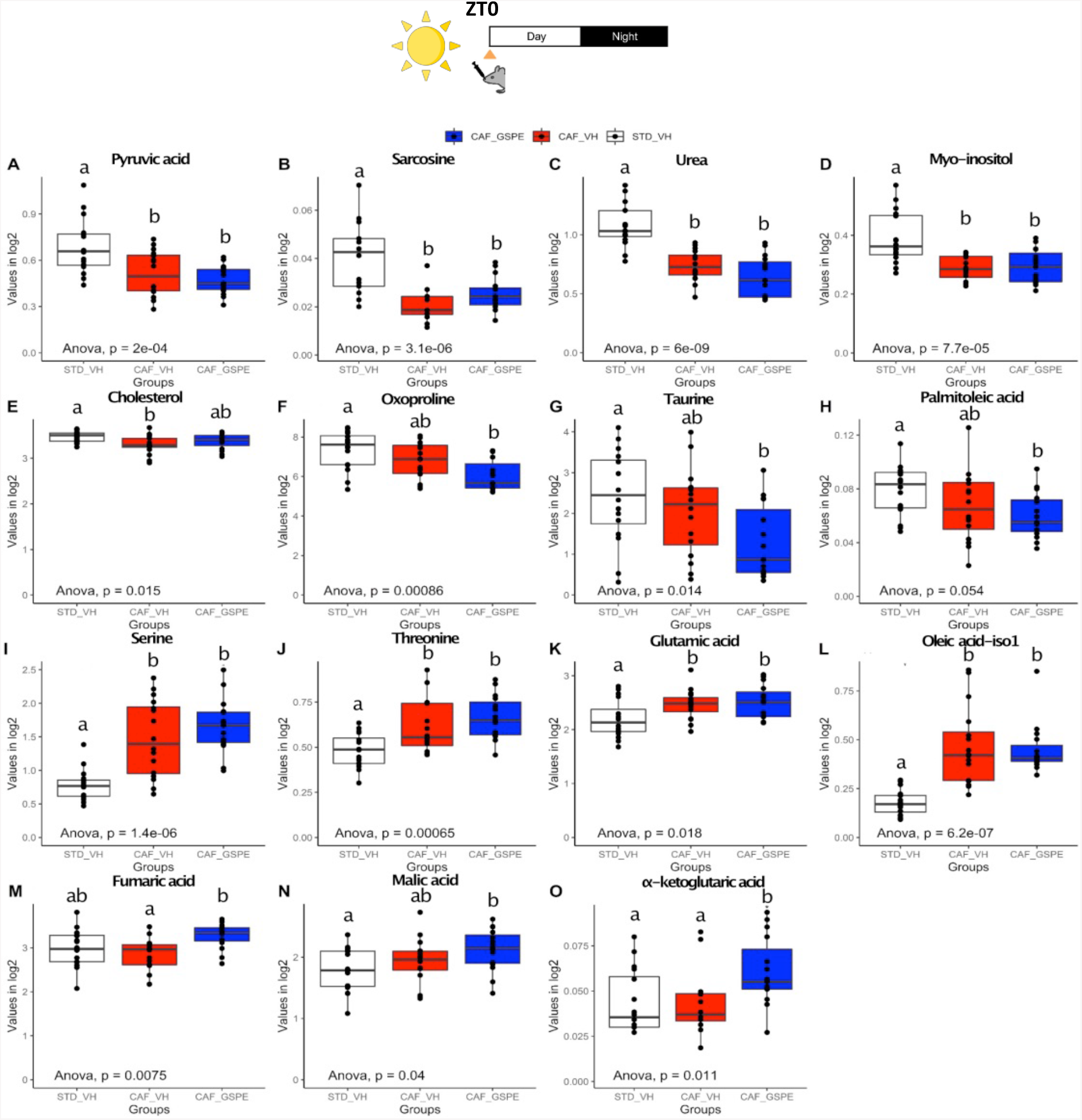
Box plots of metabolite levels in the liver of rats treated at ZT0. Rats were fed a STD or CAF diet and received a daily dosage of vehicle or GSPE at the beginning of the light phase (ZT0). One-way ANOVA followed by Tukey post-test were performed to compare the values between groups. Significant differences (p ≤ 0.05) are represented by different letters (a-c). Shown are means ± SEM (n=16/group).

**Figure 6.**
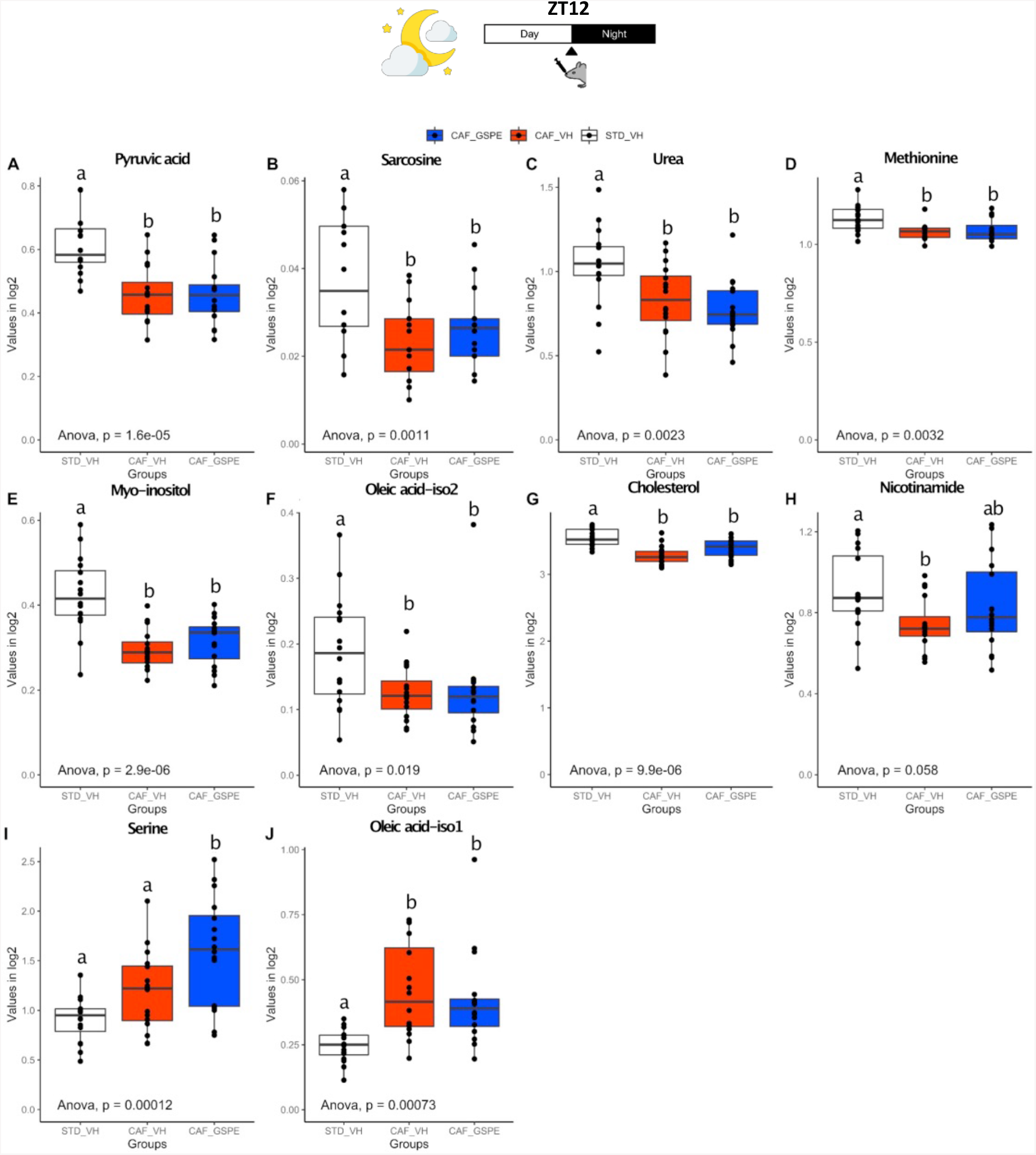
Box plots of metabolite levels in the liver of rats treated at ZT12. Rats were fed a STD or CAF diet and received a daily dosage of vehicle or GSPE at the beginning of the light phase (ZT12). One-way ANOVA followed by Tukey post-test were performed to compare the values between groups. Significant differences (p ≤ 0.05) are represented by different letters (a-c). Shown are means ± SEM (n=16/group).

In the morning treatment ZT0 (Figure 5), CAF diet lowered the levels of pyruvic acid, sarcosine, urea, myo-inositol, and cholesterol when compared to the STD-VH group, whereas CAF-GSPE did not significantly reduce the levels of sarcosine, myo-inositol and cholesterol but instead showed a reduction on the levels of oxoproline, taurine and palmitoleic acid in addition to pyruvic acid and urea when compared to the STD-VH group. Levels of serine, threonine, glutamic acid, and oleic acid iso1 were higher in both CAF groups when compared to STD-VH. Meanwhile GSPE treatment increased concentration of fumaric acid with respect to CAF-VH, malic acid compared to STD-VH and α-ketoglutaric acid relative to STD and CAF-VH groups.

Analysis of metabolites in the night treatment (ZT12) showed that the CAF groups (VH and GSPE) had lower levels of pyruvic acid, sarcosine, urea, methionine, myo-inositol, oleic acid iso2, and cholesterol than the STD-VH group (Figure 6). Interestingly, nicotinamide levels were decreased in CAF-VH (p = 0.046) but not in animals treated with GSPE compared to STD-VH. In addition, the concentrations of serine in CAF-GSPE rats were sharply increased when compared to CAF (p = 0.023) and STD VH (p = 0.001). Meanwhile, oleic acid iso1 was the only metabolite increased in both CAF groups.

### 3.6. Rhythmicity of liver metabolites is disrupted by CAF diet and partially restored by GSPE

Since the coordinated oscillations of metabolites are critical for the optimal functioning of liver metabolism, we analyzed the diurnal expression of the 61 quantified hepatic metabolites using the CircaCompare algorithm [43] (ST9 and ST10).

A Venn Diagram was created only with those metabolites that displayed statistically significant rhythms (Figure 7). Among them, we selected those that showed an exclusive rhythmic expression in each group as it was reported that a HFD causes an unexpected genesis of *de novo* oscillating metabolites, that lead to a reprogramming of the metabolic liver pathways [20]. In addition, an enrichment analysis was performed to identify the pathways in which these metabolites were involved (Figure 8). In STD-VH animals treated at ZT0 rhythmic metabolites showed enrichment for urea cycle, mitochondrial transport chain, and arginine and proline metabolism. On the other hand, in CAF-VH treated animals rhythmic metabolites showed an enrichment for lactose and galactose metabolism, while malate-aspartate shuttle, glucose-alanine cycle and alanine metabolism were found enriched in CAF-GSPE group (Figure 8A; ST11). Upon animals treated at ZT12, pathways that were enriched in STD-VH group were mainly associated with thiamine, riboflavin, and alanine metabolism, among others. While glycine and serine metabolism, malate-aspartate shuttle, glucose-alanine cycle, and glutathione metabolism were the metabolic pathways that were observed to be enriched in CAF-VH group. In the case of CAF-GSPE group no significant enriched pathway was found among its exclusive rhythmic metabolites. (Figure 8B; ST11).

**Figure 7.**
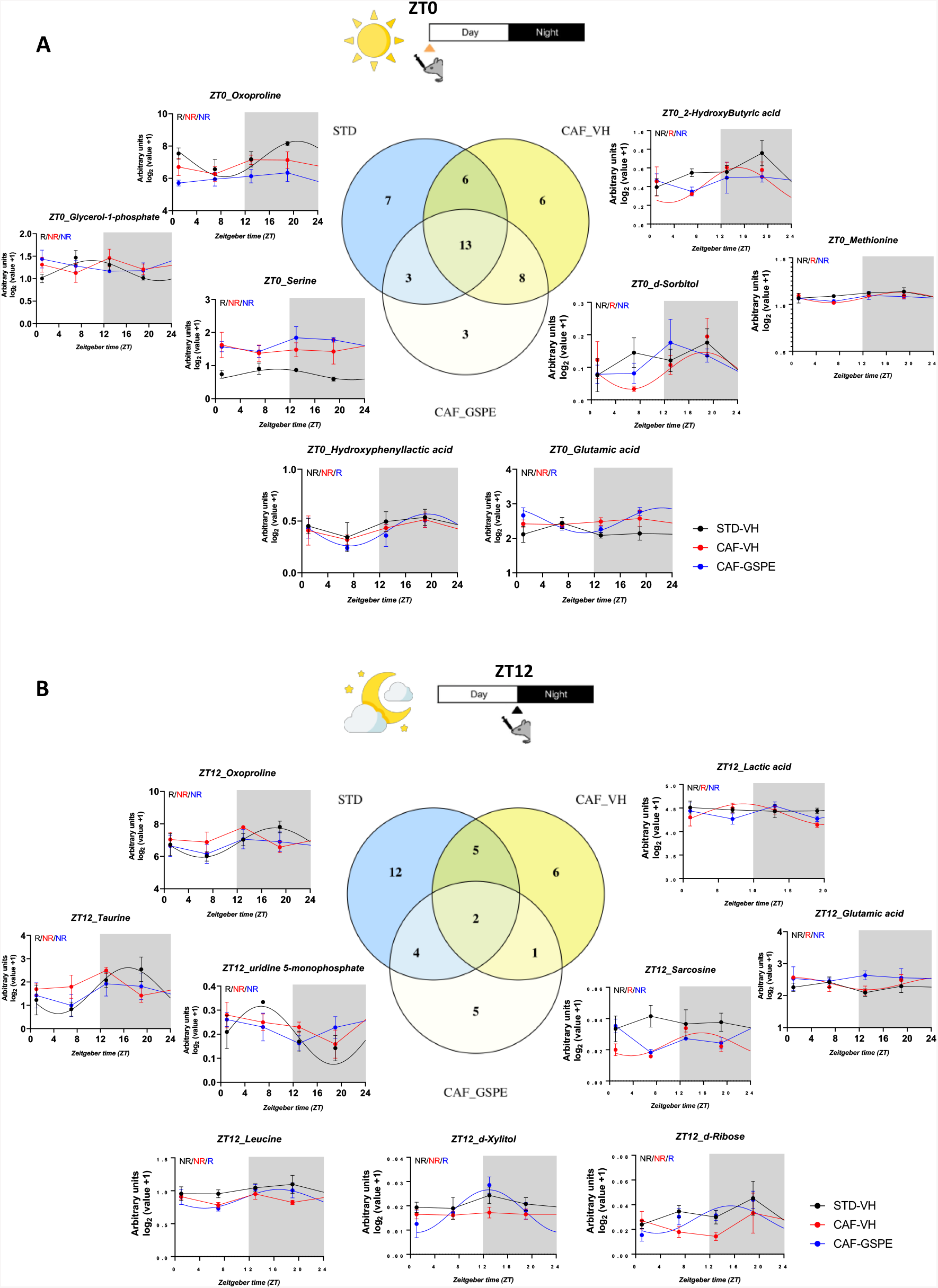
**M**etabolite rhythms under different diet/GSPE conditions. Rats were fed a STD or CAF diet and received a daily dosage of vehicle or GSPE at the beginning of the light phase (ZT0) **(A)** or at the beginning of the dark phase (ZT12) **(B)**. Venn diagrams depict numbers of rhythmic metabolites under each condition. Profiles show exemplary metabolites. Rhythm parameter determination and comparison were performed using CircaCompare. R/NR indicates significant/non-significant rhythmicity (p < 0.05). Shown are means ± SEM (n=4/group at each time point).

**Figure 8.**
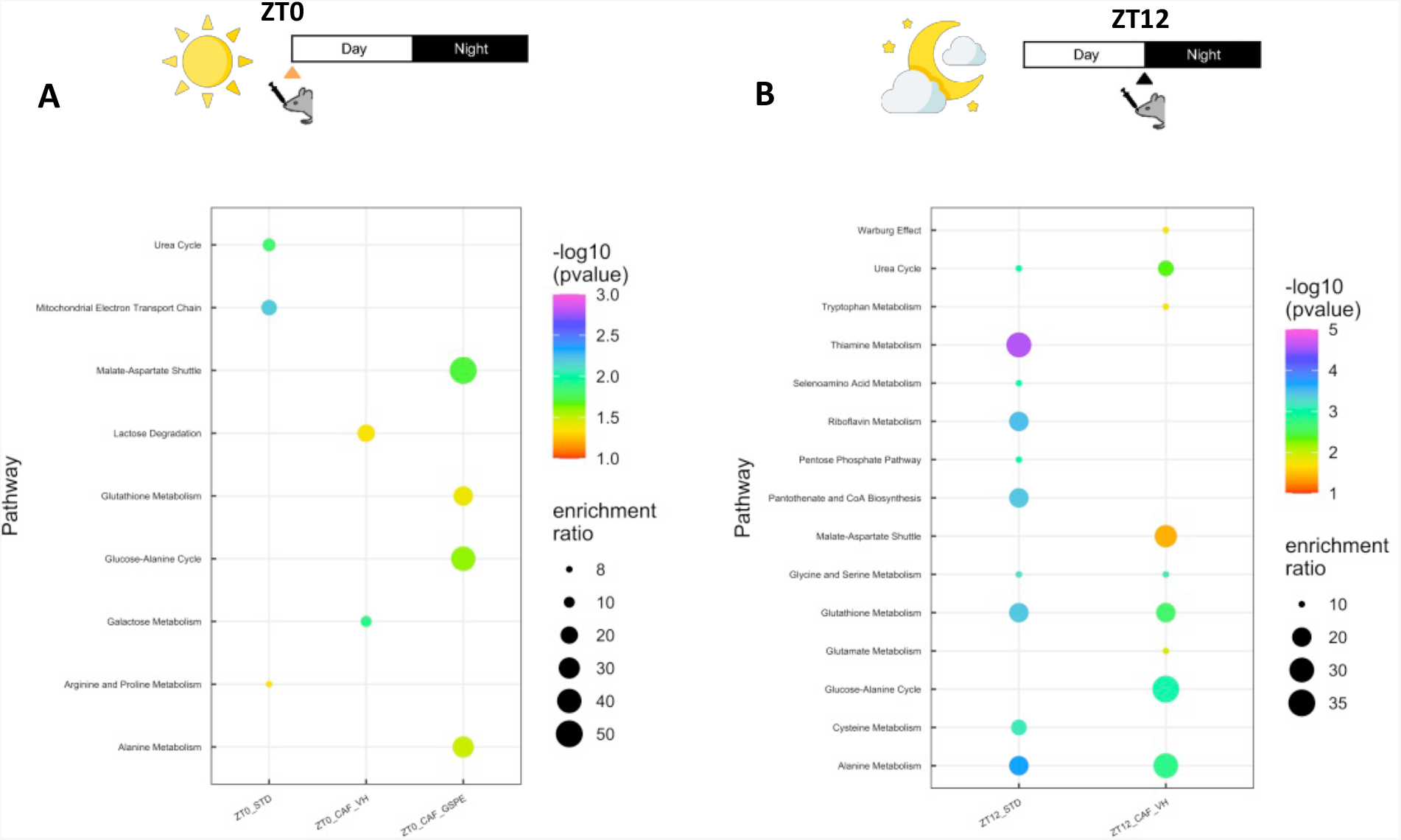
Metabolic pathways enrichment analysis of metabolites that showed an exclusive rhythmic expression in each group. Rats were fed a STD or CAF diet and received a daily dosage of vehicle or GSPE at the beginning of the light phase (ZT0) **(A)** or at the beginning of the dark phase (ZT12) **(B)**. The analysis was performed using MetaboAnalyst (Version 5.0, URL: http://www.metaboanalyst.ca).

In addition, we focused on the metabolites that lost rhythmicity in the CAF-VH group compared to STD-VH and observed that in the morning treatment (ZT0) oscillations of several metabolites were affected due to CAF diet: urea, glycine, fumaric acid, serine, threonine, malic acid, oxoproline, 4-hydroxyproline, glycerol 1-phosphate, ornithine, adenosine, and adenosine-5-mono-(AMP), -di-(ADP), and -triphosphate (ATP). Interestingly, GSPE treatment restored diurnal oscillations of urea, glycine, threonine, and AMP, ADP, and ATP (Figure 9 A-D). A phase delay was observed in lactic acid, alanine, proline, nicotinamide, d-galactitol, myo-inositol, sedoheptulose, and pyruvic acid (tendency p = 0.055) metabolites in CAF-VH animals, whereas CAF-GSPE rats exhibit a phase delay only in proline, nicotinamide and glycine - both compared to STD-VH. Moreover, compared to CAF-VH animals treated with GSPE showed a 6-hour phase advance in α-ketoglutaric acid, an increase of MESOR for sarcosine, fumaric acid, α-ketoglutaric acid and d-glucuronic acid, and a MESOR decrease for oxoproline, taurine, fructose 6-phosphate and glucose 6-phosphate.

**Figure 9.**
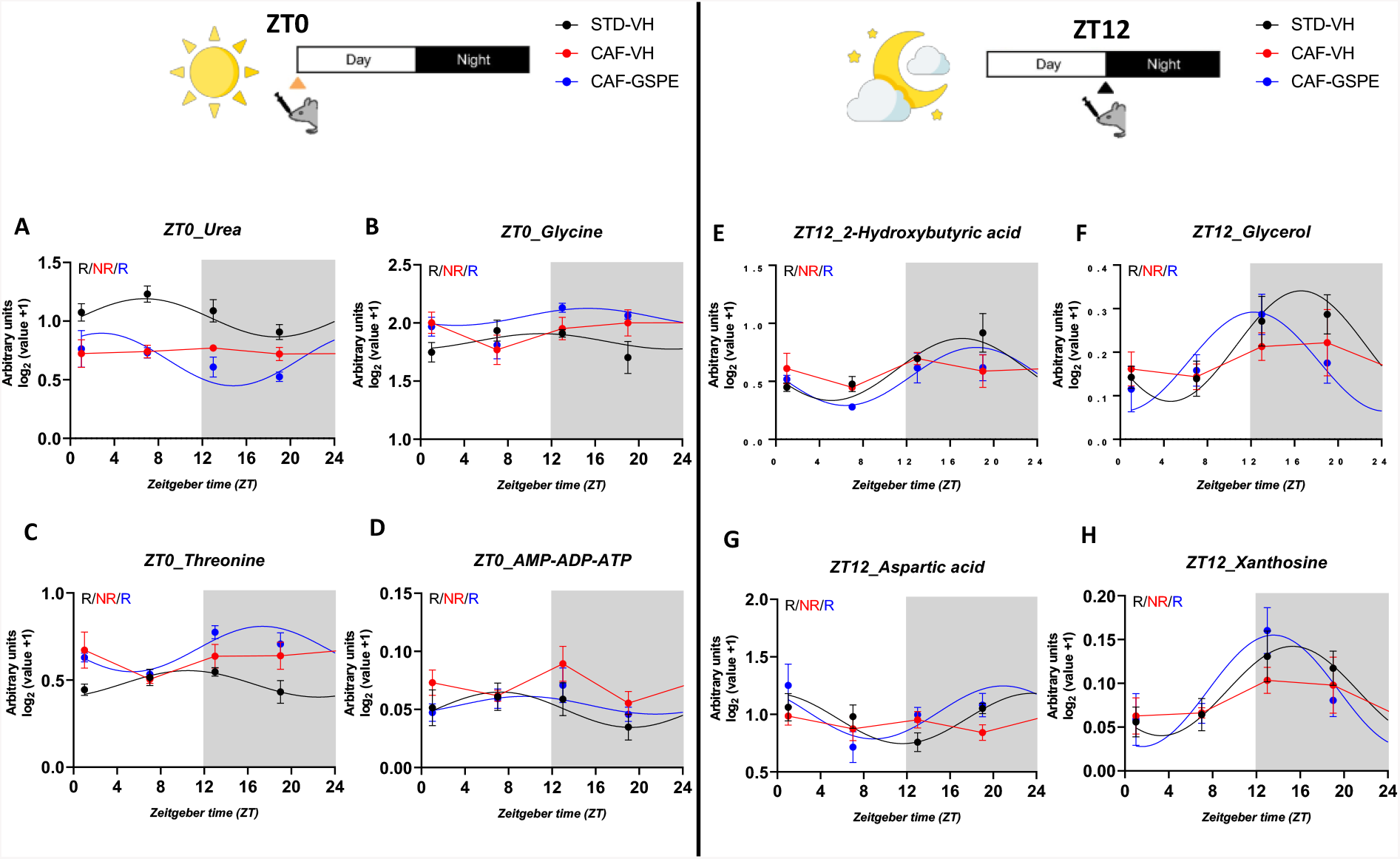
Restored diurnal oscillations of liver metabolites after chronic GSPE treatment. Rats were fed a STD or CAF diet and received a daily dosage of vehicle or GSPE at the beginning of the light phase (ZT0) **(A-D)** or at the beginning of the dark phase (ZT12) **(E-H)**. Rhythm parameter determination and comparison were performed using CircaCompare. R/NR indicates significant/non-significant rhythmicity (p < 0.05). Shown are means ± SEM (n=4/group at each time point).

In the case of the night treatment CAF diet caused loss of circadian oscillations in several metabolites including 2-hydroxybutyric acid, glycerol, succinic acid, serine, methionine, oxoproline, aspartic acid, taurine, citric acid, hydroxyphenyllactic acid, d-glucose, d-sorbitol, xanthosine, and inosine 5-monophosphate. Treatment with GSPE restored the rhythmic expression of 2-hydroxybutyric acid, glycerol, aspartic acid, and xanthosine (Figure 9 E-H) and tended to do the same for succinic, citric acid (p = 0.08), d-glucose (p = 0.055) and hydroxyphenyllactic acid (p=0.06). Moreover, the last-mentioned metabolite had a lower amplitude in CAF-VH but not in CAF-GSPE compared to STD-VH animals. When comparing CAF-VH and CAF-GSPE groups, animals that were treated with GSPE showed an increase in serine MESOR (p = 0.03) and a trend to increase in glycine MESOR (p = 0.05), a phase advance in adenine and malic acid, and higher amplitude in d-xylitol.

### 3.7. (+)-Catechin is one of the main GSPE phenolic components involved in its hepatoprotective effect against NAFLD

Predominant phenolic compounds in GSPE as determined by HPLC analysis included (+)-catechin and (-)-epicatechin. On the background of GSPE treatment’s restorative effect on mRNA and metabolite rhythms in liver, and knowing that these phenolic compounds are highly bioavailable, we decided to explore the capacity of these two polyphenols to affect circadian rhythm regulation in an *in vitro* model of NAFLD [55]. AML12 cells stably expressing a circadian *Bmal1-Luc* reporter were cultivated with 0.25 mM palmitate (to induce steatosis) or BSA and then treated at different concentrations (10 and 100 μM) with catechin or epicatechin. Circadian rhythms were determined by real-time luminescence measurements and responses to treatment analyzed for effects on rhythm parameters, i.e., phase, amplitude, period and dampening.

Cells cultured with palmitate showed a reduction of *Bmal1* amplitude, a phase shift and increased period compared to BSA cultured control cells (p < 0.001; Figure 10 A-D). Whereas treatment with (-)-epicatechin could not improve these circadian disturbances, cells treated with (+)-catechin exhibit a restoration of the amplitude, with an amelioration of the decreasing amplitude produced by palmitate treatment. Similar results were seen in acrophase, as only (+)-catechin treatments could restore the phase delay caused by palmitate (Figure 10 C). In addition, the lengthening of the period observed in palmitate and (-)-epicatechin treated-cells (p < 0.001) was not observed in cells treated with (+)-catechin (Figure 10 D). Based on these results, we analyzed the expression of glucose and lipid metabolism associated genes as mediators of the circadian regulation of metabolic control. The first step of glycolysis is catalyzed by glucokinase (enconded by *Gk)*. As is shown in Figure 10 F an increase of *Gk* expression was observed in cells treated with (+)-catechin (100 μM) compared to palmitate and with cells treated with (-)-epicatechin (100 μM). Moreover, expression of *Gys2* (glycogen synthase 2), which encodes the rate-limiting enzyme of glycogen storage in the liver, was downregulated due to palmitate, whereas treatment with (+)-catechin 100 μM reverted this decrease in *Gys2* expression (Figure 10 G). A downregulation of *G6pc* expression in palmitate-treated cells was observed, whereas this was not seen in cells treated with polyphenol extracts at the highest concentration (100 μM) (Figure 10 H). Regarding genes involved in lipid metabolism, a decrease in *Acaca* expression as well as in *Fasn* was observed in palmitate-treated cells, whereas cells treated with (+)-catechin did not show this downregulation in *Acaca* expression (Figure 10 I-J). Non-significant differences were seen for *Pfkl* (phosphofructokinase) and *Pck1* (phosphoenolpyruvate carboxykinase 1) (Figure 10 K-L).

**Figure 10.**
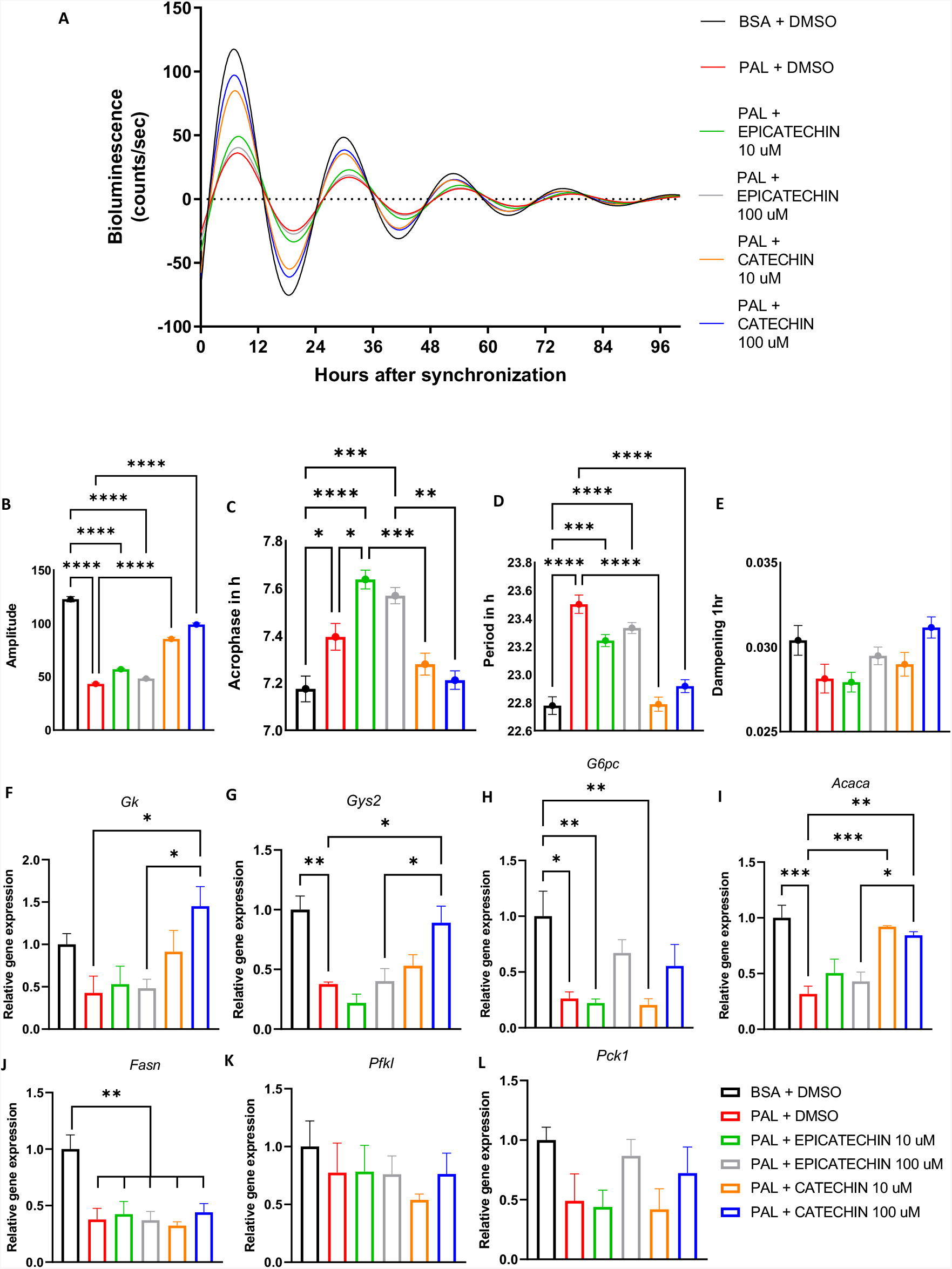
Effects of palmitate/catechin or epicatechin treatment on circadian clock rhythms in hepatocytes *in vitro*. **(A)** Luminescence rhythms in synchronized AML12 hepatocytes stably expressing *Bmal1-luc* reporter and after treatment with GSPE. Representative dampened sine curve fits on normalized bioluminescence data are shown. **(B-E)** Circadian rhythm parameters of AML12 *Bmal1-luc* cells (amplitude, acrophase, period, and dampening) in response to palmitate/GSPE treatment. Cells were treated with catechin or epicatechin (10 μM or 100 μM) for 96 h and cultured with sterile bovine serum albumin (BSA) conjugated with palmitate (PAL) (0.25 mM) with 0.1 % (v/v) DMSO. Cells cultured in BSA (0.04 mM) with 0.1 % (v/v) DMSO were used as control. Bioluminescence was measured using LumiCycle incubator in 10-min intervals. Rhythm parameters were assessed on normalized bioluminescence data using CircaSingle algorithm. **(F-L)** mRNA expression levels of metabolic genes in AML12/*Bmal1-luc* cells treated with catechin or epicatechin (10 μM or 100 μM) for 96 h and cultured with sterile bovine serum albumin (BSA) conjugated with palmitate (PAL) (0.25 mM) with 0.1 % (v/v) DMSO. Cells cultured in BSA (0.04 mM) with 0.1 % (v/v) DMSO were used as control. Data are means ± SEM (n = 4). One-way ANOVA followed by Tukey’s post-test was performed to compare values between groups (*p<0.05, **p<0.01, ***p<0.001).

## 4. DISCUSSION

The liver circadian clock plays an important role in regulating glucose, lipid, and bile acid metabolism, by controlling the expression of key metabolic genes implicated in these pathways. In fact, the pathophysiology of NAFLD involves dysfunction of these regulatory pathways. Moreover, genes that control the circadian clock have been implicated with the progression of NAFLD [56]. Therefore, the aim of the present study was to investigate the hepatic diurnal rhythm of lipid and glucose metabolism in cafeteria diet induced obese rats and to observe the effects of GSPE administration considering the timing of administration. Furthermore, we wanted to investigate the role of (+)-catechin and (-)-epicatechin, as principal polyphenols’ components of GSPE, in its hepatic metabolic circadian regulation.

CLOCK-BMAL1 heterodimer, the main players of the circadian clock machinery, contributes to modulate daily lipid metabolism in the liver by influencing SREBP-1c, and its downstream targets such as *Fasn* and *Acaca* [57]. Moreover, several studies involving liver specific depletion of clock genes have demonstrated the influence of circadian rhythms on liver energy metabolism [58]. By knocking out *Rorα* in the liver of mice, Zhang and collaborators demonstrated that this clock gene controls lipid metabolism by regulating expression of *Fasn, Elovl6, Acaca*, and SREBP-1c [59]. *Per2* knockout mice developed hepatic fibrosis with activation of hepatic stellate cells in the carbon tetrachloride-induced hepatitis model [60]. In addition, *Per* or *Cry* knockout mice can develop hepatocellular carcinoma [61]. In this sense, we have previously reported that CAF diet impairs the rhythmic expression pattern of hepatic core-clock genes, including *Bmal1, Per2, Cry1*, and *Rorα*, and that treatment with GSPE can restore the circadian oscillation of these clock genes mainly by correcting the phase shift caused by the CAF diet, with more emphasis at ZT12 treatment [36]. Remarkably our present results show that many genes encoding components of the lipogenic pathway, including *Acaca, Fasn*, SREBP-1c and *Cd36* display impaired or lost circadian expression rhythms in animals fed a CAF diet. However, when GSPE treatment was administered rats showed a robust diurnal expression pattern of these genes, conceivably improving lipid metabolism in the liver by restoring clock genes diurnal oscillations. The present study confirmed that the CAF diet induced misalignment of the oscillations of lipid metabolism– related genes, which could lead to enhance the *de novo* lipogenesis. This correlates with studies of liver-specific *Bmal1* and *Rorα* knock-out mice which exhibit increased liver triglyceride accumulation [59,62]. Our results showed that CAF animals that were administered with GSPE at ZT12 had lower levels of hepatic triglycerides and decreased lipid droplets than those who were administered with the VH at the same ZT. Moreover, our *in vitro* results point to (+)-catechin as one of the main GSPE components involved in hepatic clock regulation, increasing amplitude of *Bmal1*, restoring the phase delay and period lengthening, and improving expression of glucose and lipid related metabolic genes in cells exposed to palmitate treatment. Antioxidant and anti-inflammatory properties of (+)-catechin treatment have been evidenced by *in vitro* studies as it influences gene expression of Nrf2 and NF-kB inflammatory pathways and regulates key enzymes involved in oxidative stress, showing hepatoprotective effects against liver injury [63–65]. It was also shown that (+)-catechin exerts anti-obesogenic properties in 3T3-L1 adipocytes by inhibiting 3T3-L1 preadipocyte differentiation via modulating the C/EBP/PPARγ/SREBP-1c pathway. Moreover, (+)-catechin stimulated lipid degradation of mature adipocytes through cAMP/PKA pathways thereby exerting anti-adipogenic effects [66]. It has been shown that palmitate inhibits SIRT1-dependent BMAL1/CLOCK interaction and alters circadian gene oscillations in hepatocytes [67]. Interestingly, (+)-catechin has been shown to stimulate *Sirt1* expression [68] but not (-)-epicatechin [69]. Thus, based on the *in vitro* results, it could be possible that (+)-catechin by modulating *Sirt1* expression could be improving the hepatic circadian rhythmicity of metabolic pathways that promote fatty acid oxidation and inhibit lipogenesis, reducing lipid accumulation in the liver. Nevertheless, further experiments are needed to validate this hypothesis.

Sinturel and collaborators observed diurnal oscillations in liver mass and hepatocyte size in mice, the amplitude of which depends on both feeding-fasting and light-dark cycles [70]. They discover that ribosome number, and protein content follow a daily rhythm, and that rRNA polyadenylation cycles are antiphasic to ribosomal protein synthesis oscillations. In this regard, in the present study animals treated at ZT0 showed a robust daily oscillation of liver mass in all groups, nonetheless CAF-VH animals exhibit a 6-hour phase delay compared to STD-VH. It is remarkable that treatment with GSPE is able to restore this phase shift caused by the CAF diet. Moreover, in animals treated at ZT12, CAF-VH rats completely lost rhythmicity of liver weight compared to STD, while treatment with GSPE restores hepatic mass oscillation albeit with a phase delay. Since the liver is crucial for metabolism and xenobiotic detoxification, an accurate mass oscillation is needed to ensure the proper execution of these processes [71].

Several recent studies have demonstrated a close relationship between metabolites and the circadian clock machinery [72]. Results from transcriptome and metabolome studies suggest circadian clocks in peripheral tissues have a major role in the temporal coordination of food processing [73,74]. In this sense, our study identified several metabolites that were significantly different in diet-induced obese rats compared to the control lean group. One of the most strongly affected metabolites was urea together with some urea cycle intermediates which include ornithine and aspartic acid. Several studies demonstrated an association between liver injury in NAFLD and impairment of urea biosynthesis [75–77]. In this sense, all these studies exhibit a downregulation of the urea cycle components altering the functional capacity for ureagenesis. Our results showed not only a decrease in urea levels in animals fed a CAF diet but a loss of rhythmicity on hepatic urea, ornithine, and aspartic acid. Treatment with GSPE was able to restore the diurnal oscillation of urea in ZT0 and of aspartic acid in ZT12. In addition, it has been shown that alterations in metabolites of the urea cycle act as a sensor of hepatocyte mitochondrial damage [78]. We have previously reported that the CAF diet causes dysfunction of mitochondrial dynamics by enhancing mitochondrial fission [36]. In addition, diurnal oscillations of intermediates of the TCA cycle such as pyruvic acid, succinic acid, fumaric acid, malic acid, a-ketoglutaric acid, and citric acid were lost or phase shifted in CAF fed animals. The partial restoration of TCA cycle circadian rhythmicity by GSPE supplementation may have beneficial effects on hepatic metabolic homeostasis, as mitochondria play a key role in energy metabolism. Moreover, it is known that glycine, serine and threonine can enter into and promote TCA cycle activity by converting into acetyl-CoA [79]. In this sense, our results show that glycine and its precursor threonine lose their diurnal rhythms in CAF fed rats while GSPE treatment can reestablish a daily oscillation despite a delay in phase when compared to STD-VH group in the morning treatment (ZT0). Moreover, reduced concentration of these two glucogenic amino acids, glycine and serine, have been observed in diabetic mice [80].

In insulin-resistant cells, glucose uptake is impaired, triggering hepatic gluconeogenesis and consuming these glucogenic amino acids to initiate the production of the glucose precursors pyruvate and 3-phosphoglycerate [81]. This evidence goes in line with our results as we observed a decrease in MESOR of glycine and serine in ZT12-CAF-VH animals compared to CAF-GSPE rats. In addition, concentrations of serine were significantly increased in animals treated with GSPE at ZT12, although rhythmicity of this amino acid could not be restored by GSPE treatment. However, GSPE treatment at ZT12 showed the greatest ability to restore the rhythm of metabolites affected by the CAF diet. In this sense diurnal changes of glycerol, of 2-hydroxybutyric acid, aspartic acid, and xanthosine were restored with no differences in phase or amplitude regarding the STD-VH group. It has been shown that NAFLD disrupts glycerol metabolism affecting the insulin resistant state [82]. Moreover, liver steatosis was linked to increased glycerol metabolism via pathways intersecting with the TCA cycle and delayed gluconeogenesis from glycerol [83]. Therefore, glycerol is a pivotal substrate for hepatic glucose synthesis. Besides, a trend to reestablish d-glucose oscillations is seen in rats treated with GSPE at ZT12. This result correlates with mRNA circadian expression of *Slc2a2* which codifies the glucose transporter GLUT2, a crucial protein for glucose flux in hepatocytes. *Slc2a2* mRNA was phase delayed in CAF-VH but not in CAF-GSPE animals after ZT12 dosage. 2-hydroxybutyric acid has been reported as an early biomarker of insulin resistance and impaired glucose balance [84]. Furthermore, it was identified as an exercise-induced metabolite that may reflect the cytosolic redox state and energy stress at a systemic level. Another study supports this concept showing that elevated levels of this metabolite are associated with excess glutathione demand and disrupted mitochondrial energy metabolism, leading to increased oxidative stress in insulin resistance associated states [85]. Furthermore, aspartic acid also plays a role in energy production in the body as an intermediate in the urea cycle and is involved in gluconeogenesis [86]. In this sense, there is a close relationship between ureagenesis and gluconeogenesis [87] and both processes have been seen to be affected by the obesogenic CAF diet intake. Moreover, glutathione metabolism was also observed to be compromised in CAF-VH groups by the metabolic pathway enrichment analysis. In addition to energy dysregulation, the levels of metabolites involved in purine metabolism, including xanthine and its precursor xanthosine are upregulated in NAFLD subjects, human patients, and rats, which might accumulate reactive oxygen species (ROS) in the liver promoting hepatic steatosis [88–90], showing a correlation with the histological results. The mechanism underlying may involve stimulation of superoxide radical production through the hypoxanthine-xanthine oxidase system associated with various ROS-related diseases [91,92]. Hydroxyphenyllactic acid, which is a derivative of tyrosine, can enhance oxidative stress by promoting the production of ROS in the mitochondria [93]. Indeed, a dysregulation in circadian oscillation and amplitude of hydroxyphenyllactic acid was observed in CAF-VH fed rats but not in animals treated with GSPE at ZT12.

Energy for lipogenesis is provided by NADPH, which is primarily produced by the pentose phosphate pathway (PPP). A loss of G6PD rhythmic expression due to CAF diet was observed in both day and night treatment groups. This gene encodes for a cytosolic enzyme that catalyzes the rate-limiting step of the oxidative PPP and whose principal function is to provide NADPH and pentose phosphates for fatty acid and nucleic acid synthesis [51]. In this sense, GSPE treatment at ZT0 and ZT12 is able to restore these circadian disruptions, although a decrease in MESOR was observed in the morning treatment compared to STD group. Moreover, in our previous study we had observed that circadian oscillations of nicotinamide phosphoribosyltransferase (*Nampt*), which encodes for rate limiting component in the mammalian NAD biosynthesis pathway, exhibit a 3-hour phase advance in CAF-VH group in comparison to STD, meanwhile this difference had not been seen when CAF animals were treated with GSPE at ZT12 [36]. Remarkably, also at night treatment nicotinamide levels were lower in CAF-VH but not in CAF-GSPE animals compared to STD-VH rats. This suggests that energy production through NAD-related metabolism and enzymes may be impaired in CAF diet obese rats, resulting in the accumulation of TAGs in the liver and a dysregulation of energy balance. These findings are consistent with previous studies in the field of proanthocyanidins treatment. An acute dose of 250 mg/kg of GSPE, ten times the dose used in the present study, was found to increase hepatic *Nampt* expression and NAD levels in the dark phase while the effect was the opposite when it was administered in the light phase in healthy rats [94]. Furthermore, a significant increase of hepatic NAD content was observed after a chronic treatment with GSPE in healthy rats [95]. GSPE treatment was shown to modulate hepatic concentrations of the major NAD precursors and mRNA levels of genes encoding enzymes involved in NAD metabolism together with the upregulation of *Sirt1* gene expression in a dose-dependent manner. In addition, the authors suggested that this increase in NAD concentrations and *Sirt1* mRNA levels is significantly associated with improved protection against hepatic TAG accumulation [95]. It is known that SIRT1 acts in the liver as a master metabolic regulator controlling the expression of downstream metabolic targets [96]. It forms a complex with the CLOCK-BMAL1 heterodimer binding to E-box elements promoting the expression of *Nampt*. Nevertheless, no effect on circadian oscillations with GSPE treatment were observed for *Sirt1*; but a clear rhythmic expression was seen in *Ppargc1α* after ZT0 and ZT12 GSPE treatments. In the night treatment cohort, *Ppargc1α* circadian oscillations in CAF-VH animals peaked at ZT11 whereas in STD-VH and CAF-GSPE animals the peak was at ZT13. It has been described that SIRT1 induces gluconeogenic genes and hepatic glucose output through PGC-1α [97]. Therefore, the intake of an obesogenic diet causes a dysregulation of glucose and lipid homeostasis in the liver, while a low dose of GSPE during the active phase (ZT12) ameliorates this disruption in a clock-related manner. The integrative results from hepatic lipid profiles, circadian gene expression of core clock and metabolic genes together with the metabolomic studies suggest that, in CAF diet rat livers, glucose metabolism is disrupted, and lipogenesis is enhanced, promoting fatty liver development. By restoring the oscillations of hepatic circadian clocks, liver mass, key lipogenic and glucogenic gene expression, and liver metabolites, ZT12-GSPE supplementation has liver-protecting properties and counteracts NAFLD development.

## 5. CONCLUSIONS

Our chrono-nutritional approach, in a rat model, showed that an intake of a CAF diet disturbed the hepatic circadian clock, lipid and glucose-related metabolic genes, as well as concentrations and oscillations of liver metabolites, ultimately promoting the development of NAFLD. Supplementation with GSPE ameliorates and partially corrects this disruption, with a stronger effect when administration is timed to the active phase (ZT12 treatment). This indicates that modulation of metabolic rhythms by proanthocyanidins is highly dependent on the time of administration. Furthermore, our *in vitro* results suggest (+)-catechin, one of the main phenolic compounds found in the GSPE extract, may be involved in the ameliorating effects of GSPE on NAFLD development. Our findings underline the importance of taking diurnal rhythms into account in the interpretation of metabolic studies and suggest that they should more generally be considered for therapeutic intervention targeting liver metabolism.

## ACKNOWLEDGMENTS

This research was funded by the Spanish Ministry of Economy and Competitiveness (MINECO), AGL2016-77105-R (CHRONOFOOD project), and by grants from the German Research Foundation (DFG): HO353-10/1, HO353-11/1 and CRC-296 (TP13). The authors would like to thank to Niurka Llópiz and Rosa Pastor (Tarragona), for their help and technical assistance.

