## Supplements for "Grape-seed proanthocyanidin extract (GSPE) modulates diurnal oscillations of key hepatic metabolic genes and metabolites alleviating hepatic lipid deposition in cafeteria-fed obese rats in a time-of-day-dependent manner"

### SUPPLEMENTARY MATERIAL

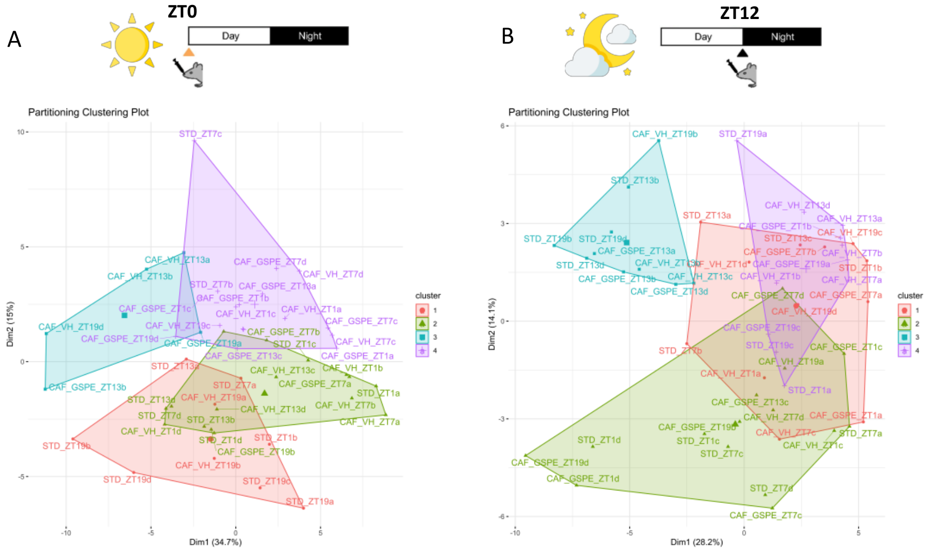

**Supplementary Figure 1.** Principal component analysis (PCA) of liver metabolome analysis. Rats were fed a STD or CAF diet and received a daily dosage of vehicle or GSPE at the beginning of the light phase (ZT0) (A) or at the beginning of the dark phase (ZT12) (B). Analyzes were performed using the factoextra package and Hartigan-Wong, Lloyd, and Forgy MacQueen algorithms (version 1.0.7) in R.

**Supplementary Table 1.** Nucleotide sequences of primers used for real-time quantitative PCR in liver tissue.

| Gene | Forward primer (5' to 3') | Reverse primer (5' to 3') |
|---|---|---|
| *Acacα* | TGCAGGTATCCCCACTCTTC | TTCTGATTCCCTTCCCTCCT |
| *Cd36* | GTCCTGGCTGTGTTTGGA | GCTCAAAGATGGCTCCATTG |
| *Cyp7a1* | CACTTGTTCAAGACCGCACA | TGCTTGAGATGCCCAGAGAA |
| *Fasn* | TAAGCGGTCTGGAAAGCTGA | CACCAGTGTTTGTTCCTCGG |
| *Fatp5* | CCTGCCAAGCTTCGTGCTAAT | GCTCATGTGATAGGATGGCTGG |
| *G6pc* | ATTCCGGTGCTTGAATGTCG | TGGAGGCTGGCATTGTAGAT |
| *G6pd* | ACCAGGCATTCAAAACGCAT | CAGTCTCAGGGAAGTGTGGT |
| *Gk* | CTGTGAAAGCGTGTCCACTC | GCCCTCCTCTGATTCGATGA |
| *Ppargc1α* | AGAGTCACCAAATGACCCCAAG | TTGGCTTTATGAGGAGGAGTCG |
| *Pparα* | CGGCGTTGAAAACAAGGAGG | TTGGGTTCCATGATGTCGCA |
| *Ppia* | CCAAACACAAATGGTTCCCAGT | ATTCCTGGACCCAAAACGCT |
| *Sirt1* | TTGGCACCGATCCTCGAA | ACAGAAACCCCAGCTCCA |
| *Slc2a2* | AGTCACACCAGCACATACGA | TGGCTTTGATCCTTCCGAGT |
| *Srebp1c* | CCCACCCCCTTACACACC | GCCTGCGGTCTTCATTGT |

**Supplementary Table 2.** Nucleotide sequences of primers used for real-time quantitative PCR in cells.

| Gene | Forward primer (5' to 3') | Reverse primer (5' to 3') |
|---|---|---|
| *Acacα* | GTCCCCAGGGATGAACCAATA | GCCATGCTCAACCAAAGTAGC |
| *eEf1a* | TGCCCCAGGACACAGAGACTTCA | AATTCACCAACACCAGCAGCAA |
| *Fasn* | GGAGGTGGTGATAGCCGGTAT | TGGGTAATCCATAGAGCCCAG |
| *G6pc* | CGACTCGCTATCTCCAAGTGA | GTTGAACCAGTCTCCGACCA |
| *Gk* | AGACCTGGGAGGAACCAACT | TTTGTCTTCACGCTCCACTG |
| *Gys2* | GCTCTCCAGACGTTCTTGCA | GTGCGGTTCCTCTGAATGATC |
| *Pck1* | AAGCATTCAACGCCAGGTTC | GGGCGAGTCTGTCAGTTCAAT |
| *Pfkl* | GAACTACGCACACTTGACCAT | CTCCAAAACAAAGGTCCTCTGG |

**Supplementary Table 3.** Rhythmic parameters of lipid liver profile

| Parameter | ZT | Groups | *p* | MESOR | Amplitude | Acrophase [h] |
|---|---|---|---|---|---|---|
| **Cholesterol** | ZT0 | STD-VH | 0.038 | 1.191 | 0.064 | 12.996 |
| | | CAF-VH | 0.734 | 1.388 | 0.021 | 3.527 |
| | | CAF-GSPE | 0.515 | 1.411 | 0.049 | 17.921 |
| | ZT12 | STD-VH | 0.034 | 1.123 | 0.064 | 19.007 |
| | | CAF-VH | 0.589 | 1.310 | 0.031 | 17.746 |
| | | CAF-GSPE | 0.681 | 1.238 | 0.019 | 11.628 |
| **Triglycerides** | ZT0 | STD-VH | 0.152 | 2.226 | 0.140 | 10.409 |
| | | CAF-VH | 0.085 | 3.497 | 0.324 | 6.704 |
| | | CAF-GSPE | 0.626 | 3.382 | 0.085 | 17.860 |
| | ZT12 | STD-VH | 0.455 | 2.141 | 0.148 | 1.642 |
| | | CAF-VH | 0.541 | 3.592 | 0.088 | 23.201 |
| | | CAF-GSPE | 0.422 | 3.307 | 0.123 | 7.382 |
| **Total lipid liver content** | ZT0 | STD-VH | 0.001 | 5.898 | 0.314 | 17.173 |
| | | CAF-VH | 0.572 | 6.404 | 0.067 | 11.436 |
| | | CAF-GSPE | 0.342 | 6.364 | 0.117 | 16.686 |
| | ZT12 | STD-VH | 0.105 | 6.034 | 0.156 | 19.857 |
| | | CAF-VH | 0.091 | 6.541 | 0.159 | 22.295 |
| | | CAF-GSPE | 0.221 | 6.357 | 0.088 | 17.305 |
| **Liver weight** | ZT0 | STD-VH | 0.016 | 4.086 | 0.145 | 9.360 |
| | | CAF-VH | 0.001 | 4.370 | 0.181 | 16.309 |
| | | CAF-GSPE | 0.019 | 4.271 | 0.225 | 10.607 |
| | ZT12 | STD-VH | 0.005 | 3.995 | 0.162 | 2.833 |
| | | CAF-VH | 0.289 | 4.403 | 0.078 | 2.726 |
| | | CAF-GSPE | 0.001 | 4.301 | 0.229 | 6.826 |

Rats were fed a STD or CAF diet and were treated with vehicle or GSPE at the beginning of the light phase (ZT0) or at the beginning of the dark phase (ZT12). The $p < 0.05$ indicates significant rhythmic expression. MESOR is a mean adjusted to the circadian rhythm. Amplitude is the difference between the peak and the mean value of a sine wave. Acrophase is the time at which the peak of a rhythm occurs in hours.

**Supplementary Table 4.** Comparison of rhythmic parameters of lipid liver profile between groups.

| Parameter | ZT | Comparison | d_MESOR | *p*(d_MESOR) | d_amplitude | *p*(d_amplitu |
|---|---|---|---|---|---|---|
| **Cholesterol** | ZT0 | STD-VH vs. CAF-VH | 0.197 | 0.000 | -0.043 | 0.520 |
| | | CAF-VH vs. CAF-GSPE | 0.023 | 0.740 | 0.028 | 0.768 |
| | | STD-VH vs. CAF-GSPE | 0.220 | 0.001 | -0.015 | 0.847 |
| | ZT12 | STD-VH vs. CAF-VH | 0.188 | 0.000 | -0.033 | 0.600 |
| | | CAF-VH vs. CAF-GSPE | -0.072 | 0.167 | -0.012 | 0.865 |
| | | STD-VH vs. CAF-GSPE | 0.116 | 0.004 | -0.045 | 0.394 |
| **Triglycerides** | ZT0 | STD-VH vs. CAF-VH | 1.271 | 0.000 | 0.184 | 0.359 |
| | | CAF-VH vs. CAF-GSPE | -0.115 | 0.528 | -0.239 | 0.337 |
| | | STD-VH vs. CAF-GSPE | 1.156 | 0.000 | -0.055 | 0.773 |
| | ZT12 | STD-VH vs. CAF-VH | 1.452 | 0.000 | -0.059 | 0.802 |
| | | CAF-VH vs. CAF-GSPE | -0.285 | 0.059 | 0.035 | 0.867 |
| | | STD-VH vs. CAF-GSPE | 1.167 | 0.000 | -0.025 | 0.918 |
| **Total lipid liver content** | ZT0 | STD-VH vs. CAF-VH | 0.506 | 0.000 | -0.248 | 0.082 |
| | | CAF-VH vs. CAF-GSPE | -0.040 | 0.731 | 0.051 | 0.763 |
| | | STD-VH vs. CAF-GSPE | 0.466 | 0.000 | -0.197 | 0.173 |
| | ZT12 | STD-VH vs. CAF-VH | 0.507 | 0.000 | 0.003 | 0.982 |
| | | CAF-VH vs. CAF-GSPE | -0.185 | 0.027 | -0.071 | 0.524 |
| | | STD-VH vs. CAF-GSPE | 0.323 | 0.001 | -0.068 | 0.554 |
| **Liver weight** | ZT0 | STD-VH vs. CAF-VH | 0.284 | 0.000 | 0.036 | 0.600 |
| | | CAF-VH vs. CAF-GSPE | -0.099 | 0.137 | 0.045 | 0.631 |
| | | STD-VH vs. CAF-GSPE | 0.185 | 0.011 | 0.081 | 0.410 |

| | | | | | | |
|---|---|---|---|---|---|---|
| | ZT12 | STD-VH vs. CAF-VH | 0.408 | 0.000 | -0.084 | 0.332 |
| | | CAF-VH vs. CAF-GSPE | -0.102 | 0.110 | 0.151 | 0.097 |
| | | STD-VH vs. CAF-GSPE | 0.306 | 0.000 | 0.067 | 0.352 |

Rats were fed a STD or CAF diet and were treated with vehicle or GSPE at the beginning of the light phase (ZT
(ZT12). d_MESOR represents the difference in MESOR values between the groups. d_amplitude represents the
the groups. d_phase represents the difference in acrophase values between the groups. The $p < 0.05$ indicates s
each rhythmic parameter.

**Supplementary Table 5.** Rhythmic parameters of hepatic lipid metabolic genes.

| Gene | ZT | Groups | *p* | MESOR | Amplitude | Acrophase [h] |
|---|---|---|---|---|---|---|
| ***Acaca*** | ZT0 | STD-VH | 0.146 | 0.933 | 0.117 | 8.075 |
| | | CAF-VH | 0.085 | 0.868 | 0.091 | 18.192 |
| | | CAF-GSPE | 0.002 | 0.844 | 0.240 | 23.118 |
| | ZT12 | STD-VH | 0.919 | 0.854 | 0.009 | 2.741 |
| | | CAF-VH | 0.012 | 0.869 | 0.198 | 19.014 |
| | | CAF-GSPE | 0.001 | 0.921 | 0.231 | 22.324 |
| ***Fasn*** | ZT0 | STD-VH | 0.040 | 0.686 | 0.248 | 3.677 |
| | | CAF-VH | 0.143 | 0.376 | 0.150 | 20.176 |
| | | CAF-GSPE | 0.010 | 0.366 | 0.262 | 23.384 |
| | ZT12 | STD-VH | 0.053 | 0.613 | 0.249 | 2.267 |
| | | CAF-VH | 0.001 | 0.530 | 0.389 | 21.007 |
| | | CAF-GSPE | 0.002 | 0.626 | 0.394 | 22.216 |
| ***Cd36*** | ZT0 | STD-VH | 0.060 | 0.745 | 0.180 | 1.224 |
| | | CAF-VH | 0.094 | 0.902 | 0.244 | 0.805 |
| | | CAF-GSPE | 0.011 | 0.834 | 0.241 | 21.407 |
| | ZT12 | STD-VH | 0.101 | 0.913 | 0.188 | 5.970 |
| | | CAF-VH | 0.141 | 0.926 | 0.186 | 3.233 |
| | | CAF-GSPE | 0.010 | 0.976 | 0.326 | 2.774 |
| ***Fatp5*** | ZT0 | STD-VH | 0.605 | 0.844 | 0.037 | 23.531 |
| | | CAF-VH | 0.017 | 1.013 | 0.155 | 18.612 |
| | | CAF-GSPE | 0.047 | 1.011 | 0.100 | 17.570 |
| | ZT12 | STD-VH | 0.011 | 0.944 | 0.074 | 20.654 |
| | | CAF-VH | 0.486 | 1.090 | 0.042 | 11.010 |
| | | CAF-GSPE | 0.259 | 1.206 | 0.074 | 2.086 |
| ***SREBP-1c*** | ZT0 | STD-VH | 0.010 | 0.623 | 0.288 | 3.502 |
| | | CAF-VH | 0.072 | 0.502 | 0.139 | 8.485 |
| | | CAF-GSPE | 0.001 | 0.502 | 0.203 | 3.507 |
| | ZT12 | STD-VH | 0.005 | 0.797 | 0.329 | 5.121 |
| | | CAF-VH | 0.017 | 1.228 | 0.244 | 6.684 |
| | | CAF-GSPE | 0.001 | 1.102 | 0.411 | 3.849 |
| ***Ppara*** | ZT0 | STD-VH | 0.004 | 1.145 | 0.293 | 11.474 |
| | | CAF-VH | 0.011 | 1.527 | 0.280 | 12.529 |
| | | CAF-GSPE | 0.047 | 1.449 | 0.210 | 15.968 |
| | ZT12 | STD-VH | 0.000 | 1.493 | 0.595 | 13.118 |
| | | CAF-VH | 0.002 | 1.613 | 0.407 | 12.296 |
| | | CAF-GSPE | 0.000 | 1.574 | 0.230 | 12.746 |
| ***Cyp7a1*** | ZT0 | STD-VH | 0.538 | 0.822 | 0.090 | 14.157 |

| | | | | | | |
|---|---|---|---|---|---|---|
| | | CAF-VH | 0.000 | 0.443 | 0.270 | 11.531 |
| | | CAF-GSPE | 0.002 | 0.332 | 0.216 | 12.615 |
| | ZT12 | STD-VH | 0.001 | 1.413 | 0.708 | 14.700 |
| | | CAF-VH | 0.000 | 1.249 | 0.939 | 11.799 |
| | | CAF-GSPE | 0.048 | 1.426 | 0.406 | 9.842 |

Rats were fed a STD or CAF diet and were treated with vehicle or GSPE at the beginning of the light phase (ZT0) or at the beginning of the dark phase (ZT12). The $p < 0.05$ indicates significant rhythmic expression. MESOR is a mean adjusted to the circadian rhythm. Amplitude is the difference between the peak and the mean value of a sine wave. Acrophase is the time at which the peak of a rhythm occurs in hours.

**Supplementary Table 6.** Comparison of rhythmic parameters of hepatic lipid metabolic genes between groups.

| Gene | ZT | Comparison | d_MESOR | p(d_MESOR) | d_amplitude | p(d_amp[illegible] |
|---|---|---|---|---|---|---|
| ***Acaca*** | ZT0 | STD-VH vs. CAF-VH | -0.065 | 0.317 | -0.026 | 0.77[illegible] |
| | | CAF-VH vs. CAF-GSPE | -0.024 | 0.668 | 0.149 | 0.06[illegible] |
| | | STD-VH vs. CAF-GSPE | -0.089 | 0.212 | 0.123 | 0.22[illegible] |
| | ZT12 | STD-VH vs. CAF-VH | 0.015 | 0.854 | 0.189 | 0.11[illegible] |
| | | CAF-VH vs. CAF-GSPE | 0.052 | 0.416 | 0.033 | 0.71[illegible] |
| | | STD-VH vs. CAF-GSPE | 0.067 | 0.387 | 0.222 | 0.04[illegible] |
| ***Fasn*** | ZT0 | STD-VH vs. CAF-VH | -0.309 | 0.006 | -0.098 | 0.50[illegible] |
| | | CAF-VH vs. CAF-GSPE | -0.011 | 0.909 | 0.112 | 0.39[illegible] |
| | | STD-VH vs. CAF-GSPE | -0.320 | 0.003 | 0.014 | 0.92[illegible] |
| | ZT12 | STD-VH vs. CAF-VH | -0.083 | 0.428 | 0.140 | 0.34[illegible] |
| | | CAF-VH vs. CAF-GSPE | 0.097 | 0.311 | 0.005 | 0.97[illegible] |
| | | STD-VH vs. CAF-GSPE | 0.014 | 0.901 | 0.145 | 0.35[illegible] |
| ***Cd36*** | ZT0 | STD-VH vs. CAF-VH | 0.157 | 0.181 | 0.064 | 0.70[illegible] |
| | | CAF-VH vs. CAF-GSPE | -0.069 | 0.550 | -0.004 | 0.98[illegible] |
| | | STD-VH vs. CAF-GSPE | 0.089 | 0.293 | 0.060 | 0.61[illegible] |
| | ZT12 | STD-VH vs. CAF-VH | 0.013 | 0.907 | -0.002 | 0.99[illegible] |
| | | CAF-VH vs. CAF-GSPE | 0.050 | 0.667 | 0.140 | 0.39[illegible] |
| | | STD-VH vs. CAF-GSPE | 0.063 | 0.563 | 0.138 | 0.37[illegible] |
| ***Fatp5*** | ZT0 | STD-VH vs. CAF-VH | 0.169 | 0.013 | 0.118 | 0.19[illegible] |
| | | CAF-VH vs. CAF-GSPE | -0.001 | 0.979 | -0.055 | 0.46[illegible] |

| | | | | | | |
|---|---|---|---|---|---|---|
| | | STD-VH vs. CAF-GSPE | 0.167 | 0.011 | 0.063 | 0.45 |
| | ZT12 | STD-VH vs. CAF-VH | 0.146 | 0.003 | -0.032 | 0.61 |
| | | CAF-VH vs. CAF-GSPE | 0.116 | 0.065 | 0.032 | 0.71 |
| | | STD-VH vs. CAF-GSPE | 0.262 | 0.000 | 0.000 | 1.00 |
| ***SREBP-1c*** | ZT0 | STD-VH vs. CAF-VH | -0.121 | 0.162 | -0.149 | 0.21 |
| | | CAF-VH vs. CAF-GSPE | 0.000 | 0.994 | 0.064 | 0.45 |
| | | STD-VH vs. CAF-GSPE | -0.121 | 0.123 | -0.085 | 0.43 |
| | ZT12 | STD-VH vs. CAF-VH | 0.431 | 0.000 | -0.085 | 0.52 |
| | | CAF-VH vs. CAF-GSPE | -0.126 | 0.182 | 0.168 | 0.20 |
| | | STD-VH vs. CAF-GSPE | 0.305 | 0.004 | 0.083 | 0.54 |
| ***Ppara*** | ZT0 | STD-VH vs. CAF-VH | 0.383 | 0.000 | -0.013 | 0.92 |
| | | CAF-VH vs. CAF-GSPE | -0.078 | 0.420 | -0.071 | 0.60 |
| | | STD-VH vs. CAF-GSPE | 0.305 | 0.002 | -0.083 | 0.51 |
| | ZT12 | STD-VH vs. CAF-VH | 0.120 | 0.228 | -0.189 | 0.18 |
| | | CAF-VH vs. CAF-GSPE | -0.040 | 0.634 | -0.176 | 0.14 |
| | | STD-VH vs. CAF-GSPE | 0.081 | 0.272 | -0.365 | 0.00 |
| ***Cyp7a1*** | ZT0 | STD-VH vs. CAF-VH | -0.379 | 0.001 | 0.180 | 0.24 |
| | | CAF-VH vs. CAF-GSPE | -0.111 | 0.037 | -0.054 | 0.46 |
| | | STD-VH vs. CAF-GSPE | -0.490 | 0.000 | 0.126 | 0.43 |
| | ZT12 | STD-VH vs. CAF-VH | -0.164 | 0.337 | 0.231 | 0.34 |
| | | CAF-VH vs. CAF-GSPE | 0.178 | 0.325 | -0.533 | 0.04 |
| | | STD-VH vs. CAF-GSPE | 0.013 | 0.941 | -0.302 | 0.24 |

**Supplementary Table 7.** Rhythmic parameters of genes related to hepatic glucose metabolism.

| Gene | ZT | Groups | *p* | MESOR | Amplitude | Acrophase [h] |
|---|---|---|---|---|---|---|
| ***G6pc*** | ZT0 | STD-VH | 0.052 | 0.810 | 0.276 | 2.718 |
| | | CAF-VH | 0.023 | 0.543 | 0.232 | 22.878 |
| | | CAF-GSPE | 0.032 | 0.536 | 0.213 | 1.868 |
| | ZT12 | STD-VH | 0.046 | 0.832 | 0.236 | 21.769 |
| | | CAF-VH | 0.002 | 0.597 | 0.290 | 0.153 |
| | | CAF-GSPE | 0.000 | 0.610 | 0.348 | 23.346 |
| ***G6pd*** | ZT0 | STD-VH | 0.002 | 0.763 | 0.318 | 4.938 |
| | | CAF-VH | 0.121 | 0.456 | 0.082 | 9.043 |
| | | CAF-GSPE | 0.040 | 0.451 | 0.140 | 1.729 |
| | ZT12 | STD-VH | 0.051 | 0.641 | 0.260 | 3.993 |
| | | CAF-VH | 0.374 | 0.470 | 0.075 | 4.536 |
| | | CAF-GSPE | 0.011 | 0.540 | 0.253 | 2.432 |
| ***Slc2a2*** | ZT0 | STD-VH | 0.035 | 1.245 | 0.320 | 12.255 |
| | | CAF-VH | 0.002 | 1.544 | 0.472 | 15.258 |
| | | CAF-GSPE | 0.002 | 1.229 | 0.305 | 13.562 |
| | ZT12 | STD-VH | 0.000 | 1.265 | 0.381 | 12.511 |
| | | CAF-VH | 0.000 | 1.332 | 0.397 | 14.920 |
| | | CAF-GSPE | 0.026 | 1.351 | 0.189 | 13.884 |
| ***Ppargc1α*** | ZT0 | STD-VH | 0.229 | 0.928 | 0.119 | 15.293 |
| | | CAF-VH | 0.106 | 0.754 | 0.139 | 12.805 |
| | | CAF-GSPE | 0.045 | 0.718 | 0.113 | 16.884 |
| | ZT12 | STD-VH | 0.003 | 1.333 | 0.476 | 13.748 |
| | | CAF-VH | 0.048 | 1.125 | 0.200 | 11.573 |
| | | CAF-GSPE | 0.015 | 1.126 | 0.251 | 13.819 |
| ***Sirt1*** | ZT0 | STD-VH | 0.280 | 0.952 | 0.070 | 9.612 |
| | | CAF-VH | 0.017 | 1.124 | 0.114 | 17.910 |
| | | CAF-GSPE | 0.042 | 1.094 | 0.107 | 19.720 |
| | ZT12 | STD-VH | 0.018 | 1.103 | 0.176 | 14.092 |
| | | CAF-VH | 0.107 | 1.184 | 0.054 | 16.010 |
| | | CAF-GSPE | 0.123 | 1.111 | 0.061 | 16.911 |

Rats were fed a STD or CAF diet and were treated with vehicle or GSPE at the beginning of the light phase (ZT0) or at the beginning of the dark phase (ZT12). The $p < 0.05$ indicates significant rhythmic expression. MESOR is a mean adjusted to the circadian rhythm. Amplitude is the difference between the peak and the mean value of a sine wave. Acrophase is the time at which the peak of a rhythm occurs in hours.

**Supplementary Table 8.** Comparison of rhythmic parameters of genes related to hepatic glucose metabolism b[illegible]

| Gene | ZT | Comparison | d_MESOR | p(d_MESOR) | d_amplitude | p(d_am[illegible] |
|---|---|---|---|---|---|---|
| ***G6pc*** | ZT0 | STD-VH vs. CAF-VH | -0.267 | 0.024 | -0.045 | 0.7[illegible] |
| | | CAF-VH vs. CAF-GSPE | -0.007 | 0.937 | -0.019 | 0.8[illegible] |
| | | STD-VH vs. CAF-GSPE | -0.274 | 0.021 | -0.064 | 0.6[illegible] |
| | ZT12 | STD-VH vs. CAF-VH | -0.235 | 0.018 | 0.053 | 0.6[illegible] |
| | | CAF-VH vs. CAF-GSPE | 0.013 | 0.859 | 0.058 | 0.5[illegible] |
| | | STD-VH vs. CAF-GSPE | -0.222 | 0.020 | 0.111 | 0.3[illegible] |
| ***G6pd*** | ZT0 | STD-VH vs. CAF-VH | -0.307 | 0.000 | -0.236 | 0.0[illegible] |
| | | CAF-VH vs. CAF-GSPE | -0.005 | 0.924 | 0.059 | 0.4[illegible] |
| | | STD-VH vs. CAF-GSPE | -0.312 | 0.000 | -0.177 | 0.1[illegible] |
| | ZT12 | STD-VH vs. CAF-VH | -0.171 | 0.109 | -0.185 | 0.2[illegible] |
| | | CAF-VH vs. CAF-GSPE | 0.070 | 0.409 | 0.178 | 0.1[illegible] |
| | | STD-VH vs. CAF-GSPE | -0.101 | 0.345 | -0.006 | 0.9[illegible] |
| ***Slc2a2*** | ZT0 | STD-VH vs. CAF-VH | 0.299 | 0.024 | 0.152 | 0.4[illegible] |
| | | CAF-VH vs. CAF-GSPE | -0.315 | 0.006 | -0.167 | 0.2[illegible] |
| | | STD-VH vs. CAF-GSPE | -0.016 | 0.884 | -0.015 | 0.9[illegible] |
| | ZT12 | STD-VH vs. CAF-VH | 0.067 | 0.234 | 0.016 | 0.8[illegible] |
| | | CAF-VH vs. CAF-GSPE | 0.019 | 0.759 | -0.208 | 0.0[illegible] |
| | | STD-VH vs. CAF-GSPE | 0.086 | 0.242 | -0.192 | 0.0[illegible] |

| | | | | | | |
|---|---|---|---|---|---|---|
| ***Ppargc1α*** | ZT0 | STD-VH vs. CAF-VH | -0.174 | 0.057 | 0.020 | 0.8[illegible] |
| | | CAF-VH vs. CAF-GSPE | -0.036 | 0.606 | -0.026 | 0.7[illegible] |
| | | STD-VH vs. CAF-GSPE | -0.210 | 0.012 | -0.006 | 0.9[illegible] |
| | ZT12 | STD-VH vs. CAF-VH | -0.208 | 0.076 | -0.276 | 0.0[illegible] |
| | | CAF-VH vs. CAF-GSPE | 0.001 | 0.991 | 0.051 | 0.6[illegible] |
| | | STD-VH vs. CAF-GSPE | -0.207 | 0.076 | -0.225 | 0.1[illegible] |
| ***Sirt1*** | ZT0 | STD-VH vs. CAF-VH | 0.172 | 0.003 | 0.044 | 0.5[illegible] |
| | | CAF-VH vs. CAF-GSPE | -0.030 | 0.513 | -0.007 | 0.9[illegible] |
| | | STD-VH vs. CAF-GSPE | 0.142 | 0.018 | 0.037 | 0.6[illegible] |
| | ZT12 | STD-VH vs. CAF-VH | 0.081 | 0.106 | -0.122 | 0.0[illegible] |
| | | CAF-VH vs. CAF-GSPE | -0.073 | 0.042 | 0.007 | 0.8[illegible] |
| | | STD-VH vs. CAF-GSPE | 0.008 | 0.878 | -0.115 | 0.1[illegible] |

Rats were fed a STD or CAF diet and were treated with vehicle or GSPE at the beginning of the light phase (ZT[illegible] (ZT12). d_MESOR represents the difference in MESOR values between the groups. d_amplitude represents the [illegible] the groups. d_phase represents the difference in acrophase values between the groups. The $p < 0.05$ indicates s[illegible] each rhythmic parameter.

**Supplementary Table 9.** Rhythmic parameters of genes related to hepatic glucose metabolism.

| Metabolite | ZT | Groups | *p* | MESOR | Amplitude | Acrophase [h] |
|---|---|---|---|---|---|---|
| **Pyruvic acid** | ZT0 | STD-VH | 0.003 | 0.692 | 0.172 | 13.488 |
| | | CAF-VH | 0.000 | 0.511 | 0.162 | 16.145 |
| | | CAF-GSPE | 0.041 | 0.469 | 0.071 | 13.902 |
| | ZT12 | STD-VH | 0.239 | 0.612 | 0.041 | 8.852 |
| | | CAF-VH | 0.062 | 0.458 | 0.060 | 8.479 |
| | | CAF-GSPE | 0.396 | 0.460 | 0.032 | 20.971 |
| **Lactic acid** | ZT0 | STD-VH | 0.004 | 4.372 | 0.341 | 10.959 |
| | | CAF-VH | 0.003 | 4.356 | 0.288 | 16.364 |
| | | CAF-GSPE | 0.202 | 4.324 | 0.092 | 13.599 |
| | ZT12 | STD-VH | 0.571 | 4.461 | 0.042 | 1.999 |
| | | CAF-VH | 0.035 | 4.349 | 0.192 | 8.512 |
| | | CAF-GSPE | 0.562 | 4.381 | 0.053 | 13.207 |
| **Alanine** | ZT0 | STD-VH | 0.015 | 3.057 | 0.263 | 9.393 |
| | | CAF-VH | 0.021 | 3.157 | 0.224 | 17.446 |
| | | CAF-GSPE | 0.178 | 3.164 | 0.098 | 17.709 |
| | ZT12 | STD-VH | 0.115 | 3.189 | 0.108 | 4.834 |
| | | CAF-VH | 0.033 | 3.195 | 0.123 | 8.499 |
| | | CAF-GSPE | 0.292 | 3.256 | 0.084 | 13.816 |
| **2-HydroxyButyric acid** | ZT0 | STD-VH | 0.082 | 0.565 | 0.133 | 16.497 |
| | | CAF-VH | 0.040 | 0.490 | 0.150 | 16.996 |
| | | CAF-GSPE | 0.255 | 0.451 | 0.081 | 18.054 |
| | ZT12 | STD-VH | 0.002 | 0.635 | 0.253 | 17.041 |
| | | CAF-VH | 0.282 | 0.586 | 0.082 | 16.885 |
| | | CAF-GSPE | 0.017 | 0.508 | 0.177 | 17.928 |
| **Sarcosine** | ZT0 | STD-VH | 0.211 | 0.040 | 0.007 | 21.291 |
| | | CAF-VH | 0.013 | 0.021 | 0.006 | 18.877 |
| | | CAF-GSPE | 0.017 | 0.025 | 0.006 | 19.362 |
| | ZT12 | STD-VH | 0.644 | 0.037 | 0.002 | 9.624 |
| | | CAF-VH | 0.010 | 0.023 | 0.008 | 14.656 |
| | | CAF-GSPE | 0.101 | 0.026 | 0.005 | 22.555 |
| **Valine** | ZT0 | STD-VH | 0.166 | 0.842 | 0.070 | 11.575 |

| | | | | | | |
|---|---|---|---|---|---|---|
| | | CAF-VH | 0.011 | 0.781 | 0.121 | 17.247 |
| | | CAF-GSPE | 0.012 | 0.798 | 0.109 | 17.878 |
| | ZT12 | STD-VH | 0.323 | 0.915 | 0.061 | 15.606 |
| | | CAF-VH | 0.694 | 0.791 | 0.015 | 14.399 |
| | | CAF-GSPE | 0.064 | 0.829 | 0.123 | 17.229 |
| **Urea** | ZT0 | STD-VH | 0.008 | 1.075 | 0.162 | 7.176 |
| | | CAF-VH | 0.602 | 0.739 | 0.026 | 11.482 |
| | | CAF-GSPE | 0.046 | 0.654 | 0.126 | 4.634 |
| | ZT12 | STD-VH | 0.054 | 1.037 | 0.162 | 6.406 |
| | | CAF-VH | 0.001 | 0.829 | 0.223 | 6.691 |
| | | CAF-GSPE | 0.398 | 0.774 | 0.057 | 2.770 |
| **Leucine** | ZT0 | STD-VH | 0.113 | 0.914 | 0.084 | 12.337 |
| | | CAF-VH | 0.007 | 0.870 | 0.148 | 17.400 |
| | | CAF-GSPE | 0.005 | 0.891 | 0.140 | 17.412 |
| | ZT12 | STD-VH | 0.221 | 1.014 | 0.084 | 16.882 |
| | | CAF-VH | 0.528 | 0.867 | 0.029 | 15.995 |
| | | CAF-GSPE | 0.040 | 0.918 | 0.148 | 17.461 |
| **Phosphoric acid** | ZT0 | STD-VH | 0.025 | 5.512 | 0.112 | 14.572 |
| | | CAF-VH | 0.001 | 5.476 | 0.200 | 18.197 |
| | | CAF-GSPE | 0.009 | 5.431 | 0.167 | 15.418 |
| | ZT12 | STD-VH | 0.068 | 5.534 | 0.099 | 15.606 |
| | | CAF-VH | 0.662 | 5.455 | 0.025 | 11.603 |
| | | CAF-GSPE | 0.037 | 5.479 | 0.089 | 13.006 |
| **Glycerol** | ZT0 | STD-VH | 0.048 | 0.177 | 0.065 | 15.971 |
| | | CAF-VH | 0.009 | 0.165 | 0.098 | 19.306 |
| | | CAF-GSPE | 0.033 | 0.172 | 0.074 | 16.456 |
| | ZT12 | STD-VH | 0.006 | 0.210 | 0.098 | 16.264 |
| | | CAF-VH | 0.174 | 0.185 | 0.047 | 16.791 |
| | | CAF-GSPE | 0.017 | 0.184 | 0.086 | 13.378 |
| **Isoleucine** | ZT0 | STD-VH | 0.249 | 0.531 | 0.042 | 12.182 |
| | | CAF-VH | 0.007 | 0.503 | 0.091 | 17.201 |
| | | CAF-GSPE | 0.006 | 0.513 | 0.082 | 17.787 |
| | ZT12 | STD-VH | 0.248 | 0.585 | 0.051 | 16.539 |
| | | CAF-VH | 0.479 | 0.500 | 0.020 | 16.393 |
| | | CAF-GSPE | 0.053 | 0.529 | 0.091 | 17.421 |
| **Proline** | ZT0 | STD-VH | 0.035 | 1.711 | 0.182 | 11.939 |

| | | | | | | |
|---|---|---|---|---|---|---|
| | | CAF-VH | 0.006 | 1.625 | 0.249 | 17.698 |
| | | CAF-GSPE | 0.040 | 1.690 | 0.171 | 17.300 |
| | ZT12 | STD-VH | 0.076 | 1.801 | 0.184 | 16.341 |
| | | CAF-VH | 0.175 | 1.627 | 0.107 | 16.346 |
| | | CAF-GSPE | 0.058 | 1.705 | 0.188 | 17.206 |
| **Glycine** | ZT0 | STD-VH | 0.046 | 1.824 | 0.142 | 9.331 |
| | | CAF-VH | 0.141 | 1.931 | 0.118 | 19.809 |
| | | CAF-GSPE | 0.017 | 1.989 | 0.155 | 16.641 |
| | ZT12 | STD-VH | 0.178 | 1.928 | 0.095 | 12.556 |
| | | CAF-VH | 0.212 | 1.843 | 0.115 | 19.551 |
| | | CAF-GSPE | 0.204 | 1.999 | 0.085 | 14.985 |
| **Succinic acid** | ZT0 | STD-VH | 0.316 | 0.938 | 0.095 | 6.651 |
| | | CAF-VH | 0.051 | 1.016 | 0.186 | 18.591 |
| | | CAF-GSPE | 0.386 | 1.023 | 0.082 | 3.130 |
| | ZT12 | STD-VH | 0.043 | 0.937 | 0.173 | 9.294 |
| | | CAF-VH | 0.280 | 1.068 | 0.085 | 20.087 |
| | | CAF-GSPE | 0.072 | 1.111 | 0.184 | 13.813 |
| **Fumaric acid** | ZT0 | STD-VH | 0.001 | 2.970 | 0.438 | 8.601 |
| | | CAF-VH | 0.368 | 2.844 | 0.130 | 7.186 |
| | | CAF-GSPE | 0.496 | 3.281 | 0.081 | 2.149 |
| | ZT12 | STD-VH | 0.001 | 3.345 | 0.603 | 2.750 |
| | | CAF-VH | 0.009 | 3.088 | 0.336 | 2.751 |
| | | CAF-GSPE | 0.078 | 3.261 | 0.257 | 23.355 |
| **Serine** | ZT0 | STD-VH | 0.046 | 0.775 | 0.168 | 8.481 |
| | | CAF-VH | 0.726 | 1.476 | 0.077 | 23.708 |
| | | CAF-GSPE | 0.149 | 1.650 | 0.228 | 16.390 |
| | ZT12 | STD-VH | 0.004 | 0.906 | 0.215 | 3.640 |
| | | CAF-VH | 0.477 | 1.191 | 0.110 | 6.121 |
| | | CAF-GSPE | 0.362 | 1.592 | 0.191 | 4.562 |
| **Threonine** | ZT0 | STD-VH | 0.051 | 0.485 | 0.066 | 10.395 |
| | | CAF-VH | 0.214 | 0.613 | 0.069 | 19.970 |
| | | CAF-GSPE | 0.005 | 0.658 | 0.117 | 16.188 |
| | ZT12 | STD-VH | 0.367 | 0.547 | 0.041 | 4.267 |
| | | CAF-VH | 0.495 | 0.568 | 0.033 | 8.151 |
| | | CAF-GSPE | 0.721 | 0.659 | 0.025 | 9.631 |
| **Nicotinamide** | ZT0 | STD-VH | 0.010 | 0.809 | 0.121 | 12.772 |

| | | | | | | |
|---|---|---|---|---|---|---|
| | | CAF-VH | 0.007 | 0.702 | 0.181 | 18.413 |
| | | CAF-GSPE | 0.029 | 0.736 | 0.160 | 18.195 |
| | ZT12 | STD-VH | 0.148 | 0.904 | 0.104 | 19.604 |
| | | CAF-VH | 0.268 | 0.742 | 0.054 | 19.344 |
| | | CAF-GSPE | 0.135 | 0.834 | 0.123 | 15.977 |
| **Malic acid** | ZT0 | STD-VH | 0.014 | 1.783 | 0.299 | 7.852 |
| | | CAF-VH | 0.215 | 1.923 | 0.174 | 7.185 |
| | | CAF-GSPE | 0.109 | 2.128 | 0.207 | 4.573 |
| | ZT12 | STD-VH | 0.000 | 1.987 | 0.574 | 2.228 |
| | | CAF-VH | 0.001 | 2.012 | 0.335 | 4.223 |
| | | CAF-GSPE | 0.025 | 2.025 | 0.262 | 0.190 |
| **Methionine** | ZT0 | STD-VH | 0.147 | 1.105 | 0.037 | 15.725 |
| | | CAF-VH | 0.035 | 1.076 | 0.039 | 18.866 |
| | | CAF-GSPE | 0.172 | 1.069 | 0.031 | 16.791 |
| | ZT12 | STD-VH | 0.047 | 1.131 | 0.048 | 17.113 |
| | | CAF-VH | 0.226 | 1.064 | 0.019 | 15.529 |
| | | CAF-GSPE | 0.145 | 1.073 | 0.031 | 17.480 |
| **Oxoproline** | ZT0 | STD-VH | 0.015 | 7.358 | 0.813 | 19.836 |
| | | CAF-VH | 0.143 | 6.812 | 0.487 | 17.191 |
| | | CAF-GSPE | 0.380 | 6.042 | 0.277 | 16.008 |
| | ZT12 | STD-VH | 0.011 | 6.895 | 0.912 | 18.343 |
| | | CAF-VH | 0.222 | 7.067 | 0.409 | 11.499 |
| | | CAF-GSPE | 0.324 | 6.688 | 0.432 | 17.107 |
| **Aspartic acid** | ZT0 | STD-VH | 0.136 | 0.916 | 0.096 | 7.102 |
| | | CAF-VH | 0.114 | 1.015 | 0.123 | 6.896 |
| | | CAF-GSPE | 0.577 | 1.017 | 0.038 | 1.730 |
| | ZT12 | STD-VH | 0.033 | 0.963 | 0.155 | 0.120 |
| | | CAF-VH | 0.703 | 0.914 | 0.023 | 4.064 |
| | | CAF-GSPE | 0.042 | 1.011 | 0.223 | 21.327 |
| **4-Hydroxyproline** | ZT0 | STD-VH | 0.013 | 0.226 | 0.079 | 8.401 |
| | | CAF-VH | 0.057 | 0.216 | 0.084 | 16.300 |
| | | CAF-GSPE | 0.062 | 0.193 | 0.048 | 0.754 |
| | ZT12 | STD-VH | 0.074 | 0.227 | 0.031 | 7.081 |
| | | CAF-VH | 0.880 | 0.226 | 0.005 | 2.136 |
| | | CAF-GSPE | 0.135 | 0.257 | 0.066 | 22.283 |
| | ZT0 | STD-VH | 0.055 | 0.044 | 0.012 | 5.489 |

| | | | | | | |
|---|---|---|---|---|---|---|
| **a-ketoglutaric acid** | | CAF-VH | 0.018 | 0.043 | 0.014 | 8.077 |
| | | CAF-GSPE | 0.027 | 0.062 | 0.016 | 1.463 |
| | ZT12 | STD-VH | 0.118 | 0.047 | 0.010 | 2.272 |
| | | CAF-VH | 0.232 | 0.041 | 0.007 | 0.366 |
| | | CAF-GSPE | 0.401 | 0.050 | 0.007 | 22.689 |
| **Glutamic acid** | ZT0 | STD-VH | 0.241 | 2.197 | 0.154 | 6.645 |
| | | CAF-VH | 0.382 | 2.470 | 0.092 | 17.568 |
| | | CAF-GSPE | 0.006 | 2.512 | 0.301 | 21.895 |
| | ZT12 | STD-VH | 0.366 | 2.268 | 0.105 | 3.484 |
| | | CAF-VH | 0.027 | 2.366 | 0.236 | 23.209 |
| | | CAF-GSPE | 0.682 | 2.545 | 0.071 | 16.056 |
| **Phenylalanine** | ZT0 | STD-VH | 0.204 | 0.449 | 0.041 | 12.041 |
| | | CAF-VH | 0.015 | 0.444 | 0.084 | 16.916 |
| | | CAF-GSPE | 0.007 | 0.441 | 0.068 | 17.157 |
| | ZT12 | STD-VH | 0.240 | 0.509 | 0.040 | 14.278 |
| | | CAF-VH | 0.714 | 0.437 | 0.008 | 19.355 |
| | | CAF-GSPE | 0.072 | 0.452 | 0.075 | 16.997 |
| **Taurine** | ZT0 | STD-VH | 0.034 | 2.437 | 0.854 | 18.933 |
| | | CAF-VH | 0.043 | 2.037 | 0.769 | 16.408 |
| | | CAF-GSPE | 0.544 | 1.302 | 0.211 | 17.536 |
| | ZT12 | STD-VH | 0.010 | 1.672 | 0.956 | 17.236 |
| | | CAF-VH | 0.087 | 1.853 | 0.444 | 11.308 |
| | | CAF-GSPE | 0.212 | 1.540 | 0.479 | 16.909 |
| **d-Ribose** | ZT0 | STD-VH | 0.550 | 0.020 | 0.001 | 12.523 |
| | | CAF-VH | 0.427 | 0.044 | 0.008 | 17.342 |
| | | CAF-GSPE | 0.198 | 0.041 | 0.021 | 10.664 |
| | ZT12 | STD-VH | 0.323 | 0.033 | 0.006 | 17.049 |
| | | CAF-VH | 0.145 | 0.023 | 0.010 | 21.650 |
| | | CAF-GSPE | 0.046 | 0.030 | 0.011 | 15.708 |
| **d-Xylitol** | ZT0 | STD-VH | 0.053 | 0.019 | 0.006 | 9.296 |
| | | CAF-VH | 0.001 | 0.016 | 0.008 | 18.328 |
| | | CAF-GSPE | 0.165 | 0.016 | 0.003 | 14.707 |
| | ZT12 | STD-VH | 0.268 | 0.021 | 0.003 | 14.323 |
| | | CAF-VH | 0.778 | 0.017 | 0.000 | 14.766 |
| | | CAF-GSPE | 0.010 | 0.019 | 0.008 | 13.167 |
| | ZT0 | STD-VH | 0.005 | 1.199 | 0.272 | 9.235 |

| | | | | | | |
|---|---|---|---|---|---|---|
| **Glycerol-1-phosphate** | | CAF-VH | 0.516 | 1.277 | 0.085 | 14.949 |
| | | CAF-GSPE | 0.176 | 1.264 | 0.140 | 2.473 |
| | ZT12 | STD-VH | 0.141 | 1.312 | 0.153 | 3.823 |
| | | CAF-VH | 0.354 | 1.308 | 0.086 | 3.906 |
| | | CAF-GSPE | 0.084 | 1.426 | 0.216 | 18.479 |
| **Hypoxanthine** | ZT0 | STD-VH | 0.013 | 0.132 | 0.042 | 15.275 |
| | | CAF-VH | 0.007 | 0.122 | 0.058 | 18.918 |
| | | CAF-GSPE | 0.004 | 0.139 | 0.065 | 16.641 |
| | ZT12 | STD-VH | 0.129 | 0.201 | 0.068 | 21.512 |
| | | CAF-VH | 0.696 | 0.126 | 0.010 | 9.644 |
| | | CAF-GSPE | 0.459 | 0.161 | 0.026 | 15.530 |
| **Ornithine** | ZT0 | STD-VH | 0.015 | 0.110 | 0.049 | 8.181 |
| | | CAF-VH | 0.408 | 0.102 | 0.010 | 0.914 |
| | | CAF-GSPE | 0.244 | 0.109 | 0.014 | 14.520 |
| | ZT12 | STD-VH | 0.076 | 0.147 | 0.058 | 3.645 |
| | | CAF-VH | 0.290 | 0.105 | 0.010 | 9.524 |
| | | CAF-GSPE | 0.720 | 0.128 | 0.007 | 4.592 |
| **Citric acid** | ZT0 | STD-VH | 0.014 | 0.355 | 0.184 | 6.240 |
| | | CAF-VH | 0.002 | 0.369 | 0.168 | 6.603 |
| | | CAF-GSPE | 0.002 | 0.460 | 0.203 | 5.450 |
| | ZT12 | STD-VH | 0.040 | 0.333 | 0.111 | 4.183 |
| | | CAF-VH | 0.278 | 0.337 | 0.061 | 4.238 |
| | | CAF-GSPE | 0.081 | 0.369 | 0.132 | 22.324 |
| **Adenine** | ZT0 | STD-VH | 0.798 | 0.034 | 0.001 | 4.051 |
| | | CAF-VH | 0.091 | 0.033 | 0.007 | 16.566 |
| | | CAF-GSPE | 0.322 | 0.030 | 0.003 | 8.821 |
| | ZT12 | STD-VH | 0.818 | 0.037 | 0.001 | 23.940 |
| | | CAF-VH | 0.034 | 0.030 | 0.007 | 0.893 |
| | | CAF-GSPE | 0.028 | 0.033 | 0.010 | 14.455 |
| **Hydroxyphenyllactic acid** | ZT0 | STD-VH | 0.176 | 0.457 | 0.098 | 18.165 |
| | | CAF-VH | 0.113 | 0.419 | 0.098 | 18.516 |
| | | CAF-GSPE | 0.021 | 0.384 | 0.138 | 19.955 |
| | ZT12 | STD-VH | 0.005 | 0.481 | 0.268 | 16.828 |
| | | CAF-VH | 0.800 | 0.474 | 0.015 | 22.610 |
| | | CAF-GSPE | 0.062 | 0.469 | 0.184 | 18.586 |
| **d-Glucose** | ZT0 | STD-VH | 0.222 | 7.260 | 0.023 | 10.526 |

| | | | | | | |
|---|---|---|---|---|---|---|
| | | CAF-VH | 0.032 | 7.269 | 0.049 | 18.564 |
| | | CAF-GSPE | 0.196 | 7.273 | 0.025 | 18.076 |
| | ZT12 | STD-VH | 0.040 | 7.219 | 0.045 | 13.824 |
| | | CAF-VH | 0.362 | 7.226 | 0.019 | 13.526 |
| | | CAF-GSPE | 0.056 | 7.252 | 0.055 | 15.437 |
| **d-Mannonic acid** | ZT0 | STD-VH | 0.010 | 0.037 | 0.008 | 15.821 |
| | | CAF-VH | 0.001 | 0.032 | 0.012 | 17.969 |
| | | CAF-GSPE | 0.023 | 0.033 | 0.010 | 16.605 |
| | ZT12 | STD-VH | 0.340 | 0.035 | 0.006 | 9.724 |
| | | CAF-VH | 0.002 | 0.033 | 0.010 | 13.877 |
| | | CAF-GSPE | 0.082 | 0.040 | 0.008 | 15.607 |
| **d-Sorbitol** | ZT0 | STD-VH | 0.410 | 0.130 | 0.027 | 15.293 |
| | | CAF-VH | 0.015 | 0.115 | 0.081 | 19.357 |
| | | CAF-GSPE | 0.095 | 0.117 | 0.057 | 14.921 |
| | ZT12 | STD-VH | 0.011 | 0.156 | 0.093 | 16.249 |
| | | CAF-VH | 0.075 | 0.118 | 0.052 | 17.136 |
| | | CAF-GSPE | 0.166 | 0.280 | 0.303 | 8.028 |
| **d-Galactitol** | ZT0 | STD-VH | 0.003 | 0.025 | 0.009 | 10.118 |
| | | CAF-VH | 0.010 | 0.029 | 0.010 | 16.555 |
| | | CAF-GSPE | 0.095 | 0.029 | 0.006 | 13.115 |
| | ZT12 | STD-VH | 0.114 | 0.020 | 0.002 | 11.369 |
| | | CAF-VH | 0.362 | 0.021 | 0.001 | 11.226 |
| | | CAF-GSPE | 0.018 | 0.023 | 0.006 | 14.471 |
| **d-Gluconic acid** | ZT0 | STD-VH | 0.075 | 0.090 | 0.039 | 16.473 |
| | | CAF-VH | 0.005 | 0.069 | 0.042 | 18.533 |
| | | CAF-GSPE | 0.032 | 0.083 | 0.045 | 16.154 |
| | ZT12 | STD-VH | 0.014 | 0.110 | 0.059 | 16.429 |
| | | CAF-VH | 0.030 | 0.079 | 0.041 | 15.859 |
| | | CAF-GSPE | 0.065 | 0.086 | 0.032 | 14.240 |
| **Palmitoleic acid** | ZT0 | STD-VH | 0.186 | 0.079 | 0.009 | 7.219 |
| | | CAF-VH | 0.309 | 0.067 | 0.010 | 14.997 |
| | | CAF-GSPE | 0.065 | 0.060 | 0.012 | 10.043 |
| | ZT12 | STD-VH | 0.033 | 0.078 | 0.018 | 7.711 |
| | | CAF-VH | 0.020 | 0.059 | 0.011 | 10.039 |
| | | CAF-GSPE | 0.377 | 0.068 | 0.013 | 19.253 |
| **Xanthine** | ZT0 | STD-VH | 0.086 | 0.093 | 0.043 | 14.284 |

| | | | | | | |
|---|---|---|---|---|---|---|
| | | CAF-VH | 0.011 | 0.112 | 0.074 | 18.593 |
| | | CAF-GSPE | 0.020 | 0.130 | 0.086 | 15.513 |
| | ZT12 | STD-VH | 0.137 | 0.158 | 0.068 | 16.350 |
| | | CAF-VH | 0.176 | 0.108 | 0.050 | 16.252 |
| | | CAF-GSPE | 0.056 | 0.140 | 0.076 | 13.330 |
| **d-Glucuronic acid** | ZT0 | STD-VH | 0.047 | 0.004 | 0.001 | 14.451 |
| | | CAF-VH | 0.004 | 0.004 | 0.001 | 17.937 |
| | | CAF-GSPE | 0.009 | 0.004 | 0.001 | 17.821 |
| | ZT12 | STD-VH | 0.031 | 0.005 | 0.002 | 16.001 |
| | | CAF-VH | 0.018 | 0.003 | 0.001 | 16.936 |
| | | CAF-GSPE | 0.033 | 0.004 | 0.001 | 13.936 |
| **myo-Inositol** | ZT0 | STD-VH | 0.033 | 0.390 | 0.067 | 11.979 |
| | | CAF-VH | 0.024 | 0.290 | 0.033 | 17.778 |
| | | CAF-GSPE | 0.112 | 0.293 | 0.034 | 18.277 |
| | ZT12 | STD-VH | 0.740 | 0.424 | 0.012 | 12.917 |
| | | CAF-VH | 0.745 | 0.295 | 0.006 | 13.357 |
| | | CAF-GSPE | 0.104 | 0.315 | 0.033 | 19.004 |
| **Ribose 5-phosphate** | ZT0 | STD-VH | 0.137 | 0.030 | 0.006 | 4.863 |
| | | CAF-VH | 0.674 | 0.051 | 0.005 | 15.347 |
| | | CAF-GSPE | 0.189 | 0.048 | 0.022 | 9.668 |
| | ZT12 | STD-VH | 0.434 | 0.043 | 0.005 | 6.218 |
| | | CAF-VH | 0.197 | 0.030 | 0.010 | 23.659 |
| | | CAF-GSPE | 0.255 | 0.036 | 0.005 | 16.715 |
| **Sedoheptulose** | ZT0 | STD-VH | 0.023 | 0.022 | 0.006 | 7.635 |
| | | CAF-VH | 0.011 | 0.019 | 0.005 | 16.618 |
| | | CAF-GSPE | 0.668 | 0.017 | 0.001 | 5.918 |
| | ZT12 | STD-VH | 0.128 | 0.025 | 0.005 | 4.922 |
| | | CAF-VH | 0.056 | 0.020 | 0.005 | 3.316 |
| | | CAF-GSPE | 0.293 | 0.022 | 0.003 | 16.727 |
| **Oleic acid-iso1** | ZT0 | STD-VH | 0.270 | 0.180 | 0.027 | 10.525 |
| | | CAF-VH | 0.161 | 0.465 | 0.104 | 12.454 |
| | | CAF-GSPE | 0.633 | 0.448 | 0.025 | 11.551 |
| | ZT12 | STD-VH | 0.136 | 0.247 | 0.035 | 0.754 |
| | | CAF-VH | 0.665 | 0.456 | 0.030 | 10.894 |
| | | CAF-GSPE | 0.390 | 0.420 | 0.060 | 18.820 |
| | ZT0 | STD-VH | 0.078 | 0.180 | 0.051 | 13.536 |

| | | | | | | |
|---|---|---|---|---|---|---|
| **Oleic acid-iso2** | | CAF-VH | 0.075 | 0.151 | 0.058 | 12.917 |
| | | CAF-GSPE | 0.563 | 0.117 | 0.010 | 17.437 |
| | ZT12 | STD-VH | 0.554 | 0.188 | 0.019 | 9.257 |
| | | CAF-VH | 0.967 | 0.126 | 0.001 | 22.417 |
| | | CAF-GSPE | 0.165 | 0.127 | 0.038 | 18.239 |
| **Stearic acid** | ZT0 | STD-VH | 0.161 | 3.904 | 0.138 | 9.770 |
| | | CAF-VH | 0.033 | 3.957 | 0.121 | 13.517 |
| | | CAF-GSPE | 0.124 | 4.082 | 0.204 | 8.096 |
| | ZT12 | STD-VH | 0.715 | 3.982 | 0.023 | 3.536 |
| | | CAF-VH | 0.634 | 3.959 | 0.035 | 16.576 |
| | | CAF-GSPE | 0.450 | 4.074 | 0.073 | 13.470 |
| **Fructose 6-phosphate** | ZT0 | STD-VH | 0.023 | 0.030 | 0.022 | 5.558 |
| | | CAF-VH | 0.003 | 0.030 | 0.017 | 8.433 |
| | | CAF-GSPE | 0.000 | 0.020 | 0.010 | 6.427 |
| | ZT12 | STD-VH | 0.004 | 0.026 | 0.018 | 5.037 |
| | | CAF-VH | 0.033 | 0.022 | 0.012 | 5.319 |
| | | CAF-GSPE | 0.113 | 0.024 | 0.006 | 4.223 |
| **Glucose 6-phosphate** | ZT0 | STD-VH | 0.022 | 0.067 | 0.057 | 5.497 |
| | | CAF-VH | 0.003 | 0.069 | 0.047 | 8.624 |
| | | CAF-GSPE | 0.000 | 0.045 | 0.024 | 5.622 |
| | ZT12 | STD-VH | 0.003 | 0.055 | 0.040 | 5.008 |
| | | CAF-VH | 0.039 | 0.047 | 0.027 | 5.634 |
| | | CAF-GSPE | 0.112 | 0.052 | 0.015 | 4.275 |
| **Arachidic acid** | ZT0 | STD-VH | 0.109 | 0.078 | 0.014 | 7.789 |
| | | CAF-VH | 0.347 | 0.074 | 0.003 | 14.030 |
| | | CAF-GSPE | 0.095 | 0.089 | 0.022 | 7.955 |
| | ZT12 | STD-VH | 0.352 | 0.081 | 0.005 | 16.309 |
| | | CAF-VH | 0.307 | 0.077 | 0.005 | 20.994 |
| | | CAF-GSPE | 0.398 | 0.087 | 0.006 | 11.474 |
| **Inosine** | ZT0 | STD-VH | 0.018 | 0.498 | 0.135 | 15.896 |
| | | CAF-VH | 0.012 | 0.492 | 0.218 | 17.017 |
| | | CAF-GSPE | 0.050 | 0.474 | 0.154 | 16.444 |
| | ZT12 | STD-VH | 0.125 | 0.666 | 0.183 | 21.206 |
| | | CAF-VH | 0.597 | 0.473 | 0.045 | 5.099 |
| | | CAF-GSPE | 0.271 | 0.565 | 0.135 | 16.291 |
| **Adenosine** | ZT0 | STD-VH | 0.070 | 0.134 | 0.038 | 7.149 |

| | | | | | | |
|---|---|---|---|---|---|---|
| | | CAF-VH | 0.814 | 0.132 | 0.004 | 15.166 |
| | | CAF-GSPE | 0.120 | 0.160 | 0.032 | 22.586 |
| | ZT12 | STD-VH | 0.236 | 0.177 | 0.059 | 11.201 |
| | | CAF-VH | 0.536 | 0.164 | 0.016 | 20.849 |
| | | CAF-GSPE | 0.152 | 0.186 | 0.044 | 14.415 |
| **Xanthosine** | ZT0 | STD-VH | 0.034 | 0.075 | 0.030 | 15.094 |
| | | CAF-VH | 0.004 | 0.077 | 0.041 | 17.330 |
| | | CAF-GSPE | 0.017 | 0.073 | 0.038 | 15.270 |
| | ZT12 | STD-VH | 0.008 | 0.092 | 0.046 | 15.359 |
| | | CAF-VH | 0.089 | 0.083 | 0.026 | 15.524 |
| | | CAF-GSPE | 0.010 | 0.091 | 0.051 | 13.556 |
| **d-Maltose** | ZT0 | STD-VH | 0.849 | 0.201 | 0.011 | 15.463 |
| | | CAF-VH | 0.123 | 0.200 | 0.113 | 18.511 |
| | | CAF-GSPE | 0.483 | 0.221 | 0.048 | 3.121 |
| | ZT12 | STD-VH | 0.097 | 0.266 | 0.165 | 23.838 |
| | | CAF-VH | 0.828 | 0.364 | 0.024 | 1.560 |
| | | CAF-GSPE | 0.495 | 0.313 | 0.064 | 9.917 |
| **Lignoceric acid** | ZT0 | STD-VH | 0.772 | 0.305 | 0.022 | 14.670 |
| | | CAF-VH | 0.119 | 0.302 | 0.154 | 18.412 |
| | | CAF-GSPE | 0.625 | 0.323 | 0.043 | 3.808 |
| | ZT12 | STD-VH | 0.080 | 0.374 | 0.206 | 23.691 |
| | | CAF-VH | 0.853 | 0.534 | 0.030 | 0.598 |
| | | CAF-GSPE | 0.413 | 0.461 | 0.106 | 9.798 |
| **uridine 5-monophosphate** | ZT0 | STD-VH | 0.113 | 0.246 | 0.072 | 4.716 |
| | | CAF-VH | 0.182 | 0.244 | 0.035 | 7.838 |
| | | CAF-GSPE | 0.057 | 0.201 | 0.051 | 3.142 |
| | ZT12 | STD-VH | 0.014 | 0.214 | 0.098 | 6.217 |
| | | CAF-VH | 0.127 | 0.229 | 0.052 | 5.064 |
| | | CAF-GSPE | 0.111 | 0.220 | 0.049 | 1.068 |
| **inosine 5-monophosphate** | ZT0 | STD-VH | 0.160 | 0.052 | 0.014 | 8.043 |
| | | CAF-VH | 0.328 | 0.070 | 0.009 | 11.585 |
| | | CAF-GSPE | 0.076 | 0.056 | 0.014 | 11.078 |
| | ZT12 | STD-VH | 0.001 | 0.053 | 0.033 | 8.097 |
| | | CAF-VH | 0.056 | 0.049 | 0.015 | 5.169 |
| | | CAF-GSPE | 0.390 | 0.057 | 0.009 | 6.153 |
| | ZT0 | STD-VH | 0.035 | 0.630 | 0.171 | 5.845 |

| | | | | | | |
|---|---|---|---|---|---|---|
| **adenosine-5-monophosphate_diphosphate_triphosphate** | | CAF-VH | 0.055 | 0.651 | 0.081 | 7.287 |
| | | CAF-GSPE | 0.005 | 0.597 | 0.121 | 4.289 |
| | ZT12 | STD-VH | 0.008 | 0.595 | 0.187 | 6.913 |
| | | CAF-VH | 0.244 | 0.590 | 0.087 | 6.289 |
| | | CAF-GSPE | 0.108 | 0.601 | 0.102 | 23.934 |
| **Cholesterol** | ZT0 | STD-VH | 0.143 | 3.478 | 0.067 | 11.149 |
| | | CAF-VH | 0.067 | 3.295 | 0.136 | 18.585 |
| | | CAF-GSPE | 0.283 | 3.369 | 0.076 | 12.257 |
| | ZT12 | STD-VH | 0.103 | 3.546 | 0.077 | 10.035 |
| | | CAF-VH | 0.607 | 3.283 | 0.028 | 22.088 |
| | | CAF-GSPE | 0.180 | 3.388 | 0.069 | 9.826 |

**Supplementary Table 10.** Comparison of rhythmic parameters of liver metabolites between groups.

| Metabolite | ZT | Comparison | d_MESOR | p(d_MESOR) | d_amplitude | p |
|---|---|---|---|---|---|---|
| **Pyruvic acid** | ZT0 | STD-VH vs. CAF-VH | -0.181 | 0.000 | -0.010 | |
| | | CAF-VH vs. CAF-GSPE | -0.043 | 0.185 | -0.090 | |
| | | STD-VH vs. CAF-GSPE | -0.223 | 0.000 | -0.100 | |
| | ZT12 | STD-VH vs. CAF-VH | -0.154 | 0.000 | 0.019 | |
| | | CAF-VH vs. CAF-GSPE | 0.002 | 0.957 | -0.029 | |
| | | STD-VH vs. CAF-GSPE | -0.152 | 0.000 | -0.010 | |
| **Lactic acid** | ZT0 | STD-VH vs. CAF-VH | -0.016 | 0.859 | -0.052 | |
| | | CAF-VH vs. CAF-GSPE | -0.032 | 0.673 | -0.196 | |
| | | STD-VH vs. CAF-GSPE | -0.048 | 0.576 | -0.249 | |
| | ZT12 | STD-VH vs. CAF-VH | -0.111 | 0.159 | 0.150 | |
| | | CAF-VH vs. CAF-GSPE | 0.031 | 0.714 | -0.139 | |
| | | STD-VH vs. CAF-GSPE | -0.080 | 0.330 | 0.011 | |
| **Alanine** | ZT0 | STD-VH vs. CAF-VH | 0.100 | 0.275 | -0.040 | |
| | | CAF-VH vs. CAF-GSPE | 0.007 | 0.927 | -0.126 | |
| | | STD-VH vs. CAF-GSPE | 0.107 | 0.213 | -0.166 | |
| | ZT12 | STD-VH vs. CAF-VH | 0.006 | 0.918 | 0.014 | |
| | | CAF-VH vs. CAF-GSPE | 0.062 | 0.352 | -0.039 | |
| | | STD-VH vs. CAF-GSPE | 0.068 | 0.346 | -0.025 | |

| | | | | | |
|---|---|---|---|---|---|
| **2-HydroxyButyric acid** | ZT0 | STD-VH vs. CAF-VH | -0.075 | 0.283 | 0.016 |
| | | CAF-VH vs. CAF-GSPE | -0.039 | 0.569 | -0.069 |
| | | STD-VH vs. CAF-GSPE | -0.114 | 0.119 | -0.052 |
| | ZT12 | STD-VH vs. CAF-VH | -0.049 | 0.491 | -0.171 |
| | | CAF-VH vs. CAF-GSPE | -0.078 | 0.267 | 0.094 |
| | | STD-VH vs. CAF-GSPE | -0.127 | 0.063 | -0.077 |
| **Sarcosine** | ZT0 | STD-VH vs. CAF-VH | -0.020 | 0.000 | -0.001 |
| | | CAF-VH vs. CAF-GSPE | 0.005 | 0.028 | 0.000 |
| | | STD-VH vs. CAF-GSPE | -0.015 | 0.001 | -0.001 |
| | ZT12 | STD-VH vs. CAF-VH | -0.014 | 0.002 | 0.005 |
| | | CAF-VH vs. CAF-GSPE | 0.003 | 0.233 | -0.003 |
| | | STD-VH vs. CAF-GSPE | -0.011 | 0.015 | 0.003 |
| **Valine** | ZT0 | STD-VH vs. CAF-VH | -0.061 | 0.181 | 0.050 |
| | | CAF-VH vs. CAF-GSPE | 0.017 | 0.674 | -0.012 |
| | | STD-VH vs. CAF-GSPE | -0.044 | 0.321 | 0.039 |
| | ZT12 | STD-VH vs. CAF-VH | -0.123 | 0.019 | -0.046 |
| | | CAF-VH vs. CAF-GSPE | 0.038 | 0.458 | 0.108 |
| | | STD-VH vs. CAF-GSPE | -0.085 | 0.165 | 0.062 |
| **Urea** | ZT0 | STD-VH vs. CAF-VH | -0.336 | 0.000 | -0.135 |
| | | CAF-VH vs. CAF-GSPE | -0.084 | 0.126 | 0.099 |

| | | | | | |
|---|---|---|---|---|---|
| | | STD-VH vs. CAF-GSPE | -0.420 | 0.000 | -0.036 |
| | ZT12 | STD-VH vs. CAF-VH | -0.208 | 0.004 | 0.061 |
| | | CAF-VH vs. CAF-GSPE | -0.055 | 0.367 | -0.166 |
| | | STD-VH vs. CAF-GSPE | -0.264 | 0.001 | -0.105 |
| **Leucine** | ZT0 | STD-VH vs. CAF-VH | -0.044 | 0.362 | 0.064 |
| | | CAF-VH vs. CAF-GSPE | 0.021 | 0.642 | -0.007 |
| | | STD-VH vs. CAF-GSPE | -0.023 | 0.613 | 0.057 |
| | ZT12 | STD-VH vs. CAF-VH | -0.147 | 0.015 | -0.055 |
| | | CAF-VH vs. CAF-GSPE | 0.051 | 0.372 | 0.119 |
| | | STD-VH vs. CAF-GSPE | -0.096 | 0.155 | 0.064 |
| **Phosphoric acid** | ZT0 | STD-VH vs. CAF-VH | -0.036 | 0.436 | 0.088 |
| | | CAF-VH vs. CAF-GSPE | -0.045 | 0.380 | -0.033 |
| | | STD-VH vs. CAF-GSPE | -0.081 | 0.109 | 0.055 |
| | ZT12 | STD-VH vs. CAF-VH | -0.079 | 0.143 | -0.075 |
| | | CAF-VH vs. CAF-GSPE | 0.024 | 0.622 | 0.065 |
| | | STD-VH vs. CAF-GSPE | -0.055 | 0.224 | -0.010 |
| **Glycerol** | ZT0 | STD-VH vs. CAF-VH | -0.012 | 0.689 | 0.033 |
| | | CAF-VH vs. CAF-GSPE | 0.008 | 0.809 | -0.023 |
| | | STD-VH vs. CAF-GSPE | -0.005 | 0.877 | 0.009 |
| | ZT12 | STD-VH vs. CAF-VH | -0.025 | 0.430 | -0.051 |

| | | | | | |
|---|---|---|---|---|---|
| | | CAF-VH vs. CAF-GSPE | -0.001 | 0.974 | 0.039 |
| | | STD-VH vs. CAF-GSPE | -0.026 | 0.404 | -0.012 |
| **Isoleucine** | ZT0 | STD-VH vs. CAF-VH | -0.028 | 0.388 | 0.048 |
| | | CAF-VH vs. CAF-GSPE | 0.010 | 0.713 | -0.009 |
| | | STD-VH vs. CAF-GSPE | -0.018 | 0.564 | 0.039 |
| | ZT12 | STD-VH vs. CAF-VH | -0.085 | 0.026 | -0.031 |
| | | CAF-VH vs. CAF-GSPE | 0.029 | 0.437 | 0.071 |
| | | STD-VH vs. CAF-GSPE | -0.056 | 0.200 | 0.040 |
| **Proline** | ZT0 | STD-VH vs. CAF-VH | -0.085 | 0.273 | 0.068 |
| | | CAF-VH vs. CAF-GSPE | 0.065 | 0.399 | -0.079 |
| | | STD-VH vs. CAF-GSPE | -0.020 | 0.793 | -0.011 |
| | ZT12 | STD-VH vs. CAF-VH | -0.174 | 0.053 | -0.078 |
| | | CAF-VH vs. CAF-GSPE | 0.079 | 0.350 | 0.082 |
| | | STD-VH vs. CAF-GSPE | -0.095 | 0.317 | 0.004 |
| **Glycine** | ZT0 | STD-VH vs. CAF-VH | 0.107 | 0.140 | -0.023 |
| | | CAF-VH vs. CAF-GSPE | 0.058 | 0.396 | 0.037 |
| | | STD-VH vs. CAF-GSPE | 0.165 | 0.012 | 0.014 |
| | ZT12 | STD-VH vs. CAF-VH | -0.085 | 0.285 | 0.020 |
| | | CAF-VH vs. CAF-GSPE | 0.156 | 0.051 | -0.030 |
| | | STD-VH vs. CAF-GSPE | 0.071 | 0.285 | -0.010 |

| | | | | | |
|---|---|---|---|---|---|
| **Succinic acid** | ZT0 | STD-VH vs. CAF-VH | 0.078 | 0.387 | 0.091 |
| | | CAF-VH vs. CAF-GSPE | 0.006 | 0.942 | -0.104 |
| | | STD-VH vs. CAF-GSPE | 0.085 | 0.359 | -0.013 |
| | ZT12 | STD-VH vs. CAF-VH | 0.131 | 0.096 | -0.088 |
| | | CAF-VH vs. CAF-GSPE | 0.043 | 0.620 | 0.099 |
| | | STD-VH vs. CAF-GSPE | 0.174 | 0.053 | 0.012 |
| **Fumaric acid** | ZT0 | STD-VH vs. CAF-VH | -0.127 | 0.322 | -0.307 |
| | | CAF-VH vs. CAF-GSPE | 0.438 | 0.002 | -0.049 |
| | | STD-VH vs. CAF-GSPE | 0.311 | 0.009 | -0.357 |
| | ZT12 | STD-VH vs. CAF-VH | -0.257 | 0.060 | -0.267 |
| | | CAF-VH vs. CAF-GSPE | 0.173 | 0.171 | -0.078 |
| | | STD-VH vs. CAF-GSPE | -0.084 | 0.558 | -0.345 |
| **Serine** | ZT0 | STD-VH vs. CAF-VH | 0.701 | 0.000 | -0.090 |
| | | CAF-VH vs. CAF-GSPE | 0.174 | 0.365 | 0.151 |
| | | STD-VH vs. CAF-GSPE | 0.875 | 0.000 | 0.060 |
| | ZT12 | STD-VH vs. CAF-VH | 0.285 | 0.019 | -0.105 |
| | | CAF-VH vs. CAF-GSPE | 0.401 | 0.033 | 0.082 |
| | | STD-VH vs. CAF-GSPE | 0.685 | 0.000 | -0.024 |
| **Threonine** | ZT0 | STD-VH vs. CAF-VH | 0.129 | 0.006 | 0.003 |
| | | CAF-VH vs. CAF-GSPE | 0.045 | 0.333 | 0.048 |

| | | | | | |
|---|---|---|---|---|---|
| | | STD-VH vs. CAF-GSPE | 0.173 | 0.000 | 0.051 |
| | ZT12 | STD-VH vs. CAF-VH | 0.021 | 0.641 | -0.008 |
| | | CAF-VH vs. CAF-GSPE | 0.091 | 0.131 | -0.009 |
| | | STD-VH vs. CAF-GSPE | 0.112 | 0.059 | -0.016 |
| **Nicotinamide** | ZT0 | STD-VH vs. CAF-VH | -0.107 | 0.039 | 0.060 |
| | | CAF-VH vs. CAF-GSPE | 0.034 | 0.587 | -0.021 |
| | | STD-VH vs. CAF-GSPE | -0.073 | 0.190 | 0.039 |
| | ZT12 | STD-VH vs. CAF-VH | -0.162 | 0.010 | -0.050 |
| | | CAF-VH vs. CAF-GSPE | 0.092 | 0.163 | 0.069 |
| | | STD-VH vs. CAF-GSPE | -0.070 | 0.344 | 0.019 |
| **Malic acid** | ZT0 | STD-VH vs. CAF-VH | 0.140 | 0.255 | -0.125 |
| | | CAF-VH vs. CAF-GSPE | 0.205 | 0.124 | 0.033 |
| | | STD-VH vs. CAF-GSPE | 0.345 | 0.005 | -0.092 |
| | ZT12 | STD-VH vs. CAF-VH | 0.025 | 0.807 | -0.238 |
| | | CAF-VH vs. CAF-GSPE | 0.013 | 0.890 | -0.073 |
| | | STD-VH vs. CAF-GSPE | 0.038 | 0.734 | -0.311 |
| **Methionine** | ZT0 | STD-VH vs. CAF-VH | -0.029 | 0.177 | 0.002 |
| | | CAF-VH vs. CAF-GSPE | -0.007 | 0.701 | -0.008 |
| | | STD-VH vs. CAF-GSPE | -0.036 | 0.128 | -0.006 |
| | ZT12 | STD-VH vs. CAF-VH | -0.066 | 0.001 | -0.029 |

| | | | | | |
|---|---|---|---|---|---|
| | | CAF-VH vs. CAF-GSPE | 0.009 | 0.635 | 0.012 |
| | | STD-VH vs. CAF-GSPE | -0.058 | 0.010 | -0.016 |
| **Oxoproline** | ZT0 | STD-VH vs. CAF-VH | -0.546 | 0.082 | -0.326 |
| | | CAF-VH vs. CAF-GSPE | -0.770 | 0.020 | -0.211 |
| | | STD-VH vs. CAF-GSPE | -1.316 | 0.000 | -0.537 |
| | ZT12 | STD-VH vs. CAF-VH | 0.172 | 0.587 | -0.503 |
| | | CAF-VH vs. CAF-GSPE | -0.379 | 0.319 | 0.023 |
| | | STD-VH vs. CAF-GSPE | -0.207 | 0.580 | -0.480 |
| **Aspartic acid** | ZT0 | STD-VH vs. CAF-VH | 0.098 | 0.153 | 0.027 |
| | | CAF-VH vs. CAF-GSPE | 0.002 | 0.971 | -0.084 |
| | | STD-VH vs. CAF-GSPE | 0.101 | 0.119 | -0.058 |
| | ZT12 | STD-VH vs. CAF-VH | -0.050 | 0.436 | -0.132 |
| | | CAF-VH vs. CAF-GSPE | 0.097 | 0.243 | 0.199 |
| | | STD-VH vs. CAF-GSPE | 0.048 | 0.572 | 0.067 |
| **4-Hydroxyproline** | ZT0 | STD-VH vs. CAF-VH | -0.010 | 0.763 | 0.004 |
| | | CAF-VH vs. CAF-GSPE | -0.023 | 0.494 | -0.036 |
| | | STD-VH vs. CAF-GSPE | -0.033 | 0.197 | -0.032 |
| | ZT12 | STD-VH vs. CAF-VH | -0.001 | 0.965 | -0.026 |
| | | CAF-VH vs. CAF-GSPE | 0.031 | 0.425 | 0.061 |
| | | STD-VH vs. CAF-GSPE | 0.029 | 0.356 | 0.035 |

| | | | | | |
|---|---|---|---|---|---|
| **a-ketoglutaric acid** | ZT0 | STD-VH vs. CAF-VH | 0.000 | 0.950 | 0.002 |
| | | CAF-VH vs. CAF-GSPE | 0.019 | 0.002 | 0.002 |
| | | STD-VH vs. CAF-GSPE | 0.018 | 0.004 | 0.004 |
| | ZT12 | STD-VH vs. CAF-VH | -0.006 | 0.333 | -0.003 |
| | | CAF-VH vs. CAF-GSPE | 0.009 | 0.176 | -0.001 |
| | | STD-VH vs. CAF-GSPE | 0.003 | 0.618 | -0.004 |
| **Glutamic acid** | ZT0 | STD-VH vs. CAF-VH | 0.273 | 0.024 | -0.062 |
| | | CAF-VH vs. CAF-GSPE | 0.043 | 0.663 | 0.209 |
| | | STD-VH vs. CAF-GSPE | 0.315 | 0.009 | 0.147 |
| | ZT12 | STD-VH vs. CAF-VH | 0.098 | 0.353 | 0.130 |
| | | CAF-VH vs. CAF-GSPE | 0.179 | 0.206 | -0.164 |
| | | STD-VH vs. CAF-GSPE | 0.277 | 0.066 | -0.034 |
| **Phenylalanine** | ZT0 | STD-VH vs. CAF-VH | -0.006 | 0.853 | 0.043 |
| | | CAF-VH vs. CAF-GSPE | -0.003 | 0.917 | -0.016 |
| | | STD-VH vs. CAF-GSPE | -0.009 | 0.755 | 0.027 |
| | ZT12 | STD-VH vs. CAF-VH | -0.072 | 0.015 | -0.031 |
| | | CAF-VH vs. CAF-GSPE | 0.016 | 0.622 | 0.066 |
| | | STD-VH vs. CAF-GSPE | -0.057 | 0.119 | 0.035 |
| **Taurine** | ZT0 | STD-VH vs. CAF-VH | -0.400 | 0.266 | -0.085 |
| | | CAF-VH vs. CAF-GSPE | -0.735 | 0.043 | -0.559 |

| | | | | | |
|---|---|---|---|---|---|
| | | STD-VH vs. CAF-GSPE | -1.135 | 0.004 | -0.644 |
| | ZT12 | STD-VH vs. CAF-VH | 0.181 | 0.528 | -0.512 |
| | | CAF-VH vs. CAF-GSPE | -0.313 | 0.320 | 0.035 |
| | | STD-VH vs. CAF-GSPE | -0.132 | 0.703 | -0.477 |
| **d-Ribose** | ZT0 | STD-VH vs. CAF-VH | 0.024 | 0.004 | 0.007 |
| | | CAF-VH vs. CAF-GSPE | -0.003 | 0.814 | 0.012 |
| | | STD-VH vs. CAF-GSPE | 0.021 | 0.056 | 0.019 |
| | ZT12 | STD-VH vs. CAF-VH | -0.010 | 0.109 | 0.004 |
| | | CAF-VH vs. CAF-GSPE | 0.007 | 0.220 | 0.001 |
| | | STD-VH vs. CAF-GSPE | -0.003 | 0.564 | 0.004 |
| **d-Xylitol** | ZT0 | STD-VH vs. CAF-VH | -0.003 | 0.190 | 0.002 |
| | | CAF-VH vs. CAF-GSPE | 0.001 | 0.782 | -0.005 |
| | | STD-VH vs. CAF-GSPE | -0.003 | 0.312 | -0.003 |
| | ZT12 | STD-VH vs. CAF-VH | -0.004 | 0.032 | -0.002 |
| | | CAF-VH vs. CAF-GSPE | 0.002 | 0.253 | 0.008 |
| | | STD-VH vs. CAF-GSPE | -0.002 | 0.475 | 0.005 |
| **Glycerol-1-phosphate** | ZT0 | STD-VH vs. CAF-VH | 0.077 | 0.474 | -0.187 |
| | | CAF-VH vs. CAF-GSPE | -0.013 | 0.913 | 0.055 |
| | | STD-VH vs. CAF-GSPE | 0.065 | 0.467 | -0.132 |
| | ZT12 | STD-VH vs. CAF-VH | -0.004 | 0.964 | -0.067 |

| | | | | | |
|---|---|---|---|---|---|
| | | CAF-VH vs. CAF-GSPE | 0.118 | 0.262 | 0.130 |
| | | STD-VH vs. CAF-GSPE | 0.114 | 0.295 | 0.063 |
| **Hypoxanthine** | ZT0 | STD-VH vs. CAF-VH | -0.009 | 0.574 | 0.016 |
| | | CAF-VH vs. CAF-GSPE | 0.016 | 0.382 | 0.008 |
| | | STD-VH vs. CAF-GSPE | 0.007 | 0.677 | 0.023 |
| | ZT12 | STD-VH vs. CAF-VH | -0.075 | 0.040 | -0.058 |
| | | CAF-VH vs. CAF-GSPE | 0.035 | 0.267 | 0.016 |
| | | STD-VH vs. CAF-GSPE | -0.041 | 0.298 | -0.042 |
| **Ornithine** | ZT0 | STD-VH vs. CAF-VH | -0.008 | 0.581 | -0.039 |
| | | CAF-VH vs. CAF-GSPE | 0.008 | 0.528 | 0.004 |
| | | STD-VH vs. CAF-GSPE | -0.001 | 0.956 | -0.035 |
| | ZT12 | STD-VH vs. CAF-VH | -0.043 | 0.065 | -0.047 |
| | | CAF-VH vs. CAF-GSPE | 0.024 | 0.147 | -0.003 |
| | | STD-VH vs. CAF-GSPE | -0.019 | 0.460 | -0.050 |
| **Citric acid** | ZT0 | STD-VH vs. CAF-VH | 0.013 | 0.808 | -0.016 |
| | | CAF-VH vs. CAF-GSPE | 0.091 | 0.067 | 0.035 |
| | | STD-VH vs. CAF-GSPE | 0.105 | 0.090 | 0.019 |
| | ZT12 | STD-VH vs. CAF-VH | 0.004 | 0.932 | -0.050 |
| | | CAF-VH vs. CAF-GSPE | 0.032 | 0.612 | 0.071 |
| | | STD-VH vs. CAF-GSPE | 0.037 | 0.550 | 0.021 |

| | | | | | |
|---|---|---|---|---|---|
| **Adenine** | ZT0 | STD-VH vs. CAF-VH | -0.001 | 0.748 | 0.006 |
| | | CAF-VH vs. CAF-GSPE | -0.003 | 0.361 | -0.004 |
| | | STD-VH vs. CAF-GSPE | -0.004 | 0.273 | 0.002 |
| | ZT12 | STD-VH vs. CAF-VH | -0.007 | 0.125 | 0.005 |
| | | CAF-VH vs. CAF-GSPE | 0.003 | 0.349 | 0.004 |
| | | STD-VH vs. CAF-GSPE | -0.004 | 0.469 | 0.009 |
| **Hydroxyphenyllactic acid** | ZT0 | STD-VH vs. CAF-VH | -0.038 | 0.553 | 0.000 |
| | | CAF-VH vs. CAF-GSPE | -0.035 | 0.540 | 0.040 |
| | | STD-VH vs. CAF-GSPE | -0.073 | 0.254 | 0.039 |
| | ZT12 | STD-VH vs. CAF-VH | -0.007 | 0.923 | -0.254 |
| | | CAF-VH vs. CAF-GSPE | -0.005 | 0.949 | 0.169 |
| | | STD-VH vs. CAF-GSPE | -0.012 | 0.893 | -0.084 |
| **d-Glucose** | ZT0 | STD-VH vs. CAF-VH | 0.009 | 0.627 | 0.026 |
| | | CAF-VH vs. CAF-GSPE | 0.003 | 0.861 | -0.024 |
| | | STD-VH vs. CAF-GSPE | 0.013 | 0.488 | 0.002 |
| | ZT12 | STD-VH vs. CAF-VH | 0.008 | 0.708 | -0.026 |
| | | CAF-VH vs. CAF-GSPE | 0.026 | 0.274 | 0.036 |
| | | STD-VH vs. CAF-GSPE | 0.033 | 0.159 | 0.010 |
| **d-Mannonic acid** | ZT0 | STD-VH vs. CAF-VH | -0.004 | 0.146 | 0.004 |
| | | CAF-VH vs. CAF-GSPE | 0.000 | 0.924 | -0.002 |

| | | | | | |
|---|---|---|---|---|---|
| | | STD-VH vs. CAF-GSPE | -0.004 | 0.252 | 0.002 |
| | ZT12 | STD-VH vs. CAF-VH | -0.002 | 0.669 | 0.005 |
| | | CAF-VH vs. CAF-GSPE | 0.007 | 0.079 | -0.002 |
| | | STD-VH vs. CAF-GSPE | 0.005 | 0.355 | 0.003 |
| **d-Sorbitol** | ZT0 | STD-VH vs. CAF-VH | -0.015 | 0.630 | 0.054 |
| | | CAF-VH vs. CAF-GSPE | 0.003 | 0.928 | -0.024 |
| | | STD-VH vs. CAF-GSPE | -0.012 | 0.703 | 0.030 |
| | ZT12 | STD-VH vs. CAF-VH | -0.038 | 0.211 | -0.041 |
| | | CAF-VH vs. CAF-GSPE | 0.161 | 0.283 | 0.250 |
| | | STD-VH vs. CAF-GSPE | 0.123 | 0.410 | 0.209 |
| **d-Galactitol** | ZT0 | STD-VH vs. CAF-VH | 0.004 | 0.155 | 0.001 |
| | | CAF-VH vs. CAF-GSPE | 0.000 | 0.972 | -0.004 |
| | | STD-VH vs. CAF-GSPE | 0.004 | 0.146 | -0.002 |
| | ZT12 | STD-VH vs. CAF-VH | 0.001 | 0.627 | -0.001 |
| | | CAF-VH vs. CAF-GSPE | 0.002 | 0.299 | 0.005 |
| | | STD-VH vs. CAF-GSPE | 0.003 | 0.174 | 0.004 |
| **d-Gluconic acid** | ZT0 | STD-VH vs. CAF-VH | -0.020 | 0.230 | 0.003 |
| | | CAF-VH vs. CAF-GSPE | 0.013 | 0.399 | 0.003 |
| | | STD-VH vs. CAF-GSPE | -0.007 | 0.720 | 0.006 |
| | ZT12 | STD-VH vs. CAF-VH | -0.031 | 0.117 | -0.018 |

| | | | | | | |
|---|---|---|---|---|---|---|
| | | CAF-VH vs. CAF-GSPE | 0.007 | 0.673 | -0.009 | [illegible] |
| | | STD-VH vs. CAF-GSPE | -0.024 | 0.212 | -0.027 | [illegible] |
| **Palmitoleic acid** | ZT0 | STD-VH vs. CAF-VH | -0.012 | 0.165 | 0.001 | [illegible] |
| | | CAF-VH vs. CAF-GSPE | -0.007 | 0.360 | 0.002 | [illegible] |
| | | STD-VH vs. CAF-GSPE | -0.019 | 0.005 | 0.002 | [illegible] |
| | ZT12 | STD-VH vs. CAF-VH | -0.019 | 0.004 | -0.007 | [illegible] |
| | | CAF-VH vs. CAF-GSPE | 0.009 | 0.394 | 0.002 | [illegible] |
| | | STD-VH vs. CAF-GSPE | -0.010 | 0.390 | -0.005 | [illegible] |
| **Xanthine** | ZT0 | STD-VH vs. CAF-VH | 0.019 | 0.438 | 0.032 | [illegible] |
| | | CAF-VH vs. CAF-GSPE | 0.018 | 0.530 | 0.012 | [illegible] |
| | | STD-VH vs. CAF-GSPE | 0.037 | 0.190 | 0.043 | [illegible] |
| | ZT12 | STD-VH vs. CAF-VH | -0.050 | 0.214 | -0.018 | [illegible] |
| | | CAF-VH vs. CAF-GSPE | 0.032 | 0.382 | 0.026 | [illegible] |
| | | STD-VH vs. CAF-GSPE | -0.018 | 0.650 | 0.008 | [illegible] |
| **d-Glucuronic acid** | ZT0 | STD-VH vs. CAF-VH | 0.000 | 0.391 | 0.000 | [illegible] |
| | | CAF-VH vs. CAF-GSPE | 0.001 | 0.042 | 0.000 | [illegible] |
| | | STD-VH vs. CAF-GSPE | 0.001 | 0.250 | 0.000 | [illegible] |
| | ZT12 | STD-VH vs. CAF-VH | -0.001 | 0.024 | 0.000 | [illegible] |
| | | CAF-VH vs. CAF-GSPE | 0.001 | 0.095 | 0.000 | [illegible] |
| | | STD-VH vs. CAF-GSPE | 0.000 | 0.571 | 0.000 | [illegible] |

| | | | | | |
|---|---|---|---|---|---|
| **myo-Inositol** | ZT0 | STD-VH vs. CAF-VH | -0.100 | 0.000 | -0.034 |
| | | CAF-VH vs. CAF-GSPE | 0.003 | 0.865 | 0.001 |
| | | STD-VH vs. CAF-GSPE | -0.097 | 0.001 | -0.032 |
| | ZT12 | STD-VH vs. CAF-VH | -0.129 | 0.000 | -0.006 |
| | | CAF-VH vs. CAF-GSPE | 0.020 | 0.284 | 0.028 |
| | | STD-VH vs. CAF-GSPE | -0.109 | 0.001 | 0.022 |
| **Ribose 5--phosphate** | ZT0 | STD-VH vs. CAF-VH | 0.021 | 0.021 | -0.001 |
| | | CAF-VH vs. CAF-GSPE | -0.003 | 0.817 | 0.017 |
| | | STD-VH vs. CAF-GSPE | 0.018 | 0.128 | 0.016 |
| | ZT12 | STD-VH vs. CAF-VH | -0.013 | 0.077 | 0.004 |
| | | CAF-VH vs. CAF-GSPE | 0.006 | 0.347 | -0.004 |
| | | STD-VH vs. CAF-GSPE | -0.007 | 0.227 | 0.000 |
| **Sedoheptulose** | ZT0 | STD-VH vs. CAF-VH | -0.003 | 0.149 | -0.001 |
| | | CAF-VH vs. CAF-GSPE | -0.002 | 0.407 | -0.004 |
| | | STD-VH vs. CAF-GSPE | -0.005 | 0.055 | -0.005 |
| | ZT12 | STD-VH vs. CAF-VH | -0.005 | 0.119 | 0.000 |
| | | CAF-VH vs. CAF-GSPE | 0.001 | 0.599 | -0.002 |
| | | STD-VH vs. CAF-GSPE | -0.003 | 0.319 | -0.002 |
| **Oleic acid-iso1** | ZT0 | STD-VH vs. CAF-VH | 0.285 | 0.000 | 0.078 |
| | | CAF-VH vs. CAF-GSPE | -0.017 | 0.789 | -0.079 |

| | | | | | |
|---|---|---|---|---|---|
| | | STD-VH vs. CAF-GSPE | 0.268 | 0.000 | -0.002 |
| | ZT12 | STD-VH vs. CAF-VH | 0.209 | 0.000 | -0.004 |
| | | CAF-VH vs. CAF-GSPE | -0.036 | 0.599 | 0.030 |
| | | STD-VH vs. CAF-GSPE | 0.173 | 0.002 | 0.025 |
| **Oleic acid-iso2** | ZT0 | STD-VH vs. CAF-VH | -0.030 | 0.304 | 0.007 |
| | | CAF-VH vs. CAF-GSPE | -0.034 | 0.180 | -0.048 |
| | | STD-VH vs. CAF-GSPE | -0.064 | 0.009 | -0.041 |
| | ZT12 | STD-VH vs. CAF-VH | -0.063 | 0.018 | -0.019 |
| | | CAF-VH vs. CAF-GSPE | 0.001 | 0.954 | 0.038 |
| | | STD-VH vs. CAF-GSPE | -0.061 | 0.044 | 0.019 |
| **Stearic acid** | ZT0 | STD-VH vs. CAF-VH | 0.053 | 0.482 | -0.017 |
| | | CAF-VH vs. CAF-GSPE | 0.125 | 0.195 | 0.083 |
| | | STD-VH vs. CAF-GSPE | 0.179 | 0.117 | 0.065 |
| | ZT12 | STD-VH vs. CAF-VH | -0.022 | 0.743 | 0.012 |
| | | CAF-VH vs. CAF-GSPE | 0.115 | 0.182 | 0.038 |
| | | STD-VH vs. CAF-GSPE | 0.092 | 0.258 | 0.050 |
| **Fructose 6-phosphate** | ZT0 | STD-VH vs. CAF-VH | 0.000 | 0.992 | -0.005 |
| | | CAF-VH vs. CAF-GSPE | -0.009 | 0.017 | -0.008 |
| | | STD-VH vs. CAF-GSPE | -0.009 | 0.165 | -0.013 |
| | ZT12 | STD-VH vs. CAF-VH | -0.004 | 0.464 | -0.006 |

| | | | | | |
|---|---|---|---|---|---|
| | | CAF-VH vs. CAF-GSPE | 0.002 | 0.637 | -0.005 |
| | | STD-VH vs. CAF-GSPE | -0.002 | 0.720 | -0.011 |
| **Glucose 6-phosphate** | ZT0 | STD-VH vs. CAF-VH | 0.002 | 0.907 | -0.010 |
| | | CAF-VH vs. CAF-GSPE | -0.025 | 0.022 | -0.023 |
| | | STD-VH vs. CAF-GSPE | -0.023 | 0.177 | -0.033 |
| | ZT12 | STD-VH vs. CAF-VH | -0.008 | 0.503 | -0.013 |
| | | CAF-VH vs. CAF-GSPE | 0.004 | 0.683 | -0.012 |
| | | STD-VH vs. CAF-GSPE | -0.003 | 0.731 | -0.025 |
| **Arachidic acid** | ZT0 | STD-VH vs. CAF-VH | -0.003 | 0.619 | -0.011 |
| | | CAF-VH vs. CAF-GSPE | 0.015 | 0.108 | 0.019 |
| | | STD-VH vs. CAF-GSPE | 0.012 | 0.277 | 0.008 |
| | ZT12 | STD-VH vs. CAF-VH | -0.004 | 0.352 | 0.000 |
| | | CAF-VH vs. CAF-GSPE | 0.010 | 0.110 | 0.002 |
| | | STD-VH vs. CAF-GSPE | 0.005 | 0.391 | 0.001 |
| **Inosine** | ZT0 | STD-VH vs. CAF-VH | -0.006 | 0.927 | 0.082 |
| | | CAF-VH vs. CAF-GSPE | -0.018 | 0.806 | -0.064 |
| | | STD-VH vs. CAF-GSPE | -0.024 | 0.698 | 0.019 |
| | ZT12 | STD-VH vs. CAF-VH | -0.193 | 0.060 | -0.138 |
| | | CAF-VH vs. CAF-GSPE | 0.093 | 0.371 | 0.090 |
| | | STD-VH vs. CAF-GSPE | -0.101 | 0.388 | -0.048 |

| | | | | | |
|---|---|---|---|---|---|
| **Adenosine** | ZT0 | STD-VH vs. CAF-VH | -0.003 | 0.875 | -0.034 |
| | | CAF-VH vs. CAF-GSPE | 0.028 | 0.104 | 0.028 |
| | | STD-VH vs. CAF-GSPE | 0.026 | 0.191 | -0.006 |
| | ZT12 | STD-VH vs. CAF-VH | -0.013 | 0.733 | -0.043 |
| | | CAF-VH vs. CAF-GSPE | 0.023 | 0.413 | 0.028 |
| | | STD-VH vs. CAF-GSPE | 0.010 | 0.811 | -0.015 |
| **Xanthosine** | ZT0 | STD-VH vs. CAF-VH | 0.002 | 0.865 | 0.012 |
| | | CAF-VH vs. CAF-GSPE | -0.004 | 0.750 | -0.003 |
| | | STD-VH vs. CAF-GSPE | -0.002 | 0.880 | 0.008 |
| | ZT12 | STD-VH vs. CAF-VH | -0.009 | 0.515 | -0.020 |
| | | CAF-VH vs. CAF-GSPE | 0.009 | 0.576 | 0.025 |
| | | STD-VH vs. CAF-GSPE | -0.001 | 0.966 | 0.005 |
| **d-Maltose** | ZT0 | STD-VH vs. CAF-VH | -0.001 | 0.982 | 0.102 |
| | | CAF-VH vs. CAF-GSPE | 0.021 | 0.754 | -0.065 |
| | | STD-VH vs. CAF-GSPE | 0.020 | 0.747 | 0.037 |
| | ZT12 | STD-VH vs. CAF-VH | 0.099 | 0.340 | -0.141 |
| | | CAF-VH vs. CAF-GSPE | -0.051 | 0.615 | 0.040 |
| | | STD-VH vs. CAF-GSPE | 0.047 | 0.613 | -0.101 |
| **Lignoceric acid** | ZT0 | STD-VH vs. CAF-VH | -0.003 | 0.973 | 0.132 |
| | | CAF-VH vs. CAF-GSPE | 0.021 | 0.814 | -0.112 |

| | | | | | |
|---|---|---|---|---|---|
| | | STD-VH vs. CAF-GSPE | 0.018 | 0.821 | 0.021 |
| | ZT12 | STD-VH vs. CAF-VH | 0.160 | 0.252 | -0.176 |
| | | CAF-VH vs. CAF-GSPE | -0.073 | 0.616 | 0.075 |
| | | STD-VH vs. CAF-GSPE | 0.087 | 0.463 | -0.100 |
| **uridine 5-monophosphate** | ZT0 | STD-VH vs. CAF-VH | -0.002 | 0.945 | -0.037 |
| | | CAF-VH vs. CAF-GSPE | -0.043 | 0.087 | 0.017 |
| | | STD-VH vs. CAF-GSPE | -0.046 | 0.204 | -0.021 |
| | ZT12 | STD-VH vs. CAF-VH | 0.016 | 0.640 | -0.046 |
| | | CAF-VH vs. CAF-GSPE | -0.009 | 0.768 | -0.002 |
| | | STD-VH vs. CAF-GSPE | 0.007 | 0.837 | -0.048 |
| **inosine 5-monophosphate** | ZT0 | STD-VH vs. CAF-VH | 0.018 | 0.048 | -0.005 |
| | | CAF-VH vs. CAF-GSPE | -0.014 | 0.083 | 0.005 |
| | | STD-VH vs. CAF-GSPE | 0.004 | 0.629 | 0.000 |
| | ZT12 | STD-VH vs. CAF-VH | -0.004 | 0.630 | -0.018 |
| | | CAF-VH vs. CAF-GSPE | 0.008 | 0.330 | -0.006 |
| | | STD-VH vs. CAF-GSPE | 0.005 | 0.594 | -0.024 |
| **adenosine-5-monophosphate_adenosine-5-diphosphate_adenosine-5-triphosphate** | ZT0 | STD-VH vs. CAF-VH | 0.020 | 0.728 | -0.090 |
| | | CAF-VH vs. CAF-GSPE | -0.054 | 0.162 | 0.040 |
| | | STD-VH vs. CAF-GSPE | -0.033 | 0.573 | -0.050 |
| | ZT12 | STD-VH vs. CAF-VH | -0.005 | 0.944 | -0.100 |

| | | | | | |
|---|---|---|---|---|---|
| | | CAF-VH vs. CAF-GSPE | 0.010 | 0.874 | 0.016 |
| | | STD-VH vs. CAF-GSPE | 0.006 | 0.924 | -0.085 |
| **Cholesterol** | ZT0 | STD-VH vs. CAF-VH | -0.183 | 0.003 | 0.069 |
| | | CAF-VH vs. CAF-GSPE | 0.074 | 0.282 | -0.059 |
| | | STD-VH vs. CAF-GSPE | -0.110 | 0.055 | 0.010 |
| | ZT12 | STD-VH vs. CAF-VH | -0.263 | 0.000 | -0.049 |
| | | CAF-VH vs. CAF-GSPE | 0.105 | 0.051 | 0.041 |
| | | STD-VH vs. CAF-GSPE | -0.158 | 0.002 | -0.008 |

Rats were fed a STD or CAF diet and were treated with vehicle or GSPE at the beginning of the light phase (ZT (ZT12). d_MESOR represents the difference in MESOR values between the groups. d_amplitude represents the the groups. d_phase represents the difference in acrophase values between the groups. The $p < 0.05$ indicates s each rhythmic parameter.

**Supplementary Table 11.** Enrichment analysis of group-exclusive rhythmic liver metabolites.

| | **Rhythmic metabolites enriched pathways** | **Hits** | **Expect** | **P value** | **H** |
|---|---|---|---|---|---|

| | | | | | |
|---|---|---|---|---|---|
| ZT0 STD-VH | Mitochondrial Electron Transport Chain | 19 | 2 | 0.13 | [illegible] |
| | Urea Cycle | 29 | 2 | 0.198 | [illegible] |
| | Arginine and Proline Metabolism | 53 | 2 | 0.362 | [illegible] |
| | De Novo Triacylglycerol Biosynthesis | 9 | 1 | 0.0615 | [illegible] |
| | Malate-Aspartate Shuttle | 10 | 1 | 0.0684 | [illegible] |
| | Glycerol Phosphate Shuttle | 11 | 1 | 0.0752 | |
| | Cardiolipin Biosynthesis | 11 | 1 | 0.0752 | |
| | Spermidine and Spermine Biosynthesis | 18 | 1 | 0.123 | |
| | Glutathione Metabolism | 21 | 1 | 0.144 | |
| | Glycerolipid Metabolism | 25 | 1 | 0.171 | |
| | Phenylalanine and Tyrosine Metabolism | 28 | 1 | 0.191 | |
| | Phospholipid Biosynthesis | 29 | 1 | 0.198 | |
| | Citric Acid Cycle | 32 | 1 | 0.219 | |
| | Aspartate Metabolism | 35 | 1 | 0.239 | |
| | Gluconeogenesis | 35 | 1 | 0.239 | |
| | Warburg Effect | 58 | 1 | 0.396 | |
| | Glycine and Serine Metabolism | 59 | 1 | 0.403 | |
| | Tyrosine Metabolism | 72 | 1 | 0.492 | |
| | Purine Metabolism | 74 | 1 | 0.506 | |
| | Galactose Metabolism | 38 | 2 | 0.186 | [illegible] |

| | | | | | |
|---|---|---|---|---|---|
| **ZT0-CAF-VH** | Lactose Degradation | 9 | 1 | 0.0439 | [illegible] |
| | Glucose-Alanine Cycle | 13 | 1 | 0.0635 | |
| | Spermidine and Spermine Biosynthesis | 18 | 1 | 0.0879 | |
| | Lactose Synthesis | 20 | 1 | 0.0977 | [illegible] |
| | Betaine Metabolism | 21 | 1 | 0.103 | [illegible] |
| | Transfer of Acetyl Groups into Mitochondria | 22 | 1 | 0.107 | |
| | Glycolysis | 25 | 1 | 0.122 | |
| | Plasmalogen Synthesis | 26 | 1 | 0.127 | |
| | Mitochondrial Beta-Oxidation of Long Chain Saturated Fatty Acids | 28 | 1 | 0.137 | |
| | Fructose and Mannose Degradation | 32 | 1 | 0.156 | |
| | Gluconeogenesis | 35 | 1 | 0.171 | |
| | Sphingolipid Metabolism | 40 | 1 | 0.195 | |
| | Propanoate Metabolism | 42 | 1 | 0.205 | |
| | Methionine Metabolism | 43 | 1 | 0.21 | |
| | Warburg Effect | 58 | 1 | 0.283 | |
| | Glycine and Serine Metabolism | 59 | 1 | 0.288 | |
| **ZT0-CAF-GSPE** | Malate-Aspartate Shuttle | 10 | 1 | 0.0195 | [illegible] |
| | Glucose-Alanine Cycle | 13 | 1 | 0.0254 | [illegible] |
| | Alanine Metabolism | 17 | 1 | 0.0332 | [illegible] |
| | Glutathione Metabolism | 21 | 1 | 0.041 | [illegible] |

| | | | | | |
|---|---|---|---|---|---|
| | Cysteine Metabolism | 26 | 1 | 0.0508 | [illegible] |
| | Phenylalanine and Tyrosine Metabolism | 28 | 1 | 0.0547 | |
| | Folate Metabolism | 29 | 1 | 0.0566 | [illegible] |
| | Urea Cycle | 29 | 1 | 0.0566 | [illegible] |
| | Lysine Degradation | 30 | 1 | 0.0586 | [illegible] |
| | Ammonia Recycling | 32 | 1 | 0.0625 | [illegible] |
| | Amino Sugar Metabolism | 33 | 1 | 0.0645 | [illegible] |
| | Beta-Alanine Metabolism | 34 | 1 | 0.0664 | [illegible] |
| | Aspartate Metabolism | 35 | 1 | 0.0684 | [illegible] |
| | Nicotinate and Nicotinamide Metabolism | 37 | 1 | 0.0723 | |
| | Propanoate Metabolism | 42 | 1 | 0.082 | [illegible] |
| | Histidine Metabolism | 43 | 1 | 0.084 | [illegible] |
| | Glutamate Metabolism | 49 | 1 | 0.0957 | [illegible] |
| | Arginine and Proline Metabolism | 53 | 1 | 0.104 | |
| | Warburg Effect | 58 | 1 | 0.113 | |
| | Glycine and Serine Metabolism | 59 | 1 | 0.115 | |
| | Valine, Leucine and Isoleucine Degradation | 60 | 1 | 0.117 | |
| | Tryptophan Metabolism | 60 | 1 | 0.117 | |
| | Arachidonic Acid Metabolism | 69 | 1 | 0.135 | |
| | Tyrosine Metabolism | 72 | 1 | 0.141 | |

| | | | | | |
|---|---|---|---|---|---|
| | Purine Metabolism | 74 | 1 | 0.145 | |
| **ZT12-STD-VH** | Thiamine Metabolism | 9 | 3 | 0.0703 | [illegible] |
| | Alanine Metabolism | 17 | 3 | 0.133 | [illegible] |
| | Riboflavin Metabolism | 20 | 3 | 0.156 | [illegible] |
| | Glutathione Metabolism | 21 | 3 | 0.164 | [illegible] |
| | Pantothenate and CoA Biosynthesis | 21 | 3 | 0.164 | [illegible] |
| | Glycine and Serine Metabolism | 59 | 4 | 0.461 | [illegible] |
| | Cysteine Metabolism | 26 | 3 | 0.203 | [illegible] |
| | Selenoamino Acid Metabolism | 28 | 3 | 0.219 | [illegible] |
| | Pentose Phosphate Pathway | 29 | 3 | 0.227 | [illegible] |
| | Urea Cycle | 29 | 3 | 0.227 | [illegible] |
| | Ammonia Recycling | 32 | 3 | 0.25 | [illegible] |
| | Fructose and Mannose Degradation | 32 | 3 | 0.25 | [illegible] |
| | Phenylacetate Metabolism | 9 | 2 | 0.0703 | [illegible] |
| | Lactose Degradation | 9 | 2 | 0.0703 | [illegible] |
| | Nicotinate and Nicotinamide Metabolism | 37 | 3 | 0.289 | [illegible] |
| | Galactose Metabolism | 38 | 3 | 0.297 | [illegible] |
| | Trehalose Degradation | 11 | 2 | 0.0859 | [illegible] |
| | Propanoate Metabolism | 42 | 3 | 0.328 | [illegible] |
| | Methionine Metabolism | 43 | 3 | 0.336 | [illegible] |

| | | | | | |
|---|---|---|---|---|---|
| | Histidine Metabolism | 43 | 3 | 0.336 | [illegible] |
| | Phosphatidylethanolamine Biosynthesis | 12 | 2 | 0.0938 | [illegible] |
| | Pyruvate Metabolism | 48 | 3 | 0.375 | [illegible] |
| | Phosphatidylcholine Biosynthesis | 14 | 2 | 0.109 | [illegible] |
| | Glutamate Metabolism | 49 | 3 | 0.383 | [illegible] |
| | Arginine and Proline Metabolism | 53 | 3 | 0.414 | [illegible] |
| | Beta Oxidation of Very Long Chain Fatty Acids | 17 | 2 | 0.133 | [illegible] |
| | Phosphatidylinositol Phosphate Metabolism | 17 | 2 | 0.133 | [illegible] |
| | Spermidine and Spermine Biosynthesis | 18 | 2 | 0.141 | [illegible] |
| | Butyrate Metabolism | 19 | 2 | 0.148 | [illegible] |
| | Mitochondrial Electron Transport Chain | 19 | 2 | 0.148 | [illegible] |
| | Ethanol Degradation | 19 | 2 | 0.148 | [illegible] |
| | Nucleotide Sugars Metabolism | 20 | 2 | 0.156 | [illegible] |
| | Lactose Synthesis | 20 | 2 | 0.156 | [illegible] |
| | Threonine and 2-Oxobutanoate Degradation | 20 | 2 | 0.156 | [illegible] |
| | Betaine Metabolism | 21 | 2 | 0.164 | [illegible] |
| | Bile Acid Biosynthesis | 65 | 3 | 0.508 | [illegible] |
| | Sulfate/Sulfite Metabolism | 22 | 2 | 0.172 | [illegible] |
| | Transfer of Acetyl Groups into Mitochondria | 22 | 2 | 0.172 | [illegible] |
| | Glycerolipid Metabolism | 25 | 2 | 0.195 | [illegible] |

| | | | | | |
|---|---|---|---|---|---|
| | Glycolysis | 25 | 2 | 0.195 | [illegible] |
| | Purine Metabolism | 74 | 3 | 0.578 | [illegible] |
| | Oxidation of Branched Chain Fatty Acids | 26 | 2 | 0.203 | [illegible] |
| | Phytanic Acid Peroxisomal Oxidation | 26 | 2 | 0.203 | [illegible] |
| | Inositol Phosphate Metabolism | 26 | 2 | 0.203 | [illegible] |
| | Mitochondrial Beta-Oxidation of Short Chain Saturated Fatty Acids | 27 | 2 | 0.211 | [illegible] |
| | Mitochondrial Beta-Oxidation of Medium Chain Saturated Fatty Acids | 27 | 2 | 0.211 | [illegible] |
| | Phenylalanine and Tyrosine Metabolism | 28 | 2 | 0.219 | [illegible] |
| | Mitochondrial Beta-Oxidation of Long Chain Saturated Fatty Acids | 28 | 2 | 0.219 | [illegible] |
| | Folate Metabolism | 29 | 2 | 0.227 | [illegible] |
| | Starch and Sucrose Metabolism | 31 | 2 | 0.242 | [illegible] |
| | Citric Acid Cycle | 32 | 2 | 0.25 | [illegible] |
| | Inositol Metabolism | 33 | 2 | 0.258 | [illegible] |
| | Amino Sugar Metabolism | 33 | 2 | 0.258 | [illegible] |
| | Aspartate Metabolism | 35 | 2 | 0.273 | [illegible] |
| | Gluconeogenesis | 35 | 2 | 0.273 | [illegible] |
| | Sphingolipid Metabolism | 40 | 2 | 0.312 | [illegible] |
| | Fatty acid Metabolism | 43 | 2 | 0.336 | [illegible] |
| | Steroid Biosynthesis | 48 | 2 | 0.375 | [illegible] |
| | Biotin Metabolism | 8 | 1 | 0.0625 | [illegible] |

| | | | | | |
|---|---|---|---|---|---|
| | Warburg Effect | 58 | 2 | 0.453 | [illegible] |
| | Pyrimidine Metabolism | 59 | 2 | 0.461 | [illegible] |
| | Valine, Leucine and Isoleucine Degradation | 60 | 2 | 0.469 | [illegible] |
| | Taurine and Hypotaurine Metabolism | 12 | 1 | 0.0938 | [illegible] |
| | Tryptophan Metabolism | 60 | 1 | 0.469 | [illegible] |
| **ZT12-CAF-VH** | Glycine and Serine Metabolism | 59 | 3 | 0.23 | [illegible] |
| | Glucose-Alanine Cycle | 13 | 2 | 0.0508 | [illegible] |
| | Alanine Metabolism | 17 | 2 | 0.0664 | [illegible] |
| | Glutathione Metabolism | 21 | 2 | 0.082 | [illegible] |
| | Urea Cycle | 29 | 2 | 0.113 | [illegible] |
| | Glutamate Metabolism | 49 | 2 | 0.191 | [illegible] |
| | Warburg Effect | 58 | 2 | 0.227 | [illegible] |
| | Tryptophan Metabolism | 60 | 2 | 0.234 | [illegible] |
| | Malate-Aspartate Shuttle | 10 | 1 | 0.0391 | [illegible] |
| | Cysteine Metabolism | 26 | 1 | 0.102 | [illegible] |
| | Phenylalanine and Tyrosine Metabolism | 28 | 1 | 0.109 | [illegible] |
| | Selenoamino Acid Metabolism | 28 | 1 | 0.109 | [illegible] |
| | Folate Metabolism | 29 | 1 | 0.113 | [illegible] |
| | Lysine Degradation | 30 | 1 | 0.117 | [illegible] |
| | Ammonia Recycling | 32 | 1 | 0.125 | [illegible] |

| | | | | | |
|---|---|---|---|---|---|
| | Amino Sugar Metabolism | 33 | 1 | 0.129 | |
| | Beta-Alanine Metabolism | 34 | 1 | 0.133 | |
| | Aspartate Metabolism | 35 | 1 | 0.137 | |
| | Gluconeogenesis | 35 | 1 | 0.137 | |
| | Nicotinate and Nicotinamide Metabolism | 37 | 1 | 0.145 | |
| | Propanoate Metabolism | 42 | 1 | 0.164 | |
| | Methionine Metabolism | 43 | 1 | 0.168 | |
| | Histidine Metabolism | 43 | 1 | 0.168 | |
| | Pyruvate Metabolism | 48 | 1 | 0.188 | |
| | Arginine and Proline Metabolism | 53 | 1 | 0.207 | |
| | Valine, Leucine and Isoleucine Degradation | 60 | 1 | 0.234 | |
| | Arachidonic Acid Metabolism | 69 | 1 | 0.27 | |
| | Tyrosine Metabolism | 72 | 1 | 0.281 | |
| | Purine Metabolism | 74 | 1 | 0.289 | |
| **ZT12-CAF-GSPE** | Pentose Phosphate Pathway | 29 | 1 | 0.085 | [illegible] |
| | Valine, Leucine and Isoleucine Degradation | 60 | 1 | 0.176 | |

Group-exclusive rhythmic metabolites enriched pathways of rats that were fed a STD or CAF diet and were treat
of the light phase (ZT0) or at the beginning of the dark phase (ZT12). Metabolic pathways enrichment analysis wa
5.0, URL: http://www.metaboanalyst.ca) using the metabolites that were rhythmic exclusively in each group.
